# pH selects latent allosteric states in insulin glargine assemblies

**DOI:** 10.64898/2026.04.07.716775

**Authors:** Esra Ayan, Madan Kumar Shankar, Elek Telek, Jungmin Kang, Krisztian Fintor, Toshinori Yabuuchi, Makina Yabashi, Takehiko Tosha

## Abstract

How protein assemblies couple environmental triggers to controlled assembly and disassembly remains a central question in structural biology and biophysics. Insulin glargine provides a clinically important model for pH-triggered assembly-disassembly coupling, yet the structural pathway linking isoelectric depot to delayed release has remained unresolved. Here we combine ambient-temperature serial femtosecond crystallography, solution biophysics, and multiscale network analyses to define the pH-dependent conformational landscape of hexameric glargine across pH 8.4, 7.3, 6.4, and 5.1. We resolve full hexameric glargine structures and directly visualize pH-coupled lattice transition from P12_1_1 (near-neutral) to R3:H (acidic), accompanied by redistribution from compact phenolic R^f^_6_-state assemblies to more plastic yet structurally coherent TR^f^/T_3_R^f^_3_ states. These results suggest that isoelectric precipitation and delayed dissociation may be linked through structurally organized molten-like intermediate states and provide a testable hypothesis for how allosteric protein assemblies respond to environmental changes through redistribution of latent collective modes and suggest a broader mechanism by which allosteric protein assemblies encode environmental responsiveness through redistribution of latent collective modes.

**Significance:** This work provides a state-resolved structural framework for investigating how an environmentally responsive protein assembly may couple pH sensing to controlled release. Using insulin glargine as a clinically important pH-sensitive model, our findings suggest that precipitation-linked delayed action may involve redistribution through structurally organized intermediate hexamer states that remain connected to the intrinsic allosteric landscape. Beyond basal insulin pharmacology, these findings position precipitation-coupled allostery as a general design principle for protein assemblies whose function depends on environmentally gated stability, release, and responsiveness.

## 1. Introduction

Insulin naturally forms zinc-coordinated hexamers that exhibit classical allostery, interconverting between three conformational states: T_6_, T_3_R_3_, and R_6_ [1], [2] (**Fig. S1a**). These designations (tense *T* and relaxed *R*) were adopted by analogy to hemoglobin once distinct hexamer structures were elucidated in the 1980s [3], [4]. The R-state involves a significant conformational rearrangement of the insulin monomer: residues B1–B8, which are extended in the T-state, refold into an alpha-helix (extending B1–B19) in the R-state [5], [6]. This transition moves F1_B_ by ∼30 Å and creates an apolar ligand-binding pocket coordinated by the H10_B_-bound Zn^2+^ [2], [5]. Phenolic derivatives (e.g., phenol, m-cresol) serve as positive homotropic effectors that bind in this pocket and stabilize the R-conformation [7], [8]. Meanwhile, anions (eg., Cl⁻ or SCN⁻) act as heterotropic effectors by binding to the Zn^2+^, often promoting intermediate T_3_R_3_ states [9]. Through these allosteric ligand interactions, the phenol-bound R₆ insulin hexamer is thought to be substantially more stable than the T₆ form [2]. Accordingly, phenolic additives in insulin formulations shift the conformational equilibrium toward the R₆ state, stabilizing the quaternary structure and protecting insulin from degradation [8], [10], [11]. This allosteric hexamer system has become a valuable model for protein allostery and ligand-dependent stabilization, even though insulin’s *in vivo* functional form is the monomer [3], [12]. Notably, patients benefit from hexamer-based insulin pharmaceuticals because the hexamer serves as a depot that delays monomer release [13].

Insulin glargine is a biosynthetic insulin analog designed to provide prolonged glycemic control through pH-dependent changes in solubility and depot formation. It differs from human insulin by a G21_A_ substitution and the addition of two arginine residues at the B-chain C-terminus (R31_B_ and R32_B_) [14], [15]. These modifications raise the isoelectric point (pI) of insulin to near-neutral pH, making glargine soluble in the acidic formulation (pH 4 in pen) but poorly soluble at physiological pH (in the human body) [16]. Upon subcutaneous injection, the solution is neutralized to pH ∼7.4, causing glargine to precipitate into a subdermal depot from which insulin slowly dissociates over ∼24 hours [17]. This mechanism underpins glargine’s basal, peakless pharmacokinetic profile. However, the structural basis for glargine precipitation followed by sustained release has remained speculative; indeed, as previously noted, “the actual shape of an insulin glargine depot is unknown” [18], [19]. A longstanding question is how glargine’s unique sequence influences the assembly or conformational state of the insulin hexamer at neutral pH. Since glargine is formulated at pH 4 with phenolic preservatives and zinc, upon injection, the rise to physiological pH triggers not only isoelectric precipitation but also a pronounced allosteric shift in its hexameric structure [5]. We hypothesized that glargine precipitation following injection may be accompanied by changes in phenolic ligand binding and the allosteric equilibrium of the hexamer. Yet prior X-ray structures of insulin glargine were determined at cryogenic temperature and in non-physiological conditions (PDB IDs: 4IYD, 5VIZ, 8WU0); they captured only part of the molecule (often a monomer in the asymmetric unit). Thus, key aspects, such as how glargine adopts between T/R conformations in its precipitated form, have not been visualized. Detailed structural characterization of glargine under relevant pH conditions is needed to fill this gap [20]. Modern structural biology offers new tools to examine proteins under more native-like conditions [20]. In particular, serial femtosecond X-ray crystallography (SFX) using X-ray free-electron lasers enables structure determination at near-physiological (ambient) temperature with extremely brief X-ray pulses that outrun radiation damage [21], [22]. Unlike conventional cryogenic crystallography, SFX can reveal biologically relevant conformational ensembles that might be hidden or “quenched” at cryo temperatures [23], [24], [25]. Cryocooling, while immensely successful in reducing radiation damage, can trap proteins in a single conformational sub-state and mask dynamic or alternate conformations [25], [26]. Indeed, a growing body of evidence shows that the vast archive of cryo-structures may provide an incomplete picture of protein allostery [26], [27]. By contrast, structures at the ambient temperature tend to exhibit higher atomic displacement (B-factors) and sometimes multiple conformations, offering insight into flexibility and allosteric transition pathways [28], [29]. For allosteric systems like the insulin hexamer, this is especially pertinent: subtle shifts in conformational ensembles could underlie the cooperative ligand effects [30], [31], [32]. Moreover, glargine’s function involves the formation of microcrystals (depot) in the human body, conditions that cryo-structures do not visualize [28]. SFX is uniquely suited here [33], [34]: by collecting data from thousands of microcrystals at 298 K, we can capture glargine’s native structure in extended pH intervals free from cryo-artifacts or cumulative X-ray damage.

In this work, we combine ambient temperature SFX with solution biophysical assays and computational modeling to investigate pH-dependent structural changes potentially relevant to insulin glargine’s prolonged action. We obtained (i) crystallographic data for glargine over a pH range from 8.4 to 5.1, yielding a series of structural “snapshots” spanning conditions from its precipitated formulation to the loosely coupled arrangement at the ambient temperature. We further probed (ii) pH-dependent structural transitions *in-solution* via Raman spectroscopy, fluorescence measurement, and differential scanning calorimetry, to ensure that the crystalline states correspond to solution behavior. Finally, to connect structural changes with dynamics, we employed (iii) *in-silico* analyses (the Gaussian network model (GNM) and anisotropic network model (ANM)) [35], [36] on the determined structures. By integrating these approaches, we present a comprehensive, multi-resolution picture of how pH, homotropic phenolic ligands, and heterotropic ions control an allosteric switch in insulin glargine. The results reveal, at the molecular level under ambient-temperature conditions, how glargine undergoes an allosteric T↔R transition coupled to phenol binding, and how this transition may be related to precipitation-associated structural reorganization. We also highlight key differences between the SFX structures and prior 16 cryogenic (insulin, not glargine) structures from PDB, underscoring the value of SFX for capturing functionally essential protein motions. Our findings provide a structural framework for investigating whether pH-dependent R↔T transitions contribute to glargine depot formation and subsequent dissociation. We therefore asked whether pH-dependent delayed release in glargine reflects simple precipitation followed by passive dissociation or instead proceeds through structurally organized intermediate states that remain coupled to the insulin hexamer’s intrinsic allosteric landscape.

## 2. Results

### 2.1 pH induces in-crystallo phase transition driven by helical ‘unpeeling’ motion

We determined hexameric insulin glargine structures from microcrystals at four pH conditions (8.4, 7.3, 6.4, and 5.1), at resolutions of 1.84, 1.97, 2.07, and 2.30 Å, respectively, all measured at 298 K. For SFX data collection, an HVC injector was used, which is compatible with the diffraction chamber of the Diverse Application Platform for Hard X-ray Diffraction in SACLA (DAPHNIS) (**Fig. 1a**) [37]. Hexameric glargine formed small crystals (∼5 μm; **Fig. S1**) that did not produce detectable diffraction using conventional in-house X-ray sources (**Fig. S2**). Nine buffer conditions spanning pH 8.4 to 3.0 were applied to these microcrystals; at lower pH values (approximately 4.7–3.0), the crystals visibly disappeared (**Fig. S1c**). This loss of crystal integrity and the concomitant rise in background noise observed as the pH was lowered from 8.4 to 3.0 parallels the physical disintegration process previously reported by Bentley et al. (1978) during the transformation of rhombohedral insulin crystals [38]. In that study, forcing the conversion of 4-Zn crystals to the 2-Zn form via environmental modification—specifically, exposure to distilled water or 2-Zn mother liquor—generated lattice stress that resulted in cracking and ultimate disintegration. Analogously, the cessation of diffraction below pH 5.1 and the gradual disappearance of cubic crystals in our pH screening suggest that pH-induced structural or electrostatic perturbations exceeded the tolerance limits of the crystal lattice, thereby triggering a mechanism of lattice disorder and structural collapse similar to that described by Bentley [38]. Of the seven sampled pH conditions (**Fig. S3**), only four (pH 8.4, 7.3, 6.4, and 5.1) were replicated three times with well-defined diffraction patterns using HVC-mediated SFX at SACLA (**Figs. S4 and S5**). Data-collection statistics reveal higher hit acceptance and indexing efficiency at physiological pHs (8.4–7.3), with a pronounced reduction at more acidic pHs (6.4–5.1) (**Table S1**).

**Figure 1.**
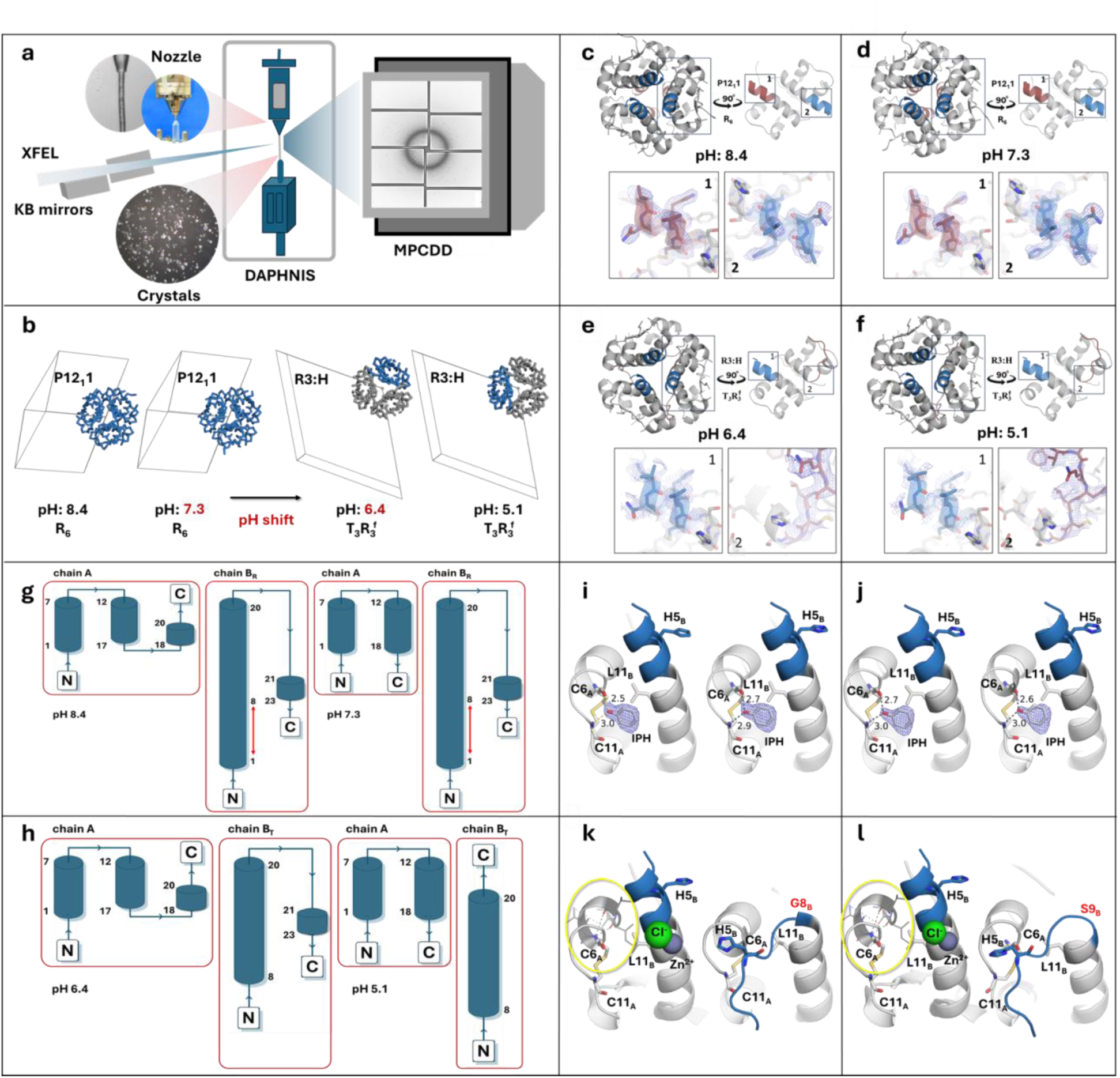
pH-driven conformational allostery and crystallographic lattice reorganization of hexameric insulin glargine resolved by SFX. a, Schematic of the SFX experimental setup at SACLA, illustrating microcrystal injection via the HVC injector into the DAPHNIS chamber, XFEL illumination, and diffraction recording on the MPCC Detector. b, pH-dependent crystal packing transition showing a monoclinic lattice (space group P12_1_1, *R_f_* hexamer) at pH 8.4 and 7.3, and a rhombohedral lattice (space group R3:H, *TR*_f_ assembly) at pH 6.4 and 5.1, highlighting the pH-induced shift in asymmetric unit content and symmetry. c–f, Overall hexamer structures at pH 8.4, 7.3, 6.4, and 5.1, respectively, with corresponding electron-density maps emphasizing conformational transitions in the B-chain N-terminus (residues 1–8). g-h, Secondary-structure topology diagrams for chains A and B across pH conditions, revealing a progressiv e, chain-asymmetric shortening of the B chain at pHs 6.4 and 5.1 as well as a cooperative reorganization at pH 5.1 in which the topological distinction of B collapses into a unified architecture. i-l, Close-up views of the B-chain N-terminal region highlighting pH-dependent rearrangements of key residues (C6_A_, C11_A_, H5_B_, L11_B_ –those are related to IPH), coordination geometry (Zn^2+^, CI^−^), and local interactions around the zinc-binding site, including altered side-chain orientations (1-8_B_). For panels c–f and i–j, the 2mFₒ–DFc electron-density maps are contoured at 1.0σ and carved within 1.6 Å of the displayed atomic models

Strikingly, the resolved structures segregate into two distinct lattice packings of the hexamer glargine corresponding to high- and low-pH values (**Fig. 1b**). Crystals obtained at near-physiological pHs adopt a monoclinic lattice (space group P12_1_1), with highly consistent unit-cell parameters across the pH 8.4 and 7.3 datasets (**Figs. S6a–d**). In contrast, within ∼10 min of acidification the same monoclinic crystals convert in crystallo to a rhombohedral lattice (space group R3 in the hexagonal setting, R3:H) (**Figs. S6e–f**), isomorphous with previously reported canonical 4Zn insulin structures [39], [40]. The monoclinic lattice contains one hexamer per asymmetric unit (ASU) (**Figs. S6a–d**), whereas the rhombohedral lattice contains a dimer per ASU (**Figs. S6e–f**). Consistently, 3D Ramachandran maps visualize this transition as a shift from rigidly locked backbone conformations to a dispersed, multi-modal landscape indicative of enhanced plasticity (**Fig. S6**). This change in crystal packing is closely associated with conformational allostery involving residues 1–8 of the B chain in each monomer (**Figs. 1c–f**). At pH 8.4 and 7.3, B-chain residues 1–8 adopt a frayed-relaxed/helical conformation (hereafter referred to as *R_f_* state) that is continuous with the remaining B-chain helices (B1-B19) (**Figs. 1c–d**). By contrast, at pH 6.4 and 5.1, these residues assume a tense conformation (hereafter referred to as *T* state), F1_B_ moves by ∼30 Å (**Figs. 1e–f**). These differences are also evident in the secondary-structure topology of the monomers (**Figs. 1g–h; Fig. S7**), showing a progressive, chain-asymmetric shortening that predominantly affects the B chain at pH 6.4 and 5.1 (**Figs. 1h**), and at pH 5.1 a further loss of helical definition in which the topological distinction in the B chain has collapsed (**Figs. 1h; Fig. S7d**); difference electron-density maps (**Fig. S8**) show that pH 5.1 represents a more disordered version of the pH 6.4 T₃R₃ intermediate rather than a new state. The apparent distinction is also captured by the diffraction data, manifested by space-group–dependent differences in scattering profiles and *d*-spacing distributions across the pH conditions (3.9–1.9 Å at pH 8.4, progressively restricted to 3.9–2.7 Å at pH 5.1) (**Fig. S5**), consistent with allostery-driven changes in unit-cell symmetry and interplanar spacings.

At pH 8.4 and 7.3, glargine hexamers are in the classic *R^f^*_6_ state – all six protomers adopt the *R_f_* conformation with their B1–B19 helices intact (**Fig. S1b** and **Fig. S8a-b**), and electron density indicates six phenol molecules bound per hexamer (one in each subunit’s hydrophobic pocket) (**Fig. 1i-j; Fig. S9a-b**). This aligns well with the *R^f^*_6_ insulin structures in the Protein Data Bank (2WS7 [41], 4P65 [42], 5EMS [43], 5HRQ [44], 5UDP [45], 6TYH [46]) (**Fig. S10**). However, in our near-physiological-temperature SFX data at pH 8.4 and 7.3, elevated B-factors (∼40–45 Å²) and perturbed hydrogen-bond networks in the allosteric region, including the S9_B_-E13_B_ scaffold and Zn²⁺ stabilized contacts involving F1_B_ and A14_B_ (**Fig. S9e-f**), contrast sharply but reasonably with the artificially rigidified scaffolds typical of cryogenic models (17Å² to 33Å²) (**Fig. S10**) [47]. Rather than static disorder, we attribute this plasticity to physiological thermal ‘breathing’ often masked by cryo-arrest [48]; specifically, the dynamic destabilization of the central E13_B_ cluster, accompanied by substantial shifts in its predicted pKa values (**Table 1**), likely modulates the local electrostatic repulsion [47], [49], thereby unlocking the entropic freedom required for the high long-range allosteric cooperativity evidenced in our cross-correlation analyses (**Fig. S10**).

**Table 1.** PROPKA-predicted pKa of titratable residues in the four SFX structures at different pH conditions.

| pH Value | Residue ID | Predicted pKa | Model pKa | Predicted Protonation State | pH Value | Residue ID | Predicted pKa | Model pKa | Predicted Protonation State |
|---|---|---|---|---|---|---|---|---|---|
| pH 8.4 | GLU 13B | 11.28 | 4.50 | Protonated (neutral) | pH 7.3 | GLU 13B | 5.70 | 4.50 | Deprot. (-) |
|  | GLU 13D | 14.57 | 4.50 |  |  | GLU 13D | 8.36 | 4.50 | Prot. (neutral) |
|  | HIS 5B | 4.15 | 6.50 | Unprotonated (neutral) |  | HIS 5B | 4.11 | 6.50 | Unprotonated (neutral) |
|  | HIS 10B | 2.22 | 6.50 |  |  | HIS 10B | 2.11 | 6.50 |  |
|  | HIS 5D | 4.29 | 6.50 |  |  | HIS 5D | 4.36 | 6.50 |  |
|  | HIS 10D | 0.76 | 6.50 |  |  | HIS 10D | 0.75 | 6.50 |  |
|  | CYS 6A | 99.99 | 9.00 | S—S |  | CYS 6A | 99.99 | 9.00 | S—S |
|  | CYS 7A | 99.99 | 9.00 |  |  | CYS 7A | 99.99 | 9.00 |  |
|  | CYS 11A | 99.99 | 9.00 |  |  | CYS 11A | 99.99 | 9.00 |  |
|  | CYS 20A | 99.99 | 9.00 |  |  | CYS 20A | 99.99 | 9.00 |  |
|  | CYS 7B | 99.99 | 9.00 |  |  | CYS 7B | 99.99 | 9.00 |  |
|  | CYS 19B | 99.99 | 9.00 |  |  | CYS 19B | 99.99 | 9.00 |  |
|  | LYS 29F | 10.36 | 10.50 | Protonated (+) |  | LYS 29F | 10.36 | 10.50 | Protonated (+) |
|  | LYS 29L | 10.50 | 10.50 |  |  | LYS 29L | 10.50 | 10.50 |  |
|  | ARG 22B | 11.94 | 12.50 |  |  | ARG 22B | 11.94 | 12.50 |  |
|  | ARG 22D | 11.87 | 12.50 |  |  | ARG 22D | 11.87 | 12.50 |  |
|  | **Residue ID** | **Predicted pKa** | **Model pKa** | **Predicted Protonation State** |  | **Residue ID** | **Predicted pKa** | **Model pKa** | **Predicted Protonation State** |
| pH 6.4 | GLU 13B | 4.42 | 4.50 | Deprotonated (-) | pH 5.1 | GLU 13B | 4.14 | 4.50 | Deprotonated (-) |
|  | GLU 13D | 4.74 | 4.50 |  |  | GLU 13D | 3.91 | 4.50 |  |
|  | HIS 5B | 6.50 | 6.50 | Protonated (+) |  | HIS 5B | 6.50 | 6.50 | Protonated (+) |
|  | HIS 10B | 6.50 | 6.50 |  |  | HIS 10B | 6.50 | 6.50 |  |
|  | HIS 5D | 6.32 | 6.50 | Unprot. (neutral) |  | HIS 5D | 6.43 | 6.50 |  |
|  | HIS 10D | 6.62 | 6.50 | Protonated (+) |  | HIS 10D | 6.50 | 6.50 |  |
|  | CYS 6A | 99.99 | 9.00 | S—S |  | CYS 6A | 99.99 | 9.00 | S—S |
|  | CYS 7A | 99.99 | 9.00 |  |  | CYS 7A | 99.99 | 9.00 |  |
|  | CYS 11A | 99.99 | 9.00 |  |  | CYS 11A | 99.99 | 9.00 |  |
|  | CYS 20A | 99.99 | 9.00 |  |  | CYS 20A | 99.99 | 9.00 |  |
|  | CYS 7B | 99.99 | 9.00 |  |  | CYS 7B | 99.99 | 9.00 |  |
|  | CYS 19B | 99.99 | 9.00 |  |  | CYS 19B | 99.99 | 9.00 |  |
|  | LYS 29F | N/A |  |  |  | LYS 29D | 10.50 | 10.50 | Protonated (+) |
|  | LYS 29L |  |  |  |  | LYS 29F/L | N/A |  |  |
|  | ARG 22B | 11.87 | 12.50 | Protonated (+) |  | ARG 22B | 11.73 | 12.50 | Protonated (+) |
|  | ARG 22D | 11.94 | 12.50 |  |  | ARG 22D | 12.36 | 12.50 |  |

Acidification of the glargine crystals from pH 8.4 and 7.3 to pH 6.4 and 5.1 yields a *T_3_R^f^_3_* hexamer, comprising three T-state and three Rf-state protomers (**Fig. S1b**), analogous to the classical 4-Zn insulin model, lacking phenol highlighted in yellow circle in **Figure 1k-l** (**Fig. S9c-d**).

Whereas the phenolic ligand-binding pockets of T-state protomers are disrupted by the extended B1–B8 segment (**Fig. S8c-d; Fig. S9c-d-S9h**), the three Rf-state protomers retain their characteristic topology, potentially stabilized by axial Cl⁻ coordination to Zn²⁺ (**Fig. 1k-l; Fig. S9c-d, h; Fig. S11a_vii-viii_**), preserving the conventional *R^f^* topology observed in the high-pH (8.4 and 7.3) structures (**Fig. 1i-j**). This is in line with previous cryogenic crystallographic data (PDB IDs: 1J73 [50], 1QJ0 [51], 2QIU [52], 2R34-35 [53]), in which the *R^f^_3_* -conformation was similarly maintained by the coordination of either Cl^−^ anions or structured water molecules to the Zn^2+^ metal center (**Fig. S11**) [1], [54], [55]. To further validate these pH-dependent structural changes independently of the refined low-pH models, we calculated real-space difference electron-density maps using *matchmaps*, accounting for the distinct crystal forms and merohedral twinning of the acidic datasets (*Methods*; **Fig. S8e–g; Fig. 2**). The maps localized the major changes to the Zn²⁺ coordination sites and the B-chain N-terminal region, particularly F1B and H10B, while the hydrophobic core remained largely unchanged. Refined Zn²⁺ occupancies decreased from 0.87–0.88 at near-neutral pH to 0.33–0.63 at acidic pH, indicating partial, site-dependent depletion rather than complete release. Additional omit maps supported the T-state B1–B8 conformation and revealed residual phenolic-ligand density at pH 6.4 but no detectable density at pH 5.1. Together, these results suggest that the principal structural reorganization occurs between pH 7.3 and 6.4, with pH 5.1 representing a more disordered T₃Rf₃ intermediate rather than a distinct conformational state.

**Figure 2.**
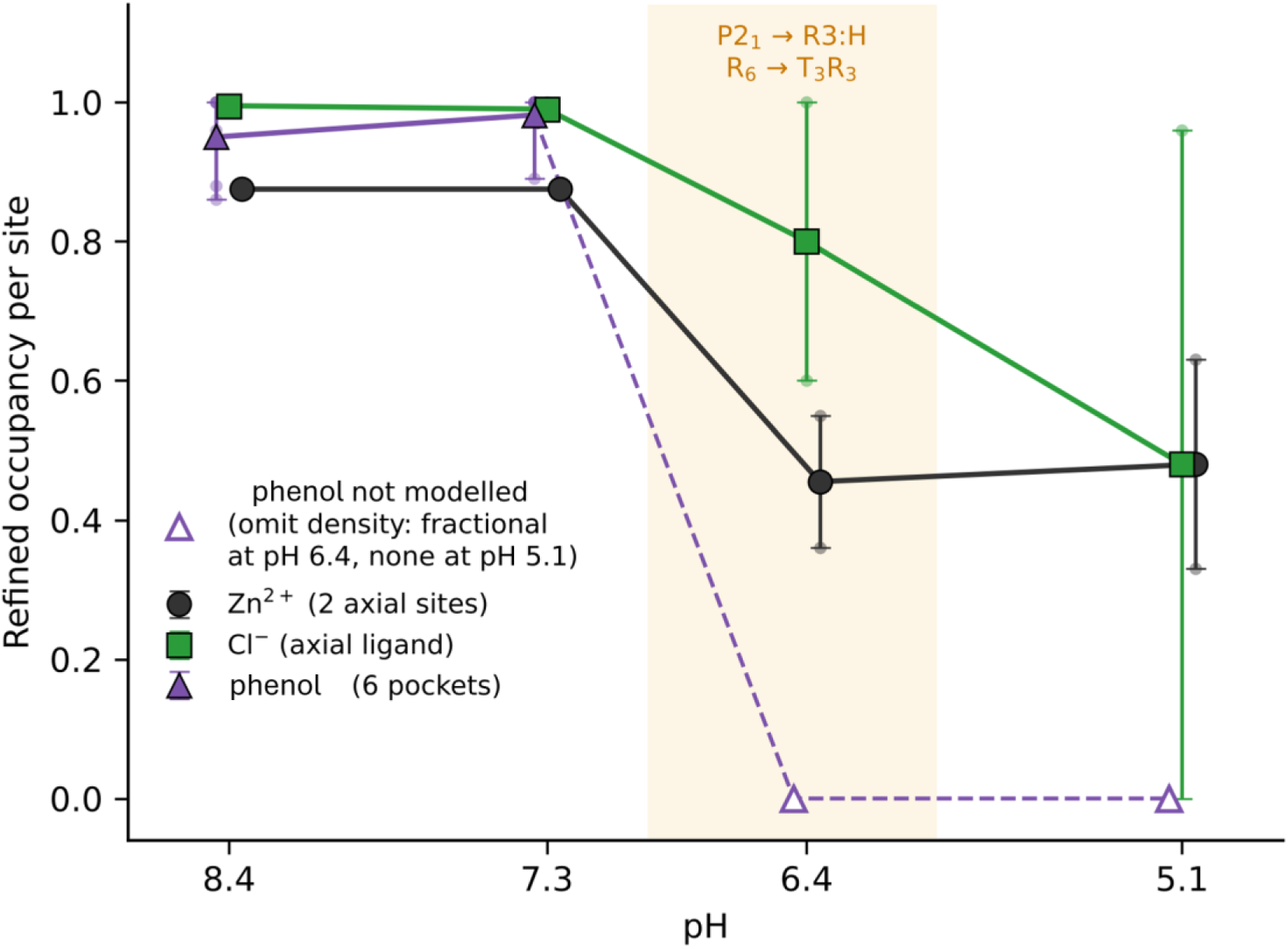
pH-dependent changes in Zn²⁺, Cl⁻, and phenol occupancy in insulin glargine assemblies. Refined occupancies of Zn²⁺ (black circles; two axial sites), Cl⁻ (green squares; axial ligands), and phenol (purple triangles; six binding pockets) across pH 8.4–5.1. Error bars indicate the range of site-specific occupancies. The shaded region highlights the principal structural transition from P12₁1 to R3:H and from Rf₆ to T₃Rf₃ between pH 7.3 and 6.4. Zn²⁺ and Cl⁻ exhibit partial, site-dependent occupancy changes upon acidification rather than complete depletion. At pH 6.4 and 5.1, phenol was not included in the refined models (open triangles); omit-map analysis revealed residual ligand-associated density at pH 6.4 but no detectable density at pH 5.1. These observations are further supported by the difference electron-density analyses in ***Fig. S8e–g***, which highlights pH-dependent changes around the metal coordination sites and B-chain N-terminal region.

On the other hand, unlike previously reported *T*-state structures rigidified by a central hydration spine (**Fig. S11b_i-vi_**) [56], our ambient-temperature SFX *T*-monomers (pH 6.4 and 5.1) lack this solvent stabilization, explaining their elevated *B*-factors of allosteric interactions (**Fig. S9c-d; Fig. S11b_vii-viii_**). Rather, their structural integrity is maintained through novel, pH-specific intra- and inter-monomer interactions. These include intra-monomer contacts Q4_B_–L6_B_, C7_B_–H10_B_, H10_B_–E13_B_, H5_B_–Q4_B_ (**Fig. S9h**); notably the rewiring of Q4_B_ from its canonical C11_A_ partner [48] to L6_B_, and a core-stabilizing H10_B_–E13_B_ interaction that helps buffer acidic protonation [57]. In parallel, specific inter-monomer polar contacts, including F1_B_–V18_D_, Q4_B_–Y16_D_, and L6_B_–C6_A_ inter-chain lock, further reinforce the assembly (**Fig. S9h; Fig. S11b_viii_; Fig. S12**). These features reveal our *T*-state not as a passive, phenol-free default [1], but as an actively stabilized alternative topology. Especially, at pH 5.1, intra-monomer interactions are further attenuated within the *T_3_*-trimer but increased the inter-monomer contacts (**Fig. S9h; Fig. S12**), accompanied by increased residue–residue cross-correlations (>90%, close to red 1.00) with a pronounced increase in B-factors (∼62.9 Å²; **Fig. S9h**). Indeed, whereas compact *R*-state subunits across all four structures show limited cooperativity (**Fig. S10** and **Fig. S11a_i-viii_**), the acidic T-state exhibits stronger collective motions (**Fig. S11b_i-viii_**), driven by inter-monomer contacts within the allosteric region of the T-subunits (**Fig. S12**) [58]. We propose that this enhanced cooperativity and structural plasticity drive an ‘unpeeling’ motion [59] that orchestrates the controlled reorganization of the hexamer. Functioning as a mechanically labile interface, this dynamic state facilitates dissociation into biologically active monomers through coherent collective oscillations. Across the entire hexamer of those glargine conformers, we observed that ***(i)*** water molecules and intra-monomer contacts decreased with decreasing pHs (**Fig. S13a**); ***(ii)*** the Poisson-Boltzmann equation reveals a striking inversion of the hexamer core properties across the pH gradient (**Fig. S13b**); and ***(iii)*** the Debye–Waller factor of the whole hexamer highlights a progressive increase (50.6 Å² to 64.5 Å²) in structural plasticity toward lower pH conditions (**Fig. S13c**). Specifically for ***ii***, at high pH (8.4 and 7.3), the hexamer core maintained a dominant positive electrostatic potential stabilized by zinc coordination; however, acidification was accompanied by an attenuation of this potential, consistent with changes in the predicted protonation equilibria (**Table 1**). This ‘electrostatic switch,’ likely associated with changes in the protonation equilibria of the central E13_B_ cluster [2], [56], [60] and H10_B_ imidazole groups, modulates core electrostatic interactions and contributes to the conformational allostery observed in the diffraction data. Furthermore, our surface accessibility (SA) and molecular surface topology (MS) analyses, which are calculated by the *CASTpFold* server [61] (**Fig. S14**), reveal a precise remodeling of the surface landscape. At near-physiological pHs, the linear correlation between solvent-accessible and molecular surfaces reflects a compact, probably ligand-locked assembly dominated by charged and aromatic residues (**Fig. S14a–b**). Upon acidification, while the global correlation profile remains largely conserved—indicating the retention of the hexameric core—residue-wise distributions expose a specific exchange of surface residues. This reorganization is marked first by the emergence of the phenol-gatekeeper I10_A_ [11], [62], [63] at pH 6.4 (**Fig. S14c**), followed at pH 5.1 by the pronounced exposure of the N-terminal anchor Q4_B_ [64], consistent with long-range Nuclear Overhauser Effects reported for the B-chain N-terminal arm that preserve T-state-specific interactions [65], along with the loss of a prominent Tyrosine (Tyr) peak observed at higher pHs (**Fig. S14d**), giving rise to characteristic upfield shifts that are lost in the T-state [32]. Collectively, these crystallographic data profiles confirm a transition from a rigid storage state to a heterogeneous plasticity involving both helix ‘unpeeling’ [59] and hydrophobic redistribution. To independently assess whether current structure prediction methods capture the ligand-associated conformational states observed by SFX, we performed AlphaFold3 (AF3) modeling of insulin glargine in the presence and absence of phenol (**Fig. 3**). In the presence of six phenol molecules and two Zn²⁺ ions, AF3 predicted an Rf6-like hexamer, closely matching the SFX structures at pH 8.4 and 7.3. Individual protomer superpositions yielded RMSD values of 0.262–0.393 Å and 0.226–0.406 Å, respectively, with a whole-hexamer RMSD of 0.316 Å against the pH 8.4 structure. In contrast, to mimic the crystallographic asymmetric units of the acidic SFX structures, phenol-free AF3 modeling was performed using two glargine protomers and Zn²⁺ ions. Strikingly, both AF3-predicted protomers adopted T-like conformations, whereas the experimentally resolved asymmetric units at pH 6.4 and 5.1 contain one T-state and one Rf-state protomer, forming mixed T3Rf3 hexamers through crystallographic symmetry. Despite local structural similarities (RMSD values of 0.564–0.831 Å and 0.858–1.406 Å, respectively), the phenol-free AF3 prediction exhibited high confidence scores (ipTM = 0.88 and pTM = 0.89) yet failed to reproduce the experimentally observed T/Rf conformational asymmetry. More broadly, these findings suggest that phenol availability may influence the conformational preferences predicted by AF3, while highlighting the limitations of current structure prediction methods in resolving intermediate allosteric states governed by ligand binding, metal coordination, and pH-dependent interactions. These results suggest that phenol availability may influence the predicted conformational preference, while highlighting the limitations of AF3 in capturing mixed allosteric states and the importance of experimental SFX measurements for resolving these structurally distinct intermediates.

**Figure 3.**
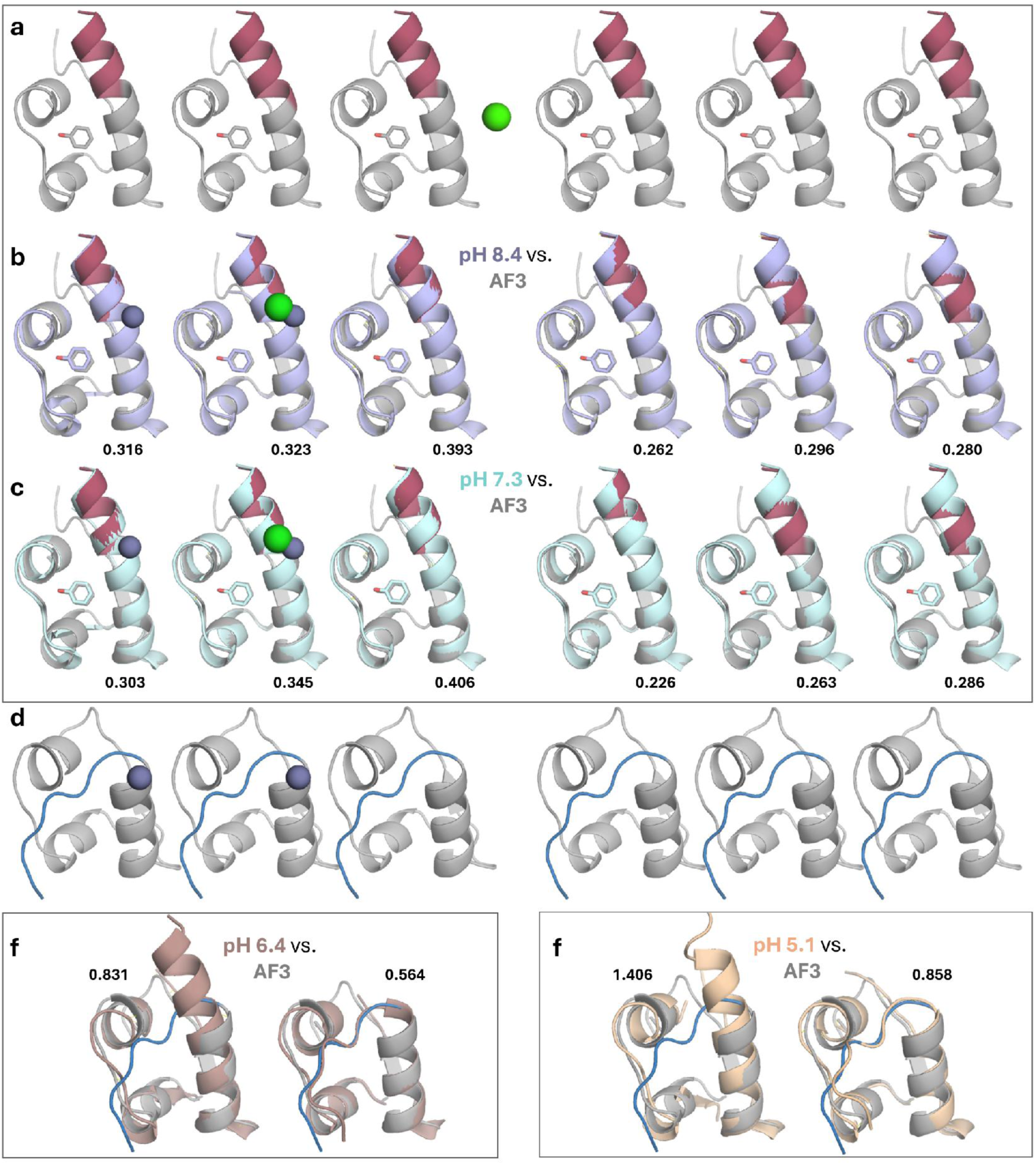
AlphaFold3 captures ligand-associated conformational preferences but misses the experimentally observed allosteric intermediate insulin glargine. **a,** AF3-predicted phenol-bound glargine hexamer, showing six Rf-like protomers. **b-c**, Structural superpositions of individual AF3 protomers with SFX structures determined at pH 8.4 and 7.3, respectively, demonstrating close agreement in the Rf conformation. **d,** Phenol-free AF3 prediction of a glargine dimer, with both protomers adopting T-like conformations. **e-f,** Structural comparisons with the experimentally resolved asymmetric-unit dimers at pH 6.4 and 5.1, respectively.

### 2.2 pH induces molten globule-like conformation in-solution, supporting in-crystallo allostery

To investigate whether the pH-dependent structural changes observed by SFX are accompanied by corresponding changes in solution, we performed complementary biophysical measurements using insulin glargine samples prepared under comparable pH conditions. Size-exclusion chromatography, Raman spectroscopy, differential scanning calorimetry, and intrinsic fluorescence were used to assess oligomeric heterogeneity, backbone conformation, thermal stability, and local aromatic environments. Together, these measurements provide solution-based evidence of pH-dependent structural reorganization, complementing the crystallographic observations.

Initially, we performed size-exclusion chromatography (SEC) (**Fig. 4a**) using insulin crystals formed at pH 8.4 or pH 5.1, which were gently redissolved in the SEC buffer at the same pH prior to analysis (**Fig. 4b**), alongside insulin solubilized at pH 2.1 as a reference [66]. SEC profiles show that insulin at pH 8.4 elutes predominantly as higher–molecular-weight species, consistent with stable hexameric assemblies. In contrast, the pH 5.1 sample displays a broader distribution spanning both hexameric and lower-molecular-weight regions, indicative of complex oligomeric heterogeneity, whereas insulin at pH 2.1 elutes as lower-molecular-weight forms (fully dimeric) (**Fig. 4a**) [66]. Notably, control experiments performed without prior crystallization, using direct buffer exchange into the corresponding pH conditions, yielded consistent SEC profiles, particularly at pH 5.1, where a similarly broad distribution of oligomeric species was observed (**Fig. S15**, *fractions(F) 1-3*). These results indicate that while the hexameric assembly is preserved at pH 8.4, resolubilization at pH 5.1 induces conformational reorganization into intermediate oligomers, in agreement with the crystalline states captured by SFX (**Fig. 1b-l**).

**Figure 4.**
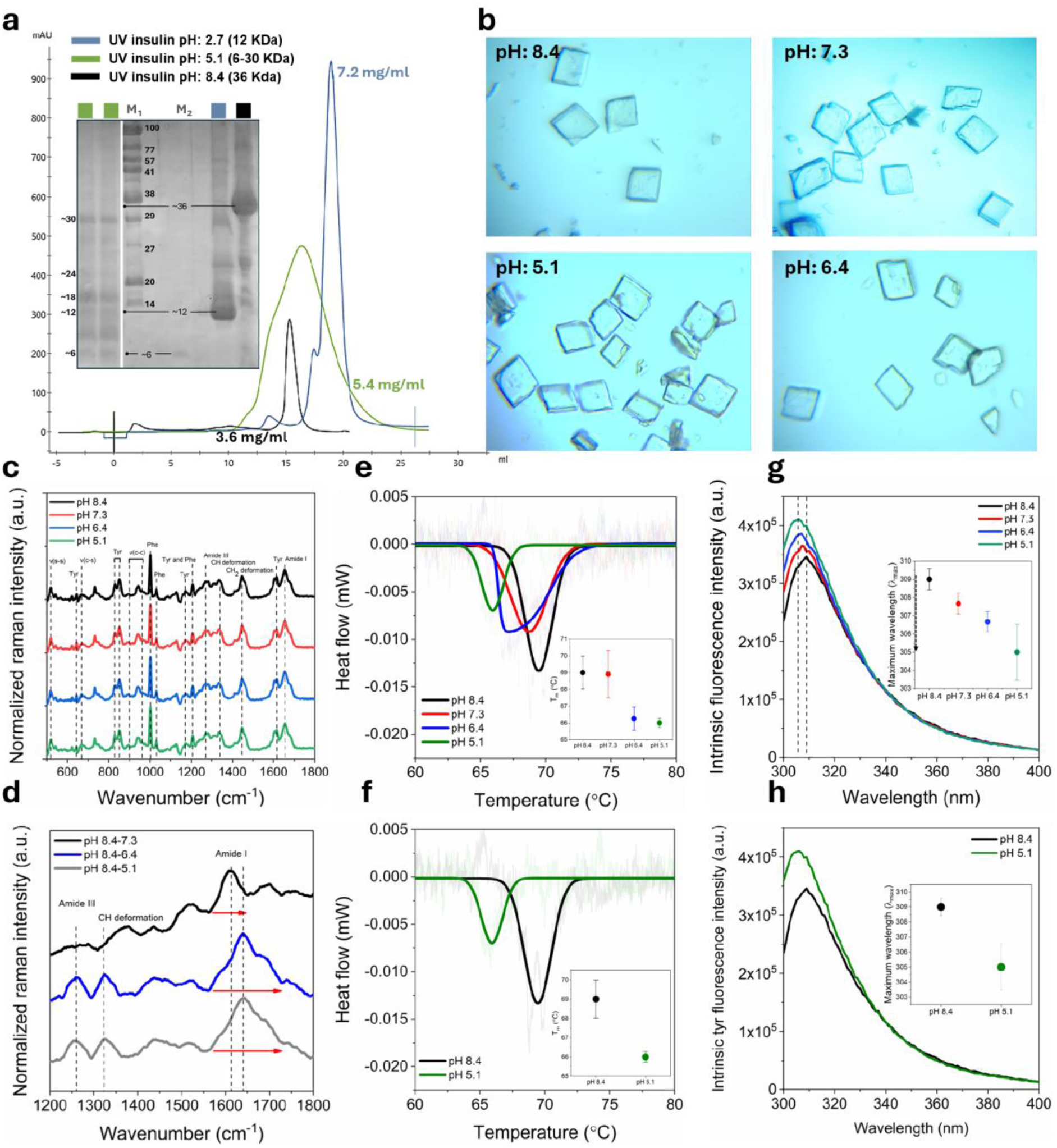
pH-dependent *in-solution* spectroscopic and chromatographic signatures of insulin glargine destabilization. **a,** SEC profiles of resolubilized glargine crystals at pH 8.4 and 5.1, with pH 2.7 as reference: pH 8.4 is eluted mainly as stable hexamer, whereas pH 5.1 shows broadened, heterogeneous oligomeric populations (hexamer to monomer). Peak fractions were analyzed by SDS-PAGE (M, marker). **b,** Representative crystal micrographs before resolubilization for SEC, Raman, and 20% SDS-PAGE analysis. **c-d,** Raman spectra across pH 8.4–5.1 and difference spectra (vs. pH 8.4) highlighting changes in Amide I/III, ν(C-C), Tyr Fermi doublets, and ν(S-S)/ν(C-S); Amide I broadening indicates helical disorder and increased conformational fluctuation. **e-f,** DSC thermal unfolding profiles show acidification-dependent loss of stability, with T_m_ values summarized in insets. **g-h,** Intrinsic fluorescence (Tyr/Phe, λex = 280 nm) shows a 4 nm hypsochromic shift in λmax (309→305 nm) at acidic pH, consistent with aromatic residue sequestration in a more hydrophobic, molten-like environment; insets show λmax versus pH values.

Raman spectra of single insulin crystals were collected under identical pH conditions (pH 8.4, 7.2, 6.3, and 5.1) over the range of 500–1800 cm⁻¹ (**Fig. 4c,d**) [67]. The Raman profiles capture pH-dependent chemical and conformational changes in insulin (**Fig. 4c**) [68]. To further observe α-helical alterations of residues 1–8 in the B chain, difference Raman spectra were calculated in the α-helical region (1200–1800 cm⁻¹) (**Fig. 4d**). The amide I band, spanning ∼1600–1670 cm⁻¹, arises predominantly from C=O stretching vibrations of the protein backbone, with minor contributions from C–N stretching and N–H in-plane bending (**Fig. 4c**) [69], [70]. In the blue (pH 8.4–6.4) and grey (pH 8.4–5.1) difference spectra, the amide I peak shifts from ∼1612 to ∼1640 cm⁻¹, indicating an increase in vibrational frequency associated with secondary structural elements. In addition, the band near 1640 cm⁻¹, within the amide I region, may reflect contributions from α-helical rearrangement and overlapping disordered structural components (**Fig. 4d**). Notably, peak broadening (red arrow) is observed in both difference spectra, consistent with less uniform hydrogen-bonding patterns along the insulin backbone [71], which is in line with our crystallographic data (**Fig. S9e-h**). In addition, CH deformation modes appear in the 1300–1350 cm⁻¹ region (**Fig. 4d**) [71]. Both blue and grey difference spectra show the emergence and increased intensity of a CH deformation peak at ∼1323 cm⁻¹, indicative of higher-frequency vibrational contributions accompanying the conformational transition (**Fig. 4d**), which is supported by our cross-correlation analysis (**Fig. S9h; Fig. S12e-f**). To further study the glargine insulin, we performed in-solution differential scanning calorimetry (DSC) and intrinsic fluorescence experiments. Insulin glargine (Lantus®) hexamers are maintained at acidic pH (pH 4). Upon transfer of insulin glargine solution to neutral or alkaline pH buffer environment, significant changes in self-association behavior can occur, resulting in alterations of the oligomer distribution from hexamer to possibly monomer, which allow us to study conformational changes at the level of single-molecule glargine.

To assess the thermal stability of glargine samples under identical pH conditions (pH 8.4 to 5.1), differential scanning calorimetry (DSC) measurements were performed. Representative calorimetric profiles of insulin across (**Fig. S16**) all the tested pH conditions (pHs 8.4, 7.3, 6.4, 5.1) are shown in **Figure 4e–f**, illustrating heat-flow changes as a function of temperature, with the melting temperature (*T_m_*) defined as the point at which 50% of the protein population is denatured [8]. DSC measurements revealed pronounced differences in the thermodynamic properties at pH 8.4 and 5.1 (**Fig. 4f**). At pH 8.4, thermal unfolding yielded a T_m_ of 69.0 ± 0.98 °C, which can be attributed to glargine monomer as it was presented earlier, and a calorimetric enthalpy (ΔH_cal_) of 28.24 ± 6.91 kJ mol⁻¹. In contrast, insulin glargine at pH 5.1 exhibited a markedly reduced thermal stability, with a T_m_ of 66.0 ± 0.28 °C and a ΔH_cal_ of 13.63 ± 0.71 kJ mol⁻¹. The observed decrease in melting temperature (ΔT_m_ ≈ 3 °C) and enthalpy change (ΔH ≈ 14.61 kJ mol⁻¹) indicates a substantial reduction in conformational stability at acidic pH (**Fig. 4f**, *inset*). Collectively, these calorimetric data demonstrate that monomer insulin undergoes moderate thermal destabilization at pH 5.1, consistent with increased conformational plasticity (**Fig. S13c**) and supporting the pH-induced allosteric reorganization of the hexamer observed in crystallographic data (**Fig. 1c-f**).

Intrinsic fluorescence emission spectra were also recorded using insulin glargine under identical pH conditions (pH 8.4 to 5.1) (**Fig. 4g–h**). Insulin contains four tyrosine and three phenylalanine residues in its native structure, whose intrinsic fluorescence sensitively reports on changes in the local aromatic microenvironment [72], [73]. We observed a progressive increase in fluorescence intensity toward decreasing pH (**Fig. 4g**). More notably, the emission maximum shifted by ∼4 nm, from 309 to 305 nm (**Fig. 4h**, *inset*). Photophysically, this blue shift reflects a shift toward a less polar microenvironment [74], [75], [76] surrounding the emitting aromatic residues, consistent with subtle conformational rearrangements due to the decreased number of water molecules at lower pHs (**Fig. S13a**). Our structural mapping of Tyr and Phe residues reveals pronounced pH-dependent reorientation and repacking of aromatic side chains, providing a mechanistic basis for the observed changes in intrinsic fluorescence (**Fig. S17**).

Consistent with the pH-dependent fluorescence changes, the predicted cross-correlation patterns suggest enhanced collective coupling within the T-state allosteric region at pH 6.4 and 5.1 (**Fig. S9h; Fig. S11b_vii-viii_**). Together, these observations support increased conformational heterogeneity in the T₃Rf₃ hexamer, with features consistent with a molten-globule-like assembly [77] [78]. This spectroscopic finding is also supported by our crystallographic surface topology (SA/MS) analyses (**Fig. S14**), which reveal pH-dependent redistribution of aromatic residues. Indeed, while the appearance of the phenol-gatekeeper I10_A_ as a dominant surface residue at pH 6.4 signifies the initial disruption of the phenolic pocket, the reorganization at pH 5.1 is defined by the pronounced exposure of the N-terminal anchor Q4_B_, along with the repositioning of a prominent tyrosine residue (Y14_A_) from the solvent-accessible surface (**Fig. S14d**).

Collectively, here *in-solution* data (SEC, Raman, DSC, fluorescence) validate the pH-dependent conformational reorganizations observed by SFX, confirming that our crystallographic models capture an authentic intermediate driven by aromatic and tertiary reorganization rather than lattice artifacts. In sum, the ‘unpeeling’ motion [59] that we observe in crystallographic data provides the structural rationale for the molten-globule-like state [72]–secondary structure is preserved while tertiary structure is loosened– observed *in-solution*.

### 2.3 In-silico intrinsic mode redistribution confirms pH-dependent lattice allostery

Once the crystallographic observations were corroborated by *in-solution* experiments, we conducted *in-silico* analyses of the SFX data to characterize the dynamic behavior of insulin hexamers. Specifically, we sought to ***(i)*** examine the conformational changes under dynamic in-solution conditions underlying the static crystallographic structures, ***(ii)*** relate these changes to increased conformational flexibility supported by the solution measurements, and ***(iii)*** compare the resulting dynamical signatures with those inferred from available PDB structures. Accordingly, we applied normal-mode analysis (GNM and ANM), GNM-TECol, predicted diffuse scattering, and principal component analysis (PCA) (**Fig. 5**). GNM provides an analytical description of the normal-mode spectrum encoded by the inter-residue contact topology. In multimeric assemblies, GNM-predicted cross-correlations across structural elements and length scales can provide insights into potential allosteric communication pathways and the influence of oligomerization on collective dynamics [79], [80], [81]. Moreover, the mode spectrum can be separated into collective low-frequency (global) motions and high-frequency (local) fluctuations, the latter often highlighting regions of localized energetic or structural rigidity [82].

**Figure 5.**
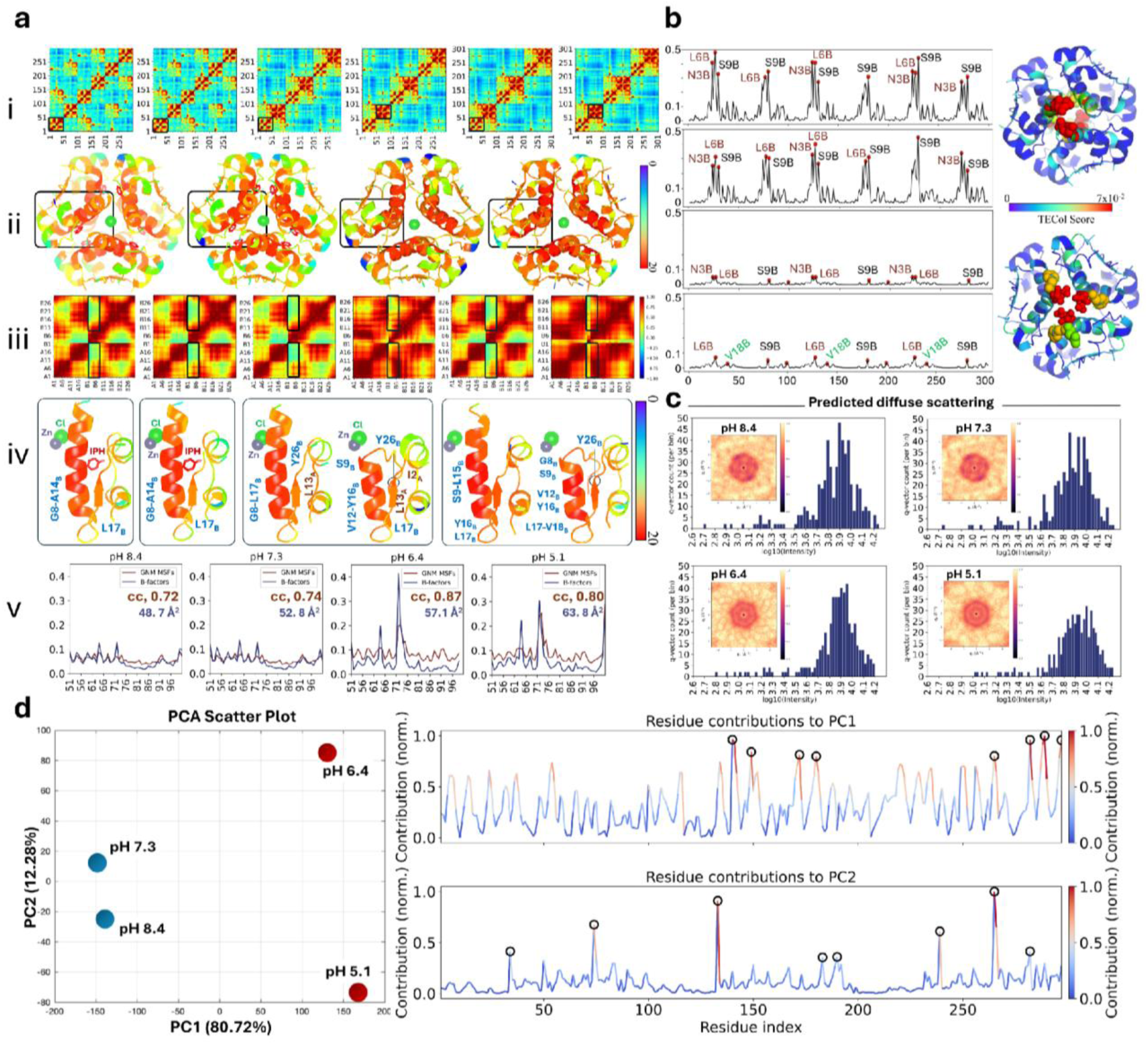
*In-silico* analyses confirm the pH-dependent lattice transition. **a,** GNM soft-mode cross-correlation maps for pH 8.4–5.1; (inset i-iii) monomer regions are boxed, (inset ii) with zoomed 3D monomer maps, (inset iii-iv) allosteric-to-monomer correlations, and (inset iv) hinge residues annotated across *R* and *R/T* subunits. (inset v) GNM means-square fluctuations (brown) align with experimental *B*-factors (blue) for A and B chains. **b,** TECol scores for slow modes (GNM modes 3–5) across pH, with residue index on the x-axis and TECol score on the y-axis; top residues highlighted in TECol highest score plots and mapped in 3D for P12_1_1 (*first and second panels*) and R3:H (*third and fourth panels*) with jet coloring and sphere markers. The 3D structures at all pH conditions were colored using the same TECol spectrum range (0–0.07), originally applied to the low-pH structures. TECol scores exceeding 0.07 are displayed at the upper limit of the color scale. **c,** Predicted diffuse scattering from covariance-based Gaussian modeling: 2D maps remain isotropic, while log _10_(*I*) histograms show pH-dependent amplitude rescaling (narrowing at pH 6.4; higher-intensity shift at pH 5.1). **d,** PCA of hexamer dynamics: PC1–PC2 separates P12_1_1 (pH 8.4–7.3) from R3:H (pH 6.4–5.1); residue loadings highlight F1, S9, V12, L16, C19, P28 (PC1) and N3, V18 (PC2), with color intensity indicating normalized loading.

Initially, we performed cross-correlation analysis using the entire hexamer over the pH range from 8.4 to 5.1 (**Fig. 5_i_**). For detailed visualization of the cross-correlation patterns, representative monomers were selected at pH 8.4–7.3 as all six protomers adopted the Rf-state conformation, whereas representative dimers containing both Rf- and T-state protomers were selected at pH 6.4 and 5.1 (**Fig. 5_i__-ii_**). Interestingly, we observed a rise in collectivity of the *T*-state protomers at pH 6.4-5.1, while all *Rf*-state protomers at both low and near-physiological pHs show similar collective dynamics, exhibiting relatively lower cooperativity (**Fig. 5a_ii__-iv_**). This aligns with the minima and maxima observed in the soft (slow) and high-frequency (fast) modes of the first dimers, emphasizing their roles as hinges and peaks, respectively, that mediate the global and local dynamics of the glargine hexamer (**Fig. S18**). The peaks in fluctuations driven by fast modes [83] are usually located in the insulin core and tend to be evolutionarily conserved [84], whereas the hinge regions driven by soft modes represent the most accessible motions to elicit allosteric responses [85]. The observed differences in the cross-correlations driven by the soft modes between the structures are also consistent with the previously reported dynamicity of the *T_3_R_3_*insulin structures. All *R*-subunits have shown moderate correlation within the monomer, whereas the *T* subunit shows a much stronger correlation (**Fig. S19**). To assess the robustness of the GNM-derived cross-correlation patterns, we performed a coordinate-perturbation sensitivity analysis using 50 alternative models for each pH condition (σ = 0.2 Å). The resulting cross-correlation matrices showed consistently high similarity to their corresponding original structures (Pearson r = 0.9899–0.9939), indicating that the overall correlation patterns are largely preserved under small structural perturbations (**Fig. S20**). This analysis evaluates model sensitivity rather than reproducibility across independently processed experimental datasets.

Consistently, a gradual and pronounced increase in the *Debye-Waller* factor that we observed at lower pHs is also in line with the GNM mean-square fluctuation (predicted B-factor) profile (**Fig. 5a_v_**). *T*-state has been characterized by vibrational thermal motion and static disorder, leading to a gradual increase in the mean-square displacements [86]. The fact that the experimental *B*-factor, supported by the GNM MSF profile, stresses the pronounced plasticity of the structure, not caused by crystal packing but rather by a mechanistic view originating in *unpeeling* motion [59] and thermal-vibration-induced conformational plasticity. This provides biophysical proof of the *molten insulin globule* [1].

To identify the causal interrelations among those dynamics, we employed Transfer Entropy–Collectivity (TECol) analysis to map residues with dynamic capacity to transfer information to other residues across the hexameric insulin structures (**Fig. 5b**). TECol scores revealed distinct pH-dependent shifts in residue-level information flow, consistent with the soft and fast mode profiles described earlier (**Fig. S18**). At pH 8.4 and 7.3, the prominent effector residues N3_B_, L6_B_, and S9_B_, which localized to the allosteric site, were detected as the primary mechanical drivers. This coincides with hinge zones defined by soft collective GNM modes (**Fig. S18a-b**; L6_B_–V18_B_) and involving key intra-monomer allosteric regulators (**Fig. S9e-f**; e.g., S9, H10, L11, V12, E13) (**Fig. 5b**, *first and second panels*). Conversely, the transition toward the *T_3_R_3_*-state (pH 6.4 and 5.1) triggered a reorganization (**Fig. 5b**, *third and fourth panels*). The appearance of additional relative TECol hotspot (green, V18_B_) suggests a redistribution of residue-level contributions to dynamic coupling toward the central/C-terminal B-chain segments, overlapping with the hinges (**Fig. S18c-d**; L15_B_–Y26_B_) and intermolecular interactions of the *T*-subunit’s allosteric regions (**Fig. S9h** and **Fig. S12**; Y16_B_, V18_B_). The spatial mapping of TECol scores onto the hexamer cartoon structure highlights the redistribution: while the effector residues at pH 8.4–7.3 cluster near the dimer interface and N-terminal turns, pH 6.4–5.1 ones radiate toward more solvent-exposed and interfacial domains, paralleling MSSA-derived surface topology at lower pHs (**Fig. 14c-d**; e.g., I10_A_, Y14_A_, Q4_B_). This strongly supports a ‘compensatory stabilization’ mechanism: as the helical rigidity (L6_B_–V18_B_) gradually *molten*, the dynamic drive of the hexamer shifted toward source residues V18_B_, P28_B_ alongside N3_B_, L6_B_, and S9_B_, and the C-terminal anchors (F24_B_–Y26_B_) to sustain coherent collective oscillations. This redistribution is highly consistent with the early onset of the hexamer detachment process [87].

To connect the pH-dependent redistribution of local dynamics to experimentally observable crystallographic signatures, we simulated X-ray diffuse scattering for the insulin hexamers (**Fig. 5c**). Using a custom Gaussian-approximation workflow [88], [89], GNM-derived covariance matrices [90] and atomic displacement parameters were projected into reciprocal space to generate 2D diffuse-scattering maps for the four pH conditions, together with accompanying *log_10_(I)* intensity histograms over the sampled *q*-vectors. Across all pH conditions, the predicted 2D diffuse-scattering maps remain predominantly radial and near-isotropic within the present Gaussian Cα framework, indicating that the modeled disorder is dominated by amplitude-like fluctuations rather than strongly directional spatial correlations. Likewise, no pronounced streak-like features are observed along specific reciprocal-space directions in the simulated patterns. Conversely, the accompanying intensity histograms show clear pH dependence: profiles at pH 8.4 and 7.3 are broadly similar, the distribution shows relative narrows at pH 6.4, and at pH 5.1 the peak shifts toward higher intensity (∼3.9 vs ∼4.0) with a more extended high-intensity tail. Taken together, these trends suggest that the principal pH effect in this model is a redistribution of effective fluctuation amplitudes encoded in the covariance matrix, rather than a major change in global map geometry. More cautiously, because the current implementation uses Cα coarse-graining, isotropic displacement assumptions, and display normalization steps, these results are best interpreted as model-consistent evidence for pH-dependent amplitude modulation, to be refined further by higher-resolution anisotropic treatments.

Consistent with the pH-dependent redistribution of dynamic amplitudes concluded from covariance rescaling, principal component analysis (PCA) clarifies how these collective motions are encoded across the hexamer. The PC1–PC2 projection separates the P12_1_1 structures (pH 8.4–7.3) from the R3:H structures (pH 6.4–5.1), suggesting a coherent shift in dominant conformational coordinates rather than stochastic structural noise (**Fig. 5d**, left). Residue-wise profiles indicate that the main variance (PC1) concentrates at F1, S9, V12, L16, C19, and P28, with additional contributions from N3 and V18 along PC2 (**Fig. 5d**, right). These hotspots align with residues identified as hinges or dynamic effectors in the GNM cross-correlation spectrum (**Fig. 5a_iv_**) and as high-transfer nodes in the TECol analysis (**Fig. 5b**). Taken together, these complementary structure-based analyses suggest that the pH-driven space-group transition is consistent with a redistribution of pre-existing intrinsic modes, rather than the appearance of new motional regimes. The transition from P1211 to R3:H can therefore be viewed as a reweighting of the hexamer’s internal communication network, in which fluctuations that are less prominent at higher pH become amplified under acidic conditions. Agreement among PCA, GNM (**Fig. 5a_iv_**), TECol (**Fig. 5b**), predicted B-factors (**Fig. 5a_v_**), and diffuse-scattering scaling (**Fig. 5c**) provides a unified *in-silico* framework: lowering pH shifts the balance from helical rigidity toward C-terminal plasticity, promoting coordinated, *unpeeling*-dependent conformational plasticity within the *T_3_R_3_* assembly.

To assess whether the observed *R* □*T* transition reflects a novel dynamical regime or selection of pre-existing intrinsic motions, we projected the pH-dependent structures onto the low-frequency conformational space defined by ANM (**Fig. 6**). Notably, the R₃ core remains structurally conserved between pH 6.4 and 5.1, despite increased atomic displacement parameters and a modest change in the buried allosteric region (∼18.8–19.0 Å). ANM modes 1 and 3 further indicate broadly similar predicted collective-motion patterns across these acidic states (**Fig. S21; Movie S1**). The geometric analysis shows a systematic reorganization of core angles and burial/exposure states during the *R*-to-*T* shift (**Fig. 6a–h**). In particular, the hinge-opening motion—evidenced by expansion of the F1_B_ span from ∼18.7 Å to ∼32.4 Å, consistent with the previously reported ∼30 Å displacement of F1_B_ [1]—paradoxically yields an “exposed” *T*-state with markedly fewer coordinated water molecules than the heavily hydrated, tightly “buried” *R_6_*-state core (**Fig. S16a-b**) [56]. These conformational changes align with intrinsic low-frequency pathways encoded in ANM modes of the allosteric regions alone (**Fig. 6m–p; Fig. S22; Movie S1**), indicating that the acidic transition exploits a pre-existing collective motion rather than a random structural trajectory. Holistic vector schematics (**Fig. 6r-s**) detail this mechanical evolution: the highly symmetric, inward-breathing dynamics at pH 8.4 progressively accumulate asymmetric rotational deviations (red arrows) at pH 7.3 [9] (**Fig. 6r**), which develops into an outward-twisting expansion at lowest pH (**Fig. 6s**). Consistently, multidimensional folding funnel (**Fig. 6t-u**), mapped along the collective order parameters of topological order (Q) and compactness (Z) [91], show that pH lowering does not introduce an isolated dynamical basin but reshapes the energetic funnel along these same intrinsic coordinates [92]. Specifically, the contour map (**Fig. 6u**) shows the pH 8.4 and 7.3 structures occupying distinct minima in the bottom-right quadrant, characterized by high Q and low Z. With acidification, the pH 6.4 and 5.1 structures shift toward the top-left quadrant, where Q decreases and Z increases. Importantly, all states remain within a single continuous landscape, sharing the same underlying funnel topology. Ultimately, these results indicate that the *molten*-like state in *T_3_R_3_* does not arise from stochastic denaturation; rather, it reflects a thermodynamically driven, directed reorganization of latent collective modes that orchestrates the *unpeeling* mechanism required for initializing the hexamer dissociation [1], [59].

**Figure 6.**
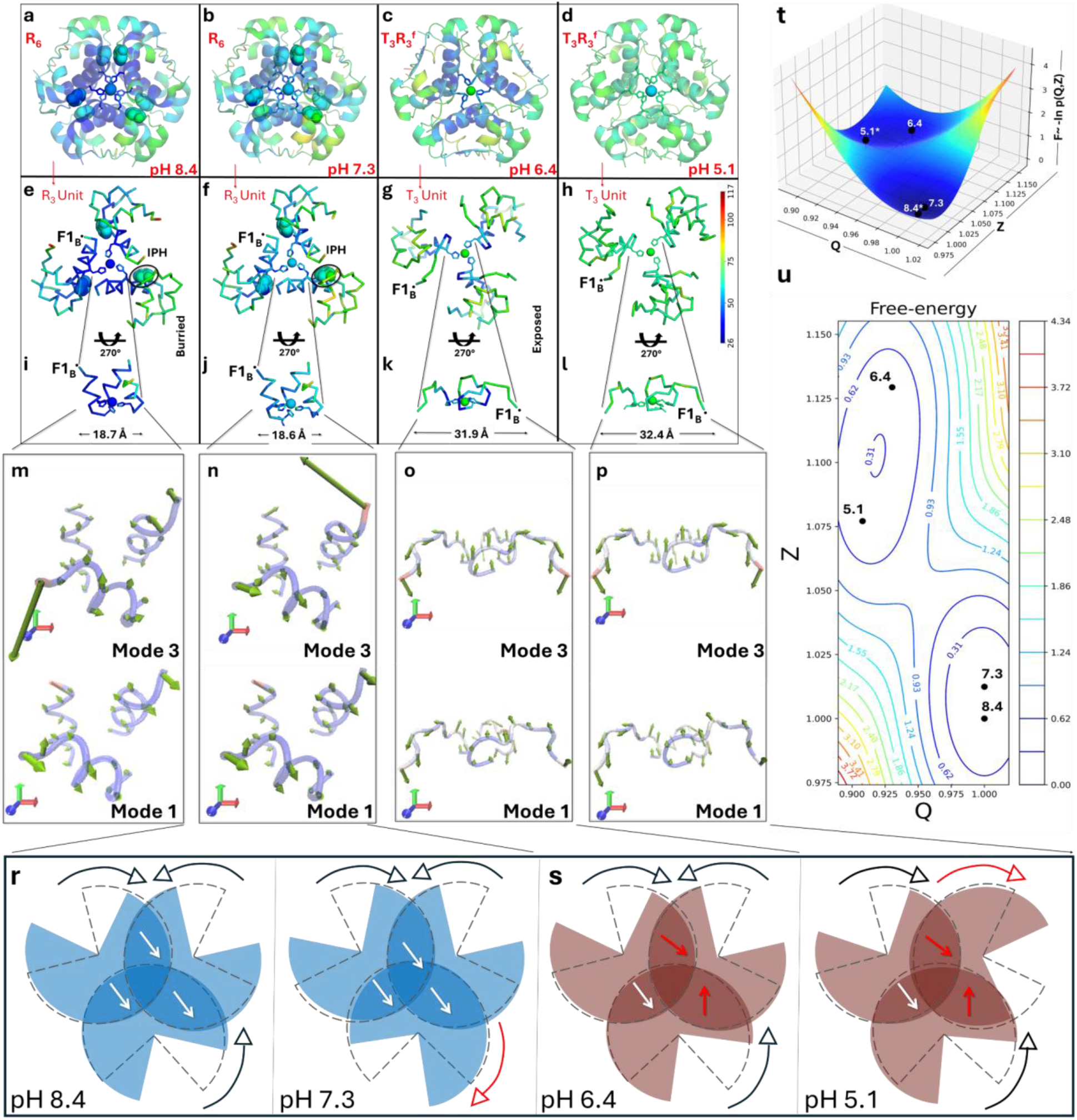
Structural, dynamic, and thermodynamic analyses describe the underlying mechanisms of reorganization during the molten-like *R*→*T* transition. a-d, Hexamer cartoons colored by *B*-factor show increasing plasticity with decreasing pH. **e-h,** Trimer ribbon views at pH 8.4–5.1; i-j, *R_3_* architecture with phenol stabilizes a buried allosteric state (∼18 Å); **k-l,** whereas *T_3_* lacks phenol and adopts an exposed state (∼32 Å). **m–p,** ANM of the allosteric regions (modes 1 and 3). **r–s,** Vector representation of hexamer ANM mode 1; red arrows mark reorientation relative to the pH 8.4 SFX structure. **t,** 3D free-energy surface F=−ln*P*(*Q,Z*) in the Onuchic–Wolynes Q–Z framework (Q: native contacts; Z: compactness); colors indicate free energy (blue low → red high), black markers show pH 8.4, 7.3, 6.4, 5.1. Near-neutral pH states cluster near the minimum, whereas acidic states shift along the same Q–Z coordinates. **u,** 2D Q–Z contour map with iso-free-energy lines (0–4.34).

Finally, to understand whether the above-observed collective reorganization is in line with previously reported *glargine* structures, we compared our SFX glargine datasets with available insulin glargine entries in the PDB *in-silico* (**Fig. S23**). Overall, no major qualitative structural deviations are evident across the reported models of 4IYD, 5VIZ, 6K59, and 8WU0 in PDB. In the *R*-subunits of previously reported structure (8WU0: R3:H lattice; dimer in ASU), relatively weak collective motions have been reported, comparable to the cooperativity observed in our SFX-derived *R_6_* form (P12_1_1 lattice; hexamer in ASU). In contrast, *T*-subunits from previously reported structures (4IYD, 5VIZ: I2_1_3 lattice; monomer in ASU) display stronger collective behavior, relatively resembling the collectivity seen in our *T_3_R_3_* form (R3:H lattice; dimer in ASU). However, the *T*-subunits in our SFX structures exhibit reduced cooperativity, probably due to the enhanced structural plasticity under more acidic conditions.

## 3. Discussion

### Reinterpreting the protracted glargine dynamicity through near-physiological SFX

The prolonged pharmacokinetics of insulin glargine are generally attributed to its pI shift toward neutrality: the acidic formulation remains soluble in the pen, whereas exposure to physiological pH promotes isoelectric precipitation after subcutaneous injection [93]. However, the molecular organization of the resulting depot, and the structural determinants of its slow release, have remained incompletely resolved, in part because prior structural descriptions have relied mainly on cryogenic snapshots and partial oligomeric states (PDB IDs: 4IYD, 5VIZ, 6K59).

Glargine was selected as the primary model because it occupies a unique mechanistic and translational position among basal insulin analogs [94]. Unlike detemir [95] and icodec [96], whose protracted action is dominated by fatty-acid-mediated albumin binding and altered systemic disposition, glargine is the canonical precipitation-driven basal insulin: after injection, pH neutralization induces depot formation, and delayed glycemic coverage depends on subsequent structural reorganization and release from this depot [93]. Compared with detemir (and mechanistically distinct agents such as icodec), glargine provides more consistent near-peakless action [97], [98], ∼24-hour basal coverage [98], stronger overall glucose-lowering/antilipolytic effects in key comparisons [99], and is commonly used with once-daily dosing [100]. Its evolution from Gla-100 to Gla-300 further demonstrates improved PK/PD stability and prolonged action [101]. As the first recombinant long-acting analog designed around isoelectric precipitation and one of the most widely used basal insulins for more than two decades in millions of patients, glargine provides a high-impact clinical benchmark and a tractable structural model for studying precipitation-coupled allostery, biosimilar comparability, and next-generation basal insulin design [94].

This focus is not only clinically justified but also biotechnologically timely. Despite extensive use, the structural basis linking depot formation to controlled dissociation has remained incompletely resolved. Defining this pathway has immediate implications for biosimilar comparability, where higher-order structure, conformational plasticity, and stability are increasingly recognized as critical quality attributes [102]. More broadly, investigating whether pH-dependent allosteric redistribution contributes to delayed release may provide a framework for future studies of basal insulin stability and release kinetics. In this sense, glargine is not simply a legacy therapeutic; it is a tractable structural prototype for precipitation-based sustained protein delivery.

Here, by integrating ambient temperature SFX with solution biophysics and multiscale network analyses, we provide an experimentally grounded framework linking glargine behavior to pH-dependent allosteric redistribution within the hexamer (**Fig. 1-6**). Across pH 8.4-5.1 at 298 K, the SFX structures separate into two reproducible regimes: near-neutral conditions favor R_6_-like/R^f^-dominant hexamers with phenolic occupancy (**Fig. 1i-j**) [5], [103], whereas acidic conditions favor rhombohedral TR^f^/T_3_R_3_-state organization with collapsed phenol pockets and reduced/absent phenolic ligands (**Fig. 1k-l**) [5]. This shift is accompanied by coordinated B-chain N-terminal remodeling (B1-B8), including characteristic F1_B_ displacement and helical *unpeeling* of the allosteric region (**Fig. S12**) [1], transitioning from a compact ligand-stabilized architecture (**Fig. S9e**) to a more exposed yet structurally coherent state (**Fig. S9h**). A central finding is that the acidic ensemble is not well described as passive ligand loss or global unfolding. Rather, it appears as a partially reorganized state stabilized by pH-dependent interaction rewiring (**Fig. S9e-h**), hydration redistribution (**Fig. S17a-b**), and electrostatic rebalancing (**Fig. S13b**). Model-independent difference maps quantify the metal-site component of this rebalancing as partial, site-dependent zinc depletion (occupancy 0.87 → 0.36 and 0.55 at pH 6.4, 0.63 and 0.33 at pH 5.1; **Fig. S8e-g**) with chloride replacing cresol at the R-state zinc and show that pH 5.1 is a more disordered form of the pH 6.4 intermediate rather than a distinct state (**Fig. 2**). This interpretation is consistent with increased disorder/flexibility metrics (**Fig. 5a_v_**; **Fig. 6a-d**) and surface-topology changes observed at lower pH (**Fig. S14c-d**), while preserving a coherent hexameric scaffold (**Fig. 6r,s**) [104].

Those crystallographic observations are reinforced by *in-solution* measurements. SEC indicates increased oligomeric heterogeneity after low-pH resolubilization (**Fig. 4a**); Raman spectra show amide I broadening compatible with increased conformational fluctuation (**Fig. 4d**); DSC reports reduced thermal stability (**Fig. 4f**); and intrinsic fluorescence exhibits a blue-shifted emission maximum (**Fig. 4h**), consistent with altered aromatic environments and hydrophobic repacking. Together, these data support a *molten*-like intermediate ensemble in which much of the secondary framework is retained while tertiary packing and hydration are loosely-coupled rearranged.

Multiscale dynamics support a directed, mechanism-level model in which GNM/ANM, TECol, diffuse-scattering prediction, and PCA all indicate that pH primarily reweights pre-existing collective modes, rather than introducing a new dynamical regime. Across methods, F1_B_, N3_B_, L6_B_, S9_B_, V12_B_, E13_B_, L16_A_, V18_B_, C19_B_, C20_A_, F24_B_-Y26_B_, and P28_B_ residues form a continuous communication axis from the allosteric N-terminus to central/C-terminal B-chain segments (**Fig. 5**). Accordingly, F1_B_ undergoes the characteristic ∼30 Å displacement associated with helical *unpeeling* [1], [54], while N3_B_ forms alternative hydrogen-bonding contacts consistent with *R^f^*-like stabilization [105], [106]. L6_B_ contributes to stabilization of Zn^2+^-anion coordination during *T*_6_→*T*_3_*R*_3_/*R*_6_ redistribution [8], and S9_B_ supports the B9-B13 interfacial hydrogen-bond network [107]. V12_B_ and L16_A_ map to the aliphatic core and receptor-relevant Site-1 region [1], whereas E13_B_ contributes to core stability through ion/water-coupled electrostatic organization that shapes the T→R barrier [60]. V18_B_ behaves as a hinge near the helix terminus [108], and the C19_B_ and C20_A_ disulfide pair supports the stress-bearing folding nucleus [109]. At the interface, P28_B_ acts as a major dimer lock controlling C-terminal detachment and subsequent hexamer/dimer dissociation [110]. Consistently, Q-Z projections place acidic states on the same pre-existing free-energy funnel, with redistributed occupancy rather than a distinct basin (**Fig. 6t**). Accordingly, phenolic ligands and protonation-dependent electrostatics may act as coupled regulators of mode selection and intermediate-state populations. Near-neutral conditions favor more compact *R*-like organization, whereas acidification weakens this stabilization and shifts occupancy toward *T*-enriched [19], [111], more exposed conformations that remain dynamically connected to the same intrinsic landscape. This helps resolve an apparent paradox in glargine behavior: precipitation and allosteric transition need not be independent events but may proceed as coupled processes in which precipitation coincides with pH-tuned redistribution of conformational substates that enable delayed dissociation (**Fig. 7**).

**Figure 7.**
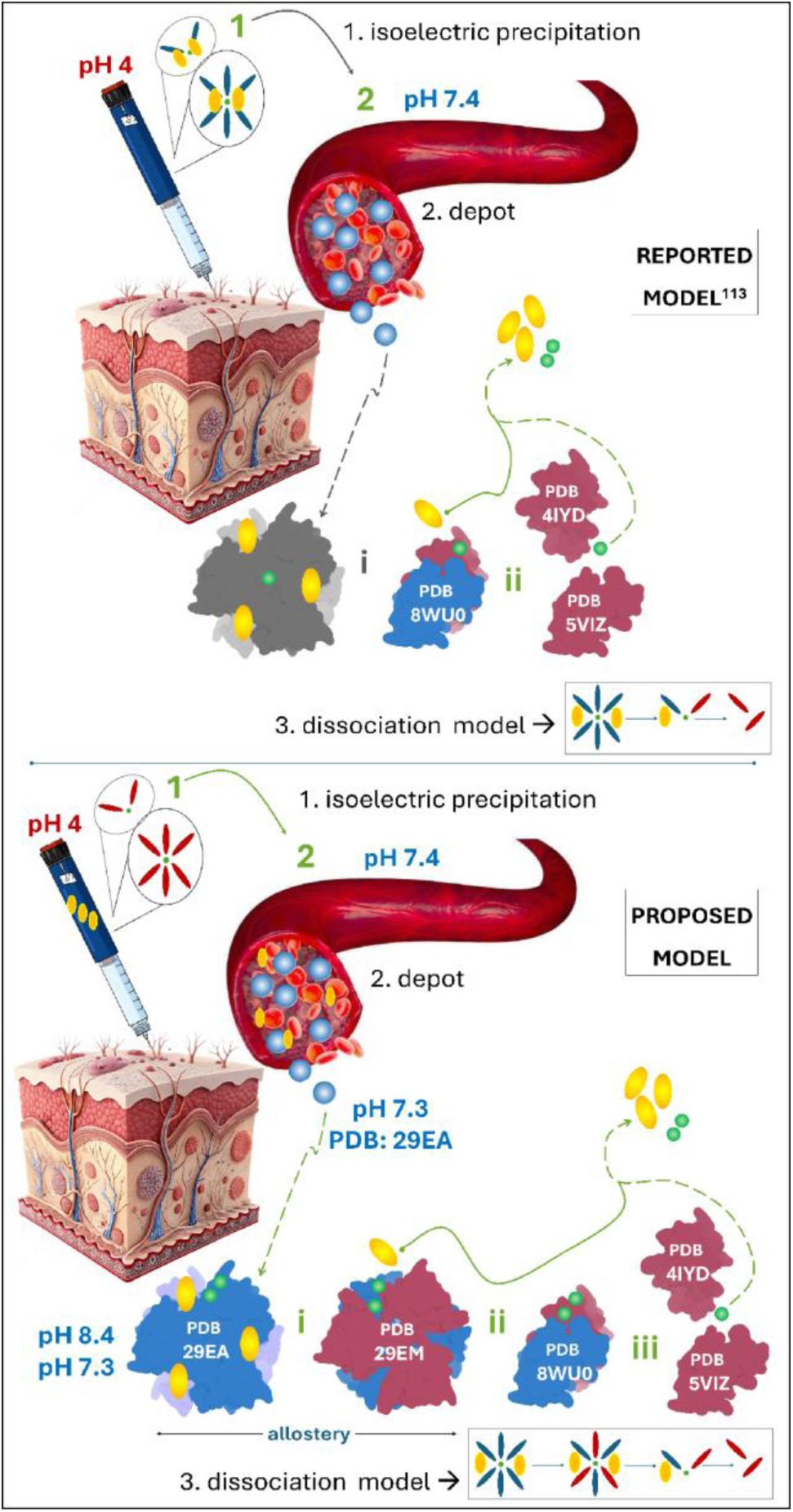
Schematic comparison of the previously reported model and the SFX-informed mechanism proposed in this study. *Top panel*: Conventional interpretation based on cryogenic PDB structures, in which Lantus (pH 4) forms a subcutaneous depot after injection (pH 7.4), primarily through isoelectric precipitation, followed by slow dissociation from hexamers to dimers, and monomers (representative PDB IDs indicated). The precise three-dimensional organization of this depot remains unresolved. *Bottom panel*: Model proposed from pH-dependent SFX structures. This study suggests that glargine hexamers/dimers [113] in the pen exist in an m-cresol-independent equilibrium while retaining R-subunits. Upon injection, isoelectric precipitation immobilizes the protein, after which the diffusible ligands (phenol, Zn²⁺) are progressively cleared by perfusion at constant near-neutral pH, driving a redistribution of hexamer conformations [116], [117]. Dissociation then proceeds through T _3_R_3_ intermediate states (molten-like state) before yielding dimers and monomers, linking precipitation to dissociation through an intrinsic allosteric transition. The scheme is a hypothesis constrained by static structures and supported by fixed-pH solution studies; arrows denote inferred population shifts, not measured kinetic steps. The lower panel represents a hypothetical model derived from in vitro structural observations; the proposed pathway and its relationship to subcutaneous release kinetics remain to be validated in vivo.

Although GNM, ANM, TECol, and PCA provide complementary insights into the proposed redistribution of pre-existing collective modes, these analyses are derived from the same four static SFX structures and therefore do not constitute fully independent evidence. Nevertheless, the robustness of the GNM-derived cross-correlation patterns to structural perturbations supports the stability of the predicted dynamical signatures. Future studies incorporating explicit conformational sampling through molecular dynamics simulations and trajectory-based network approaches, such as WISP or dCNA, could further test the proposed allosteric communication pathways and characterize the underlying conformational ensembles.

Historically, many biophysical and structural studies have shown that insulin oligomerization is highly pH-sensitive [111], [112]. Early spectroscopic and crystallographic works reported that near-neutral conditions stabilize phenol-bound *R*-state hexamers, whereas stronger acidity promotes H10_B_ protonation, zinc release, and progressive dissociation toward dimers and monomers [19], [111]. These findings established the classical model of pH-dependent insulin glargine behavior [113]. However, most prior studies captured either monomer/dimer states under cryogenic crystallographic conditions (PDB ID: 4IYD, 5VIZ), hexamer-to-monomer trends under neutral-to-acidic conditions by SAXS/CD/Raman [19], [70], [111], or highly dissociated acidic ensembles [114]. Consequently, the intermediate conformational trajectory used by basal insulin analogs across physiologically relevant pH gradients—from acidic formulations to near-neutral tissue environments—has remained insufficiently defined at atomic resolution. In particular, depot formation and dissolution have largely been interpreted as macroscopic precipitation events, with limited direct structural evidence for intermediate states [115]. The present study extends this framework by providing an ambient-temperature structural and dynamical view of pH-dependent insulin reorganization. Rather than a simple low-pH disassembly process, our data supports a redistribution of the intrinsic dynamic landscape. Across the examined pH range, the hexamer adopts compact R-state assemblies and structurally coherent TRf/T3R3-like intermediates, suggesting a potential structural link between pH-dependent conformational redistribution and controlled dissociation. These findings raise the hypothesis that environmentally regulated allosteric states may contribute to glargine’s prolonged action, although their involvement in subcutaneous depot behavior remains to be established.

Accordingly, **Figure 7** can be interpreted as a state-resolved mechanistic model that makes each release step explicit. In the previously reported model [113], Lantus in the pen (pH 4) is assumed to exist as an phenol-bound dimer/hexamer equilibrium; after injection into subcutaneous tissue (pH 7.4), depot formation is attributed to isoelectric precipitation, followed by slow dissociation over ∼24 h through hexamer to dimer to monomer states. However, the conformational identity of the depot hexamer (*R_6_* versus *T_3_R_3_*) and the three-dimensional architecture of the oligomeric depot remained unresolved [18]. Based on our pH-resolved SFX structures at ambient temperature, we proposed a more explicit pathway. We infer that the pen state (pH 4) contains a phenol-poor or phenol-free hexamer/dimer equilibrium, despite the presence of phenol in formulation, consistent with our observation that the cresol pockets are only fractionally occupied at pH 6.4 and empty at pH 5.1 (noting that the acidic titration buffers, which contain no cresol or zinc, also dilute the free ligands around the crystals (**Fig. 1j-k**), and with the established protonation of HisB10 below pH ∼5, which releases the axial zinc ions and thereby depopulates the heterotrophically coupled phenolic pockets [3]. Thus, acidic-state assemblies can retain *R*-subunit features while showing reduced phenol occupancy. After injection (pH 7.4), depot formation still occurs via isoelectric precipitation, a charge-based phenomenon governed by net charge relative to the pI and distinct from the allosteric axis described here; we propose that delayed release may involve redistribution among hexamer conformations: *R_6_*-like states observed at pH 8.4/7.3 transition through a molten-globule-like *T_3_R_3_* intermediate resolved at pH 6.4/5.1, and then proceed to dimer (8WU0) and monomer states (4IYD, 5VIZ). We emphasize that the two perturbations are mechanistically distinct: in the pen, phenol release is driven by protonation at low pH, whereas in the depot the metal site remains intact and ligand loss is driven by perfusion-mediated clearance of phenol and Zn²⁺ at constant near-neutral pH [116], [117]. Our low-pH structures are therefore presented as structural surrogates for the ligand-depleted end state rather than as a reproduction of the depot’s kinetic pathway; that partial phenol loss alone is sufficient to generate the T3R3 signature at constant neutral pH is established in solution [117], [118]. A time-resolved experiment at fixed, near-neutral pH with controlled ligand depletion (for example serial crystallography on precipitated glargine during dialysis against ligand-free buffer) would provide the direct test and is the natural next step of this work. Collectively, this model proposes a potential connection between precipitation and dissociation through intrinsic allosteric redistribution, rather than establishing a causal pathway. Importantly, our in vitro structural and biophysical measurements do not directly establish conformational states or dissociation kinetics within the subcutaneous depot. In vivo or ex vivo validation will be required to determine whether the proposed allosteric intermediates contribute to glargine’s prolonged pharmacokinetic profile.

## 4. Methods

### 4.1 Experimental design for crystallographic analysis

#### 4.1.1 Glargine sample preparation and crystallization

Commercially available insulin glargine (Lantus®; pH 4.0), one of the most widely used basal insulin formulations, was crystallized using a sitting-drop micro-batch under-oil vapor-diffusion strategy in 72-well Terasaki^TM^ plates (Greiner-Bio, Germany). Briefly, glargine solution was mixed 1:1 (v/v) with commercially available sparse-matrix screening conditions at room temperature, covering approximately 3,500 conditions like our previous insulin studies [60], [112]. Each well contained 0.83 uL total protein/condition mixture and was overlaid with 16.6 uL paraffin oil (Tekkim Kimya, Turkiye), then incubated at room temperature until harvesting [33]. Crystal formation was monitored by light microscopy, and microcrystals appeared within 1-2 h (**Fig. S2a**). Productive conditions were scaled in Wizard^TM^-1 condition #23 containing 15% (v/v) reagent alcohol, 200 mM MgCl2, and 100 mM imidazole/HCl (pH 8.7). For large-scale preparation, 12.5 mL of crystal suspension was mixed directly with 12.5 mL of the same buffer, yielding immediate microcrystal formation (**Fig. S1**). The final crystal-suspension pH was 8.4, and stocks were maintained at room temperature until use.

#### 4.1.2 Crystalline preparation at various pH

Crystal density was increased to approximately 10^8^ crystals mL^−1^ by centrifuging 500 μL slurry (3000 rpm, 5 min), removing 490 μL supernatant, and retaining 10 μL concentrated slurry for injection. To generate different pH conditions, concentrated crystals (10 μL) were mixed with defined buffer volumes before grease embedding. For pH 8.4, crystals were mixed with 90 μL of Super Lube nuclear-grade grease (No. 42150, Synco Chemical Co.) on a plastic plate, following Sugahara et al. [119]. For pH 7.3, 6.4, and 5.1, 10 μL of slurry was mixed with 5 μL of the corresponding buffer and 85 μL of grease (**Fig. S3**). For intermediate pH values (6.7, 6.2, and 5.5), 10 μL of slurry was mixed with 2 to 4 μL of pH 5.1 buffer and 86 to 88 μL of grease (**Fig. S3**). All preparations were completed ∼10 min before data collection, and the final slurry pH was verified using a micro-pH electrode. Samples were mixed thoroughly with grease (No. 42150, Synco Chemical; **Fig. S4_1-4_**) and loaded bubble-free into a 200 μL cartridge (**Fig. S4_5-6_**). The cartridge was mounted on a high-viscosity cartridge-type (HVC) injector (**Fig. S4_7-10_**) equipped with a 75 μm inner-diameter nozzle (**Fig. S4_11_**) and installed in the He-filled, coaxial gas-assisted, suction-stabilized DAPHNIS chamber (**Fig. S4_12-13_**). The HPLC flow rate was 0.0127 μL min^−1^, and the grease-embedded crystal stream was injected at 0.79 μL min^−1^. The replaceable cartridge reservoir enabled rapid sample exchange and, together with a temperature-controlled cooling jacket, minimized abrupt temperature fluctuations during data collection.

#### 4.1.3 Processing of diffraction images and structure refinement

Diffraction data from insulin glargine microcrystals were collected at Experimental Hutch 3 of SACLA BL2 using 10 keV XFEL pulses (<10 fs, 30 Hz), with the MPCCD detector positioned at a 50 mm sample-to-detector distance (**Fig. S5**). Diffraction images were identified from raw diffraction frames using `peakfinder8` in *Cheetah* [120]. Selected images were then indexed in *CrystFEL* v0.9.1 [121] with `xgandalf` (min-snr = 4.5, threshold = 100, minimum pixel count = 2). Indexed patterns were scaled and merged in *CrystFEL* using `partialator` with `--model=unity` to obtain structure-factor amplitudes. Data quality metrics were calculated with `comparehkl` and `checkhkl` (**Table 2** and **Table S1**). For the pH 5.1 and pH 6.4 datasets, indexing ambiguity was detected using `cell_tool` [121] and resolved by applying the ambigator operator `-h,-h-k,-l` to both datasets. Our SFX structures were determined by automated molecular replacement in *Phaser*, yielding space group P12_1_1 for pH 8.4 (PDB ID: 29DZ) pH 7.3 (PDB ID: 29EA) along with R3:H space group for pH 6.4 (PDB ID: 29EN), pH 5.1 (PDB ID: 29EM). Previously reported cryogenic models (PDB ID: 1XDA for P12_1_1; PDB ID: 8WU0 for R3:H) were used for initial rigid-body refinement. After simulated-annealing refinement, individual coordinates and TLS parameters were further refined, and solvent modeling was curated in *Coot* [122]: sites with strong difference density were retained, whereas waters outside significant electron density were removed. Data-collection statistics are summarized in **Table 2** and **Table S1**, and structural figures were prepared in *PyMOL* [123]. In a final round, zinc-site coordination was restrained (Zn–NE2 2.03 Å, Zn–Cl 2.25 Å; phenix.metal_coordination), the occupancies of Zn²⁺ and Cl⁻ were refined, the axial ions of the R3:H models were constrained to the crystallographic 3-fold, and single side-chain atoms left at zero occupancy by earlier rounds were restored. Analysis of the R3:H data with phenix.xtriage indicated merohedral twinning by the operator h,−h−k,−l (⟨I²⟩/⟨I⟩² = 1.57–1.59, ⟨|L|⟩ = 0.37, multivariate L-test Z = 13.9–14.1; R_twin = 0.07 in the unresolved merges), consistent with residual indexing ambiguity and/or twin domains generated by the crystallographic P12₁1 → R3:H conversion; the pH 6.4 and 5.1 structures were therefore refined against the twinned data with this twin law: pH 6.4 in phenix.refine (least-squares twin target, TLS; twin fraction 0.23; R-work/R-free 0.216/0.271) and pH 5.1 in *Servalcat* (maximum-likelihood twin target with jelly-body restraints; twin fraction 0.45; R-work/R-free 0.230/0.277, against 0.264/0.377 without the twin law), the latter because the least-squares twin target over-fitted these weaker data (R-work/R-free 0.199/0.278). Zn²⁺ and Cl⁻ occupancies were refined in phenix.refine with the twin law and held fixed in *Servalcat*. The P12₁1 data showed no twinning. Final statistics are given in **Table 2**.

**Table 2.**
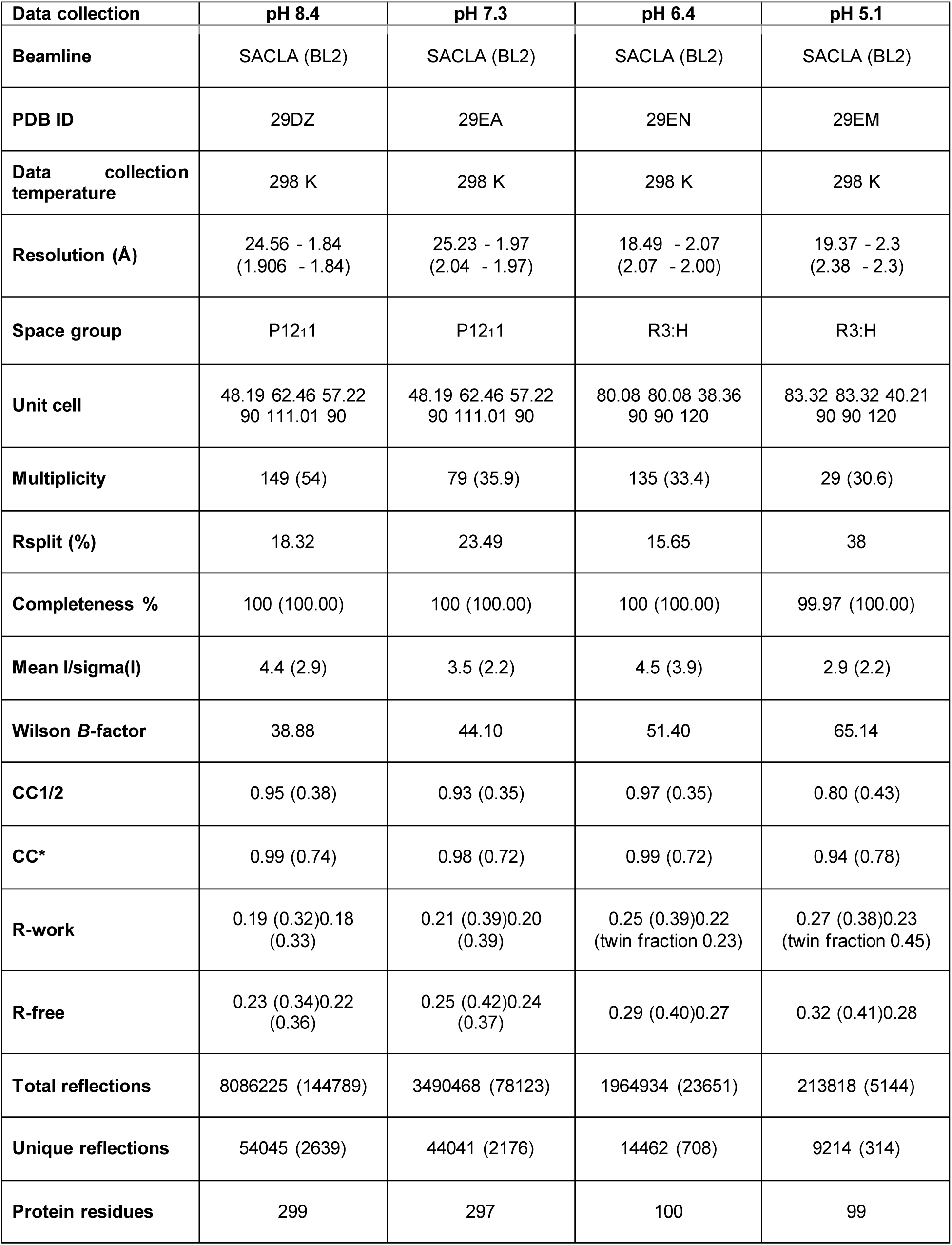
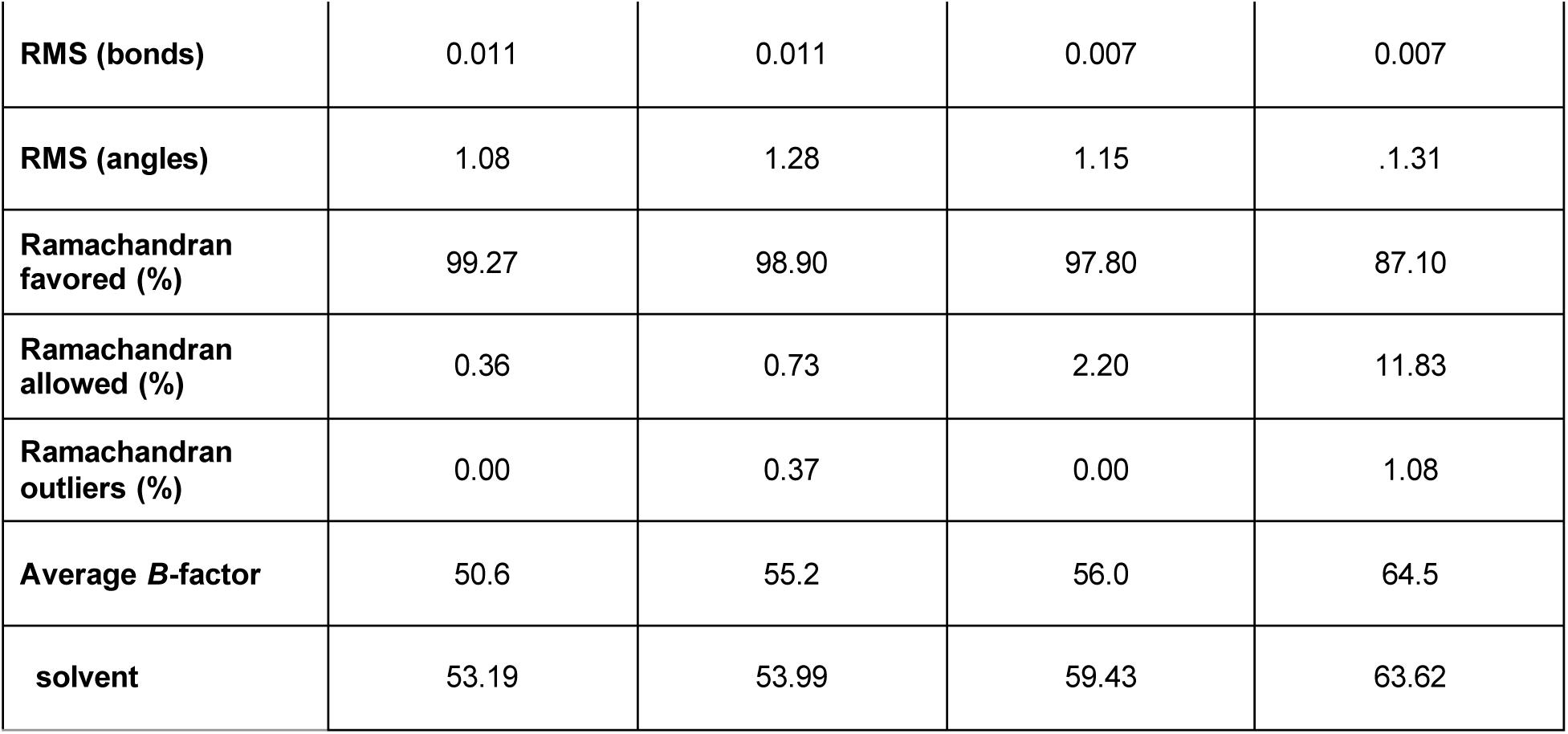
Data collection and refinement statistics.

#### 4.1.4 Difference electron-density maps

Difference maps were calculated with the matchmaps strategy using *gemmi* and *reciprocalspaceship*. CrystFEL intensities were Friedel-merged and converted to amplitudes by the French–Wilson method to 1.8 Å (⟨I/σ(I)⟩ ≈ 1 at 1.85–2.0 Å). Each perturbed dataset was scaled to the reference amplitudes with an isotropic k·exp(−Bs²/4) model (B = −7.5, −18.5 and −23 Å² for pH 7.3, 6.4 and 5.1 relative to pH 8.4). For the isomorphous pair (pH 7.3 vs 8.4) the pH 8.4 model was rigid-body refined against each dataset; for the rhombohedral datasets the pH 8.4 dimer was placed in the R3:H cell by aligning the hexamer 3-fold with the crystallographic 3-fold and scanning the rotation about it (R = 0.40 versus a scan median of 0.47), followed by rigid-body refinement, with the remaining dimers related by NCS superposition. Observed maps were phased with the bulk-solvent-corrected model of each dataset at 2.0 Å (1.9 Å for pH 7.3), the perturbed map was resampled into the reference frame through the refined-model transforms and subtracted within 5 Å of the model after scaling on the density at atom centres, and the difference was expressed in units of its standard deviation. For the twinned R3:H data the amplitudes were twin-corrected before map calculation, algebraically for pH 6.4 (twin fraction 0.23) and by partitioning each observed intensity between the twin-related reflections in proportion to the calculated intensities of the twin-refined model for pH 5.1 (twin fraction 0.45), after which the placement discriminated the two axial orientations of the hexamer (R = 0.42 versus 0.46). Maps were also computed with the pH 7.3 and pH 6.4 structures as references. Peaks ≥ 3σ within 2.5 Å of a model atom were tabulated per residue and counted as features only when present in at least half of the copies of that residue in the hexamer. Omit maps (B1–B8 of both chains, and the cresol position transplanted from the pH 7.3 model) were calculated from the twin-refined R3:H models with the same software, with the twin correction recomputed from each omit model.

### 4.2 Experimental design for in-solution analysis

#### 4.2.1 Size-exclusion chromatography (SEC) analysis

SEC analysis was performed on glargine crystals at pH 8.4 and 5.1, on a glargine sample (Lantus, pH 4) adjusted to pH 5.1 as a cross-check, and on a glargine sample adjusted to pH 2.7 as a monomer control. A Superdex 200 (S200) prep grade was used for the pH 8.4 and 5.1 conditions with SEC buffers containing 150 mM NaCl and 20 mM Tris-HCl, adjusted to the target pH (8.4 or 5.1) using 20 mM citric acid. For pH 2.7 measurements, 20 mM citric acid was used directly as the SEC buffer. Eluted fractions corresponding to each peak were collected and analyzed by 20% SDS-PAGE (**Fig. 4a; Fig. S15**).

#### 4.2.2 Raman spectroscopy

High-resolution in situ micro-Raman spectroscopy was performed to analyze pH-dependent chemical and structural changes in insulin crystals at pH 8.4, 7.3, 6.4, and 5.1 using a Thermo Scientific DXR Raman microscope (Thermo Fisher Scientific). Measurements were acquired with a 532 nm (green) diode-pumped solid-state (DPSS) Nd:YAG laser at 2 mW, using a 50× objective in confocal mode (50 μm confocal aperture/slit). For each measurement, the acquisition time was 4 min with a spectral resolution of ∼4 cm^−1^; point spacing was 10 μm, and the total measurement time was 10 min. Before crystal measurements, Raman spectra of the corresponding pH buffers were collected as controls. To correct for buffer contributions, each buffer control spectrum was subtracted from the Raman spectrum of crystals at the same pH. Raman intensities were then normalized to the highest peak in each spectrum. Difference spectra were generated by subtracting spectra at pH 7.3, 6.4, and 5.1 from the pH 8.4 crystal spectrum, followed by normalization to the maximum intensity of each difference spectrum. Data analysis was performed using Origin 2022b software.

#### 4.2.3 Differential scanning calorimetry

Differential scanning calorimetry (DSC) was performed using insulin glargine samples obtained from Lantus® SoloStar® pens (100 U/mL, Sanofi) and a SETARAM Micro DSC-III calorimeter. Samples were prepared independently of those used for SEC, without prior crystallization or crystal resolubilization. The samples were adjusted to pH 8.4, 7.3, 6.4, or 5.1 using 20 mM citric acid and diluted to a final protein concentration of 2.3 mg/mL. Measurements were acquired over a temperature range of 20-100 °C with a heating rate of 0.3 K/min. To minimize vessel heat-capacity correction, the pen insulin sample and reference buffer (15% (v/v) reagent alcohol, 200 mM MgCl2, and 100 mM imidazole/HCl) were mass-balanced to ±0.05 mg. A second thermal scan of the denatured sample was recorded and used for baseline correction. Melting temperatures (T_m_) and calorimetric enthalpies (ΔH_cal_) were analyzed using Origin® 2021 (OriginLab Corporation, Northampton, USA).

#### 4.2.4 Steady-state fluorescence emission

Intrinsic fluorescence emission from tyrosine and phenylalanine side chains was monitored using a HORIBA Jobin Yvon Fluorolog 3.22 spectrofluorometer. Measurements were performed with pen insulin solution samples (Lantus® SoloStar® 100 U/mL, Sanofi) at a protein concentration of 2.3 mg/mL under pH 8.4, 7.3, 6.4, and 5.1 conditions. Tyr and Phe residues were excited at 280 nm, and emission spectra were collected from 300 to 400 nm using 5/5 nm excitation/emission slit settings at 20 °C. Spectral emission data were evaluated using Origin 2022b software.

### 4.3 Computational design for in-silico analysis

#### 4.3.1 Gaussian network modeling and transfer entropy-collectivity analysis

GNM analysis was carried out using the ProDy package [124] on four newly determined SFX glargine structures spanning pH 8.4 to pH 5.1, together with their monomeric and dimeric components (including isolated monomers and dimers), previously available P12_1_1 and R3:H structures, and additional glargine structures available in the PDB. This framework is based on construction and eigenvalue decomposition of the Kirchhoff matrix Γ (N x N, where N is the total number of residues), which encodes network connectivity. The cutoff distance for long-range interactions was set to 8.0 Å. GNM yields N-1 nonzero normal modes, with mode frequencies and shapes defined by the corresponding eigenvalues and eigenvectors of Γ. Cross-correlations of equilibrium residue motions were obtained from the pseudoinverse of Γ and organized in covariance matrix C, where diagonal elements represent residue mean-square fluctuations and off-diagonal elements represent pairwise cross-correlations. Normalization of these cross-correlations by residue mean-square fluctuations yields orientational cross-correlations. Matrix C can also be expressed as a sum of contributions from individual modes derived from eigenvalue decomposition of Γ (or C). The heat maps shown in this study (**Fig. 5a**) correspond to normalized cross-correlations, also referred to as correlation cosines, between residue pairs contributed by the softest modes. Inverse eigenvalues of Γ were used as modal weights; relative modal contributions were quantified from fractional contributions of individual modes, and cumulative contribution or variance for mode subsets was computed by summation. Soft modes at the lowest-frequency end of the spectrum were selected to provide a minimum cumulative variance of 0.40, whereas fast motions were represented by the 10 highest-frequency modes at the opposite end of the spectrum.

GNM-TE integrates Transfer Entropy (TE) with GNM to quantify directional information flow between residues *i* and *j* [36]. TE measures the reduction in uncertainty in the motion of residue *j* given the motion of residue *i*, including an explicit time delay *τ*, denoted as *Ti→j(τ)*. Net TE is defined as the directional difference between *Ti→j(τ)* and *Tj→i(τ)*. For each residue *i*, the TECol score was computed by multiplying its cumulative positive net TE (the sum of positive net TE values transmitted from residue *i* to other residues) by its collectivity value *K_i,s_*within each subset ‘*s’*.

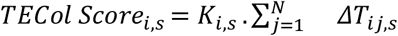

For each subset ‘*s*’, collectivity term *K_i,s_* is computed from the chosen slow-mode set

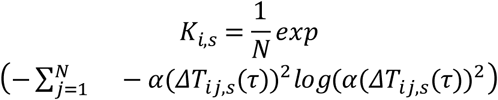

where ‘*s*’ is the selected subset of slow GNM modes, *N* is the total residue count, and alpha is the normalization constant [36], [125]. By combining directional transfer (TE) with collectivity (*K*), the TECol score provides a robust criterion for identifying the most functionally significant residues as global information sources.

#### 4.3.2 Predicted diffuse scattering analysis

To estimate the contribution of structural disorder and correlated atomic motions within the SFX glargine crystal dataset under the same pH conditions to the diffuse X-ray scattering [126], the Gaussian Approximation [88], [127] was employed. The acquisition of theoretical diffuse scattering maps and intensity distributions was performed in three main stages;

***(i)*** *set-up of dynamic correlations via GNM*: To describe the intrinsic dynamics of the protein backbone and interatomic correlations, our previous GNM method was adjusted for predicted diffuse scattering calculation. To reduce computational cost and focus on main-chain fluctuations, only alpha-carbon (*Cα*) atoms were included in the analysis. Likewise, the cutoff distance of 8.0 Å was used to determine interatomic interactions, and by calculating the slowest modes of the system, the dynamic cross-correlation matrix (*ρ_ij_*) between *Cα* atoms was obtained,
***(ii)*** *derivation the atomic displacement covariance matrix (V_ij_):* The covariance matrix of atomic displacements (*V_ij_*), which is the fundamental factor determining the diffuse scattering intensity, was constructed by combining the experimental *B*-factors with the correlations obtained from the GNM. First, the mean-square displacement (⟨*u*_*i*^2 ⟩) of each *Cα* atom was calculated from the experimental *B*-factors using the equation ⟨*u*_*i*^2 ⟩ = *B*_*i*/(8*π*^2). Under the assumption of isotropic motion, the diagonal elements (individual variances) of the covariance matrix were defined as *V*_*ii* = ⟨*u*_*i*^2 ⟩/3. The off-diagonal elements (interatomic shared variances, *V_ij_*) were derived by scaling the cross-correlation coefficients (*ρ_ij_*) obtained from the GNM with the displacement magnitudes of the respective atoms using the formula,

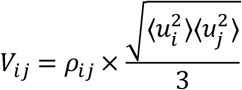
***(iii)*** *calculation of diffuse scattering intensity (I_diffuse_*(*q*)*) via Gaussian Approximation* [88], [89], [128]: After constructing the covariance matrix, the diffuse scattering intensity *I_diffuse_*(*q*) was calculated using the fundamental formula, which assumes that atomic displacements follow a Gaussian distribution [88], [128], [129]:

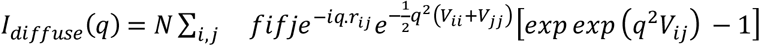 Here, *q* is the 3-dimensional scattering vector space defining such that *q_max_* = 2.5 Å^−1^. *f_i_* and *f_j_* are the atomic form factors for the *Cα* atoms, assumed to be simplified and constant (approximately 6 *e*^−^). *r_ij_* is the distance vector between atoms *i* and *j*. The first exponential term (damp) in the equation represents the Debye-Waller factor which suppresses Bragg diffraction, while the term inside the square brackets determines the contribution of interatomic correlations to the scattering. The computations were solved over complex number matrices for each vector on the grid created in *q*-space, and the scattering maps were generated by taking the real parts of the results,
***(iv)*** *data analysis and visualization*: The 3-dimensional *I_diffuse_*(*q*) values calculated for each pH state were projected onto the *q*-vector space as 2-dimensional slices for visualization purposes. To perform statistical analysis of the distribution and to observe the intensity shifts under different pH conditions, frequency histograms of the logarithmic scattering intensities (*log*_10_(*Intensity*)) were constructed (over *q*-vector counts).

#### 4.3.2 Mode-of-variation analysis

Mode-of-variation analysis was performed as described in our previous report [112]. Briefly, atomic coordinates from the four refined SFX glargine structures (pH 8.4-5.1) were extracted, and each model was represented by *Cα* atoms only. To ensure comparability, all structures were standardized to the same length by truncating to the minimum shared *Cα* count (*N*), retaining the first *N* residues in each model. Principal component (PC) analysis was then applied to aligned, truncated *Cα* coordinates to capture dominant structural variation. For each structure, x/y/z coordinates were concatenated into a single vector, producing a data matrix *X* of size *M*✕3*N* (*M* = number of structures). The analysis followed standard steps: ***(i)*** mean-centering of coordinate dimensions, ***(ii)*** covariance matrix computation, ***(iii)*** eigen decomposition to obtain principal components (eigenvectors) and their explained variances (eigenvalues), and ***(iv)*** projection of centered coordinates onto these components to generate PC scores. Explained variance for each component was calculated as its eigenvalue divided by the sum of all eigenvalues and reported as a percentage. Structural separation was visualized in the PC1-PC2 plane, with points labeled by pH. To localize residue-level effects, selected *Cα* atoms (defined by residue number and chain ID) were extracted and examined for their contributions to the PC.

#### 4.3.3 Free energy folding funnel analysis

To characterize the topological order and global collapse dynamics of the insulin structures across different pH states, a multidimensional folding funnel model was employed [91], [130]. The conformational space map was defined using two fundamental order parameters: the fraction of specific native contacts (*Q*), which represents the degree of topological folding (nativeness), and the total number of contacts (*Z*), which measures the overall tendency for spatial collapse [92]. Calculations were performed using the *ProDy* library [124]. For each conformation, the distance matrices between *Cα* atoms were computed, and an interatomic contact cutoff of 8.0 Å was applied. The *Q* and *Z* values were evaluated relative to the reference native-like structure, defining the position of each conformation within the phase space [91]. To translate the discrete distribution of these structures in the (*Q*,*Z*) space into a continuous thermodynamic landscape, a two-dimensional Gaussian Kernel Density Estimation (2D KDE) was applied. Using the resulting conformational probability density, *P*(*Q*,*Z*), the entropy (*S*) and relative free energy (*F*) surfaces of the system were derived based on the fundamental Boltzmann relationships [59]:

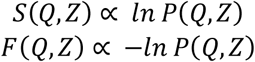

The resulting free energy matrix was shifted such that the global minimum was set to zero, and the landscape was visualized as both 2D iso-energy contour maps and a 3D free energy funnel. This approach provided a macroscopic topographical map of the folding funnel, elucidating the thermodynamic barriers and intermediate structural traps within the system [130].

## 5. Conclusion

This study provides an ambient temperature structural view of insulin glargine across a physiologically relevant pH range and provides a structural framework for investigating the molecular basis of prolonged action. Rather than supporting a simple acid-driven dissociation model, our data indicates a coordinated conformational redistribution between compact *R*-state hexamers and more plastic, yet structurally coherent, *TR_f_/T_3_R_3_*-states (**Movie S2**). This interpretation is further supported by convergent evidence from SFX structures, in-solution spectroscopy and calorimetry, and multiscale dynamic analyses. Importantly, the observed pH-dependent lattice and conformational allostery suggest that protracted action is related to reorganized intermediate conformers (**Figure 7**), not to random depot dissociation. Although isoelectric precipitation remains central to depot formation, we hypothesize that subsequent release may involve redistribution among allosteric hexamer states, including molten-globule-like T3R3 intermediates, before further dissociation into smaller oligomers. Thus, precipitation and dissociation may be mechanistically linked, a possibility that requires direct in vivo or ex vivo validation. Although the in vivo three-dimensional architecture of the depot remains unresolved, the state-resolved pathway defined here offers a practical basis for future experimental testing. Collectively, these findings provide a framework for biosimilar comparability assessment and for future basal insulin design efforts by focusing on intermediate-state stability and allosteric plasticity as tunable determinants of release behavior, while more broadly, the data suggest that environmentally triggered protein release can be mediated by redistribution among pre-existing allosteric substates rather than by simple two-state switching.

## Author Contributions

The hexamer insulin analog was crystallized by EA. In-crystallo follow-up experiments at various pH conditions were carried out by EA. Sample preparation for SFX data collection was performed by EA under the supervision of JK, TY, and TT. SFX data for glargine crystals at different pH conditions were collected at SACLA XFEL by EA and JK under the supervision of TT and MY. Crystal data processing was conducted by EA and MKS. Structural refinement and further interpretation were performed by EA. Sample preparation and Raman spectroscopy experiments were performed by EA, ET, and KF, while size exclusion chromatography, differential scanning calorimetry and fluorescence emission analyses were performed by EA and ET. GNM, GNM-TECol, predicted diffuse scattering, mode-of-variation, and free-energy folding funnel analyses were conducted by EA using custom scripts developed based on the relevant equations reported in the literature and in consultation with field experts mentioned in the *acknowledgements*.

## Acknowledgements

This work was supported by the SACLA Research Support Program for Graduate Students (proposals 2023B8058 and 2023B2761) and the Koc University Travel Support Program. The authors thank Taito Osaka, Kohei Miyanishi, Gota Yamaguchi, and Tetsuya Ishikawa for their support during XFEL data collection. The authors also sincerely thank Thomas A. White for his invaluable assistance with data processing and for confirming the unit-cell and space-group transition associated with pH-dependent changes. This work benefited from training received at the International School of Crystallography (IsoC, Erice 2022); EA and MKS gratefully acknowledge Michael E. Wall, David Wych, and especially Ariana Peck for valuable assistance in interpreting the diffuse-scattering analyses. E.A. also thanks Ivet Bahar, Hoang Nguyen, and Turkan Haliloglu for insightful discussions on GNM analyses, and Abdullah Kepceoglu for support with PCA calculations. The authors further thank the Laboratory for Structural Biology at Koc University and the University of Health Sciences for preparation of the glargine crystalline samples.

## Data availability

Atomic coordinates and structure-factor data have been deposited in the RCSB Protein Data Bank under accession codes 29DZ (pH 8.4), 29EA (pH 7.3), 29EN (pH 6.4), 29EM (pH 5.1). Additional information is available from the corresponding author upon reasonable request.

## Supplementary File

**Figure S1.**
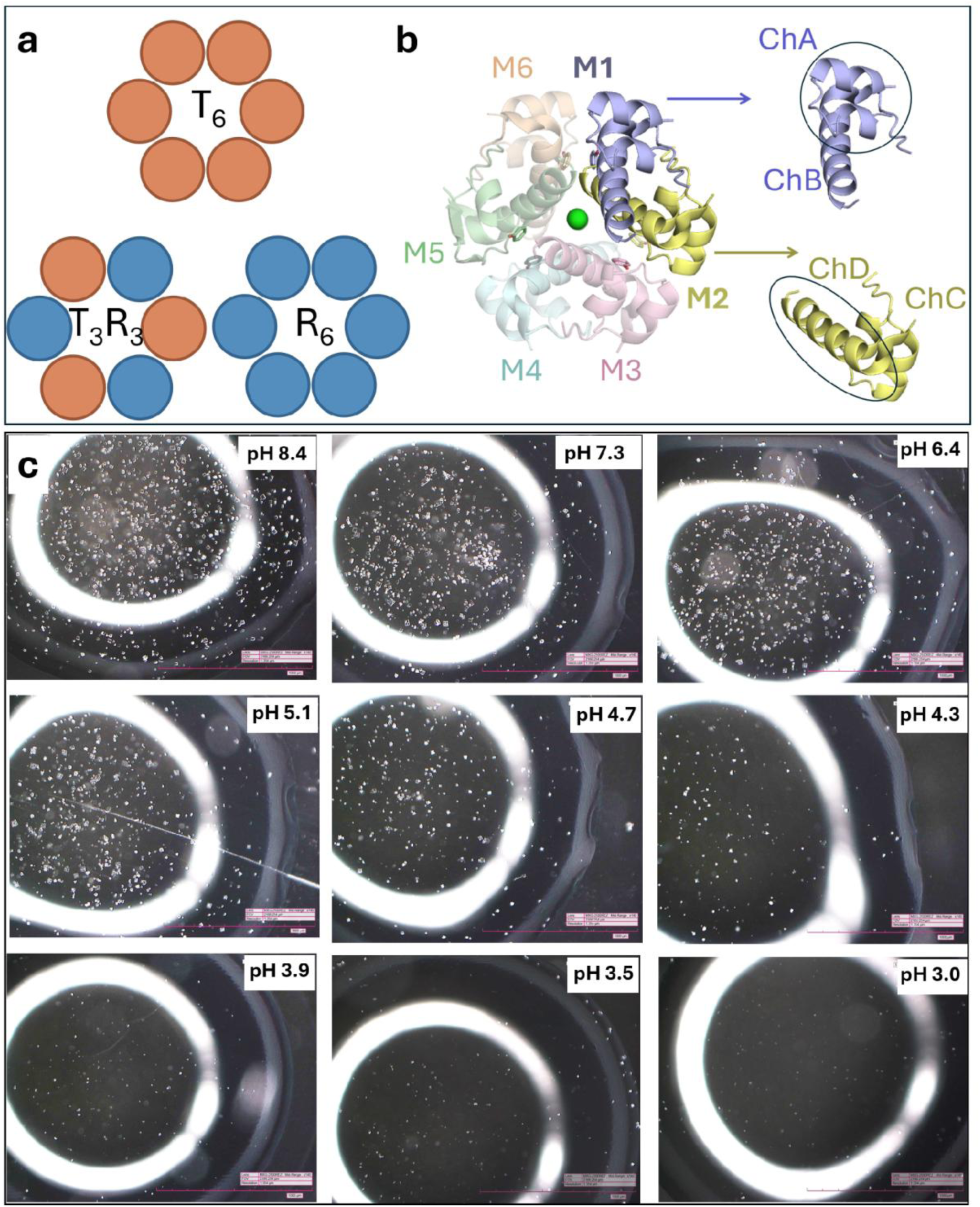
**a,** Schematic representation of the three classical insulin hexamer conformational states, T_6_, T_3_R_3_, R_6_ containing six T-state protomers, three T-state plus three R-state protomers, and six R-state protomers, respectively. **b,** Whole-hexamer representation of insulin glargine, showing six individually colored protomers (M1–M6), each comprising an A and a B chain. Representative protomers are enlarged to illustrate the constituent chains and their corresponding chain identifiers (ChA–ChD)**. c,** pH screening was performed on the crystals within the pH range of 8.4 to 3.0. A gradual disappearance of the monoclinic crystals was observed at lower pH values, likely due to lattice disorder. Diffraction data were successfully collected between pH 8.4 and 5.1. However, a notable increase in background noise was detected as the pH decreased, and no usable data could be obtained below pH 5.1.

**Figure S2.**
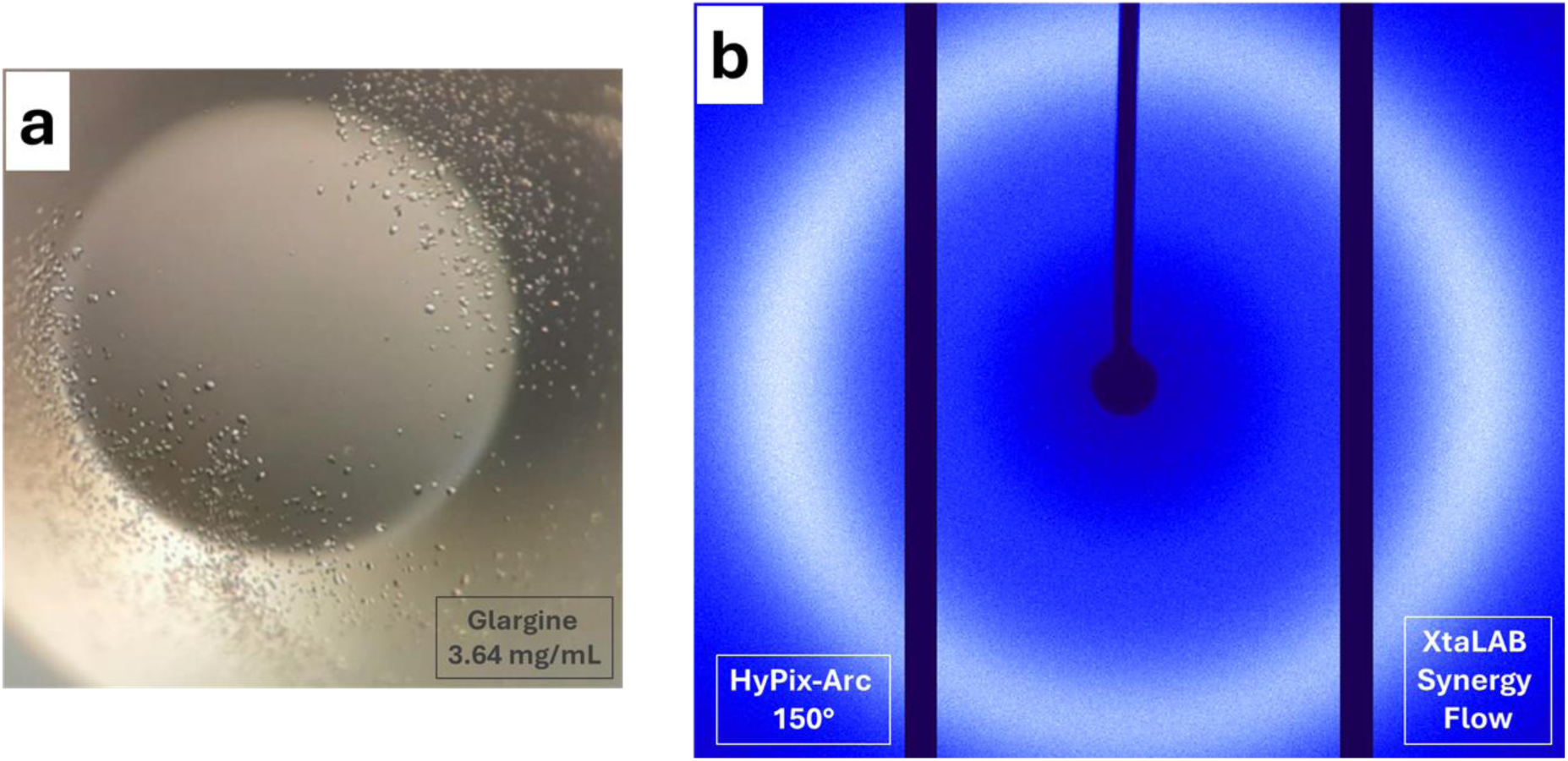
Microcrystalline insulin glargine fails to diffract at a home X-ray source. a, Optical micrograph of insulin glargine microcrystals formed in Terasaki plates at a protein concentration of 3.64 mg mL⁻¹. The crystals are extremely small and monoclinic in morphology. b, Representative diffraction image collected using a home X-ray source in plate-reader mode (Rigaku *XtaLAB* Synergy Flow), showing the absence of measurable Bragg diffraction due to the submicron size of the crystals.

**Figure S3.**
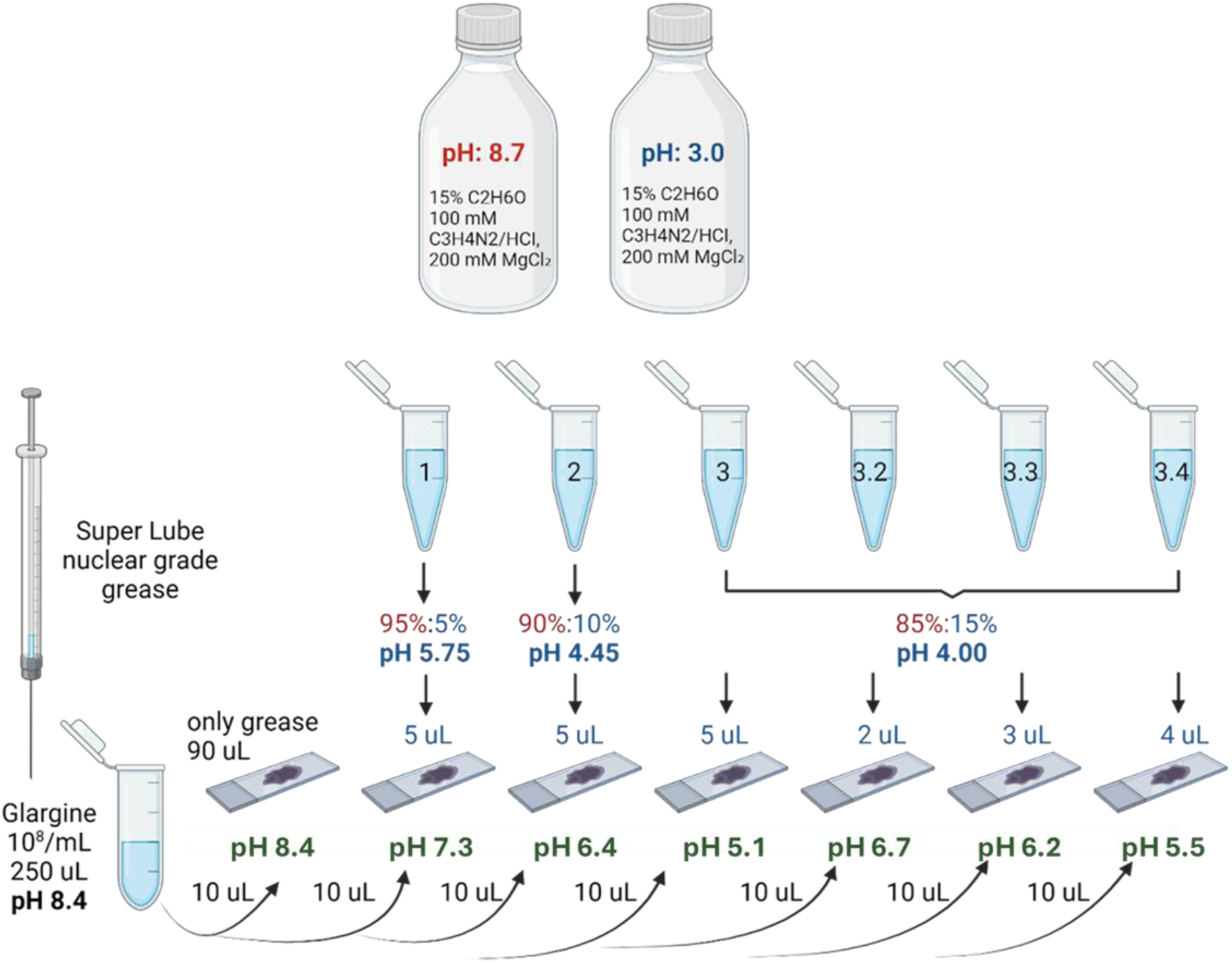
Schematic representation of the pH screening procedure for glargine crystals embedded in a grease matrix. Two crystallization buffer solutions were prepared at pH 8.7 and pH 3.0, each containing 15% PEG 600, 100 mM C₃H₈N₂/HCl, and 200 mM MgCl₂. Intermediate pH values (5.75, 4.45, and 4.00) were obtained by mixing the two crystal buffers at defined volume ratios as illustrated. These intermediate buffers were further diluted to prepare additional conditions ranging from pH 7.3 to pH 5.1 by adding 5 μL of buffer to the grease matrix, which was then mixed with 10 μL of crystal (Lantus glargine, pH 4 + crystal buffer, pH 8.7) samples. pH values ranging from 6.7 to 5.5 have been obtained by mixing the pH 4.00 crystal buffer solution at various volumes (2, 3, and 4 μL) with a grease-crystal mixture (90 μL:10 μL). A suspension of glargine crystals (10¹⁰/mL, 250 µL, pH 8.4) alone was mixed with 90 µL of Super Lube nuclear-grade grease. All pH values have been confirmed by a micro pH meter.

**Figure S4.**
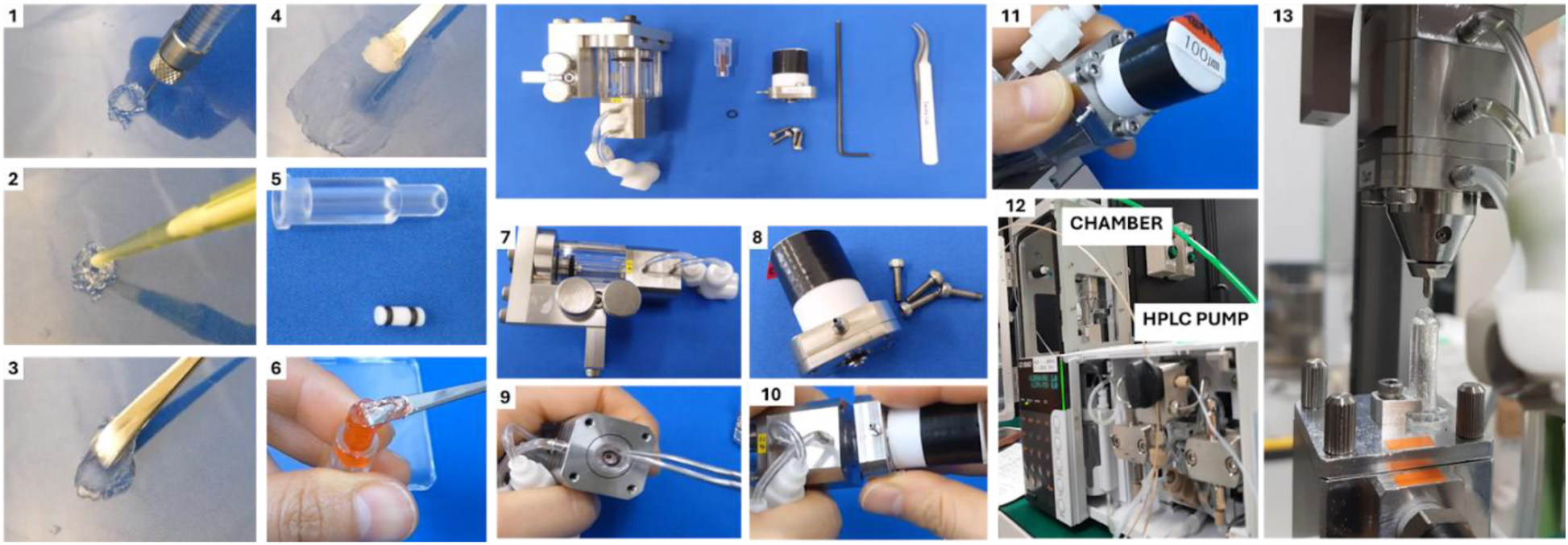
Sample preparation steps throughout the high-viscosity cartridge-type (HVC) injector. (1) A total of 100 µL of grease matrix was prepared using a microinjector. (2-3) The sample was subsequently mixed with the grease matrix and thoroughly homogenized to ensure a uniform distribution within the medium. Once a well-mixed sample was obtained (4), it was carefully loaded into a 200 µL cartridge pre-assembled with a Teflon insert and O-rings (5-6) to maintain airtightness and compatibility with the injection system. The setup process continued with the preparation and installation of the scaffolding screws required for the High-Viscosity Cartridge (HVC) injector (7-10). Following this, (11) a protective nozzle was mounted onto the injector to safeguard the system during operation. (12) The experimental chamber and High-Performance Liquid Chromatography (HPLC) pump were then configured to support the HVC injection process. (13) The HVC injector was inserted into the DAPHNIS system and securely positioned onto the suction device, completing the setup for high-viscosity sample delivery.

**Figure S5.**
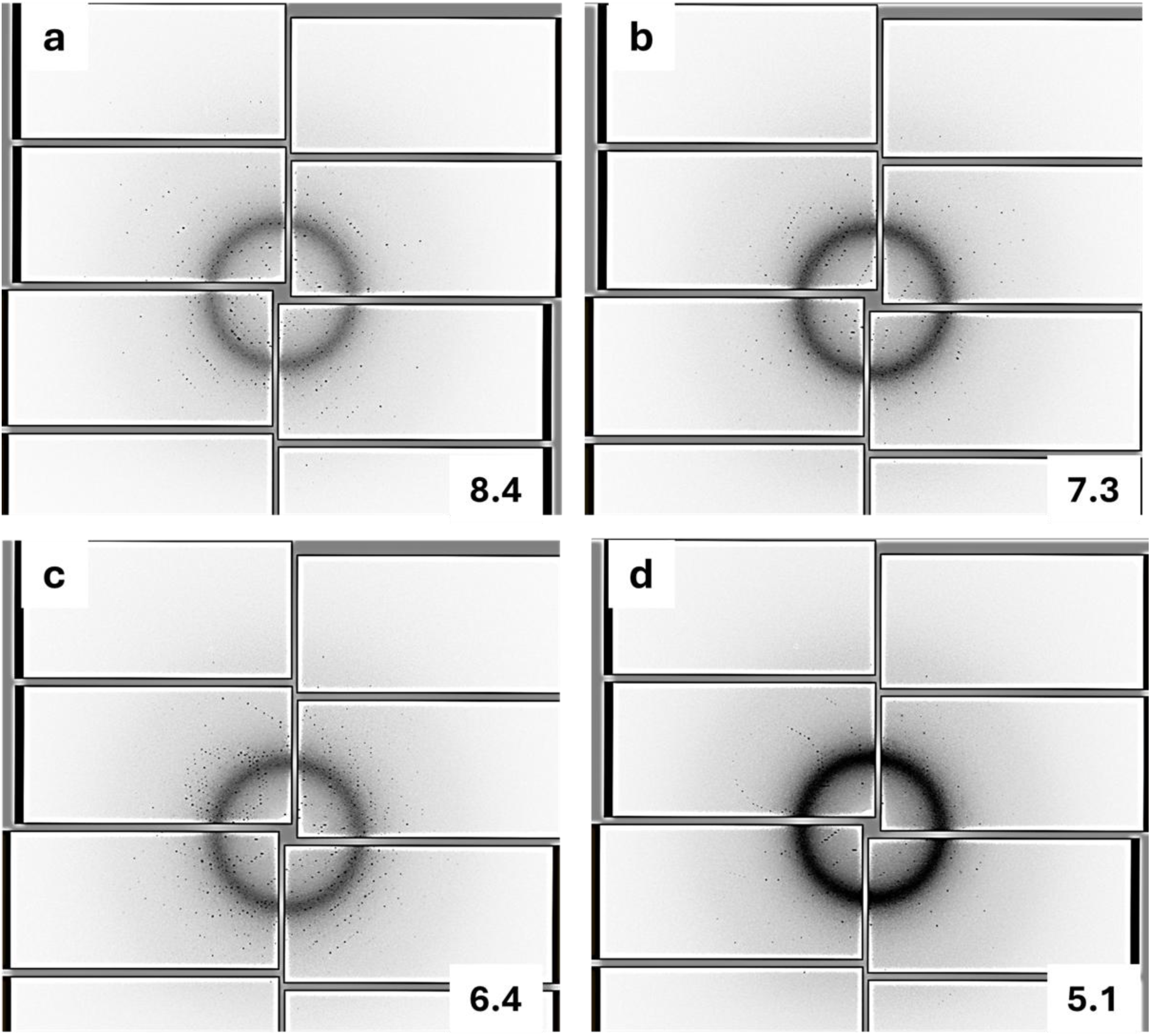
pH-dependent changes in SFX diffraction signatures of insulin glargine microcrystals. Representative diffraction patterns obtained at a, pH 8.4; b, pH 7.3; c, pH 6.4; and d, pH 5.1 using HVC-mediated SFX at SACLA. Progressive alterations in diffraction ring position and intensity distribution indicate pH-driven changes in lattice symmetry and interplanar spacing.

**Figure S6.**
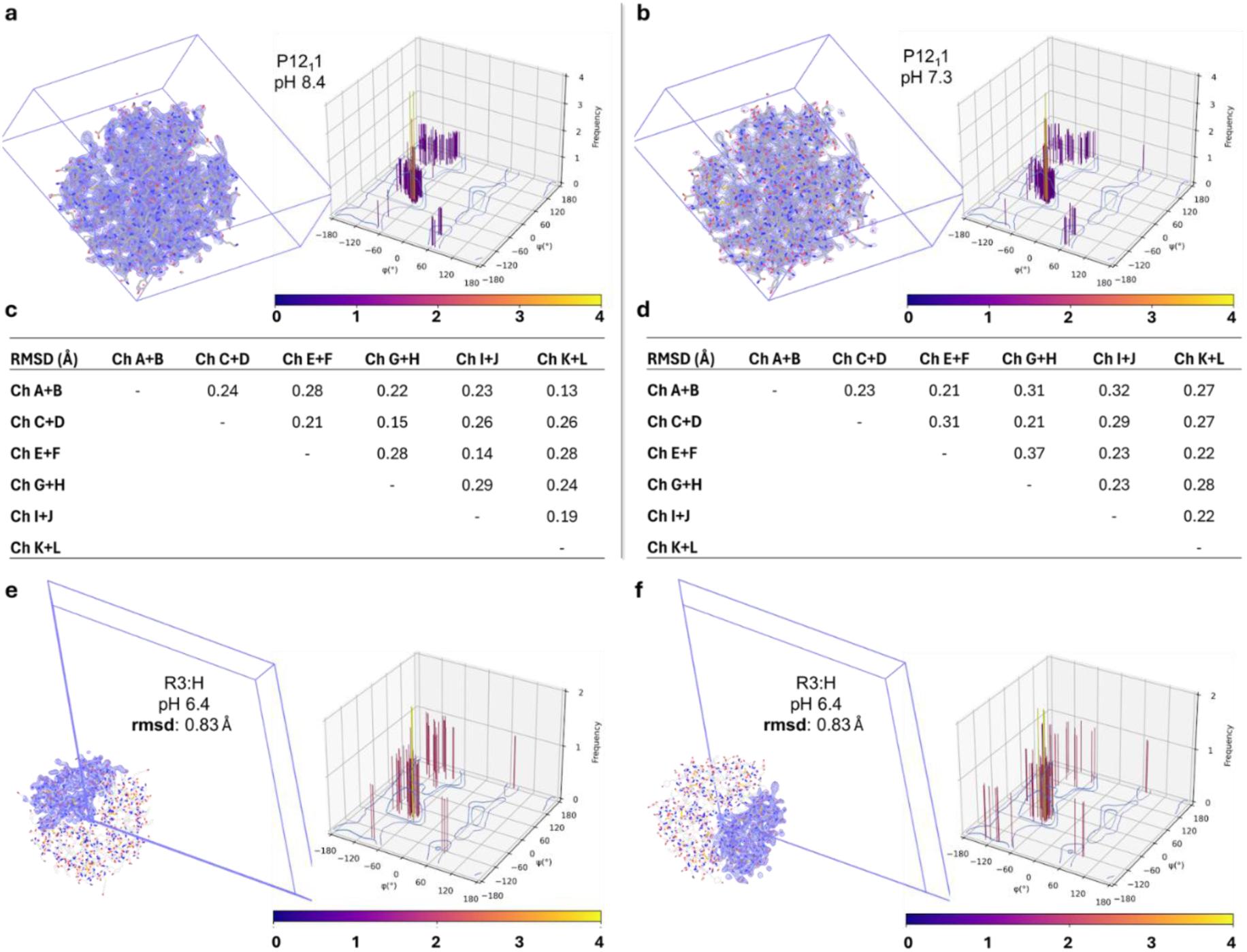
pH-dependent lattice and conformational allostery of hexameric insulin glargine revealed by 3D Ramachandran analysis and inter-monomer RMSD. (a, b) At pH 8.4 (a) and pH 7.3 (b), hexameric insulin glargine crystallizes in the monoclinic P12_1_1 space group. Three-dimensional Ramachandran plots generated via *RamPlot* display backbone torsion angles (ϕ,ψ), where the vertical bars (z-axis) and color gradient represent the frequency of occurrence. The distributions are tightly clustered with tall, sharp peaks (yellow–orange) confined strictly to the favored α-helical basin defined by the Top8000 reference dataset. This indicates a conformationally restricted, rigid backbone in which residues consistently populate the thermodynamic minimum, while widespread but low-frequency purple–blue signals reflect negligible conformational excursions. (c, d) Pairwise RMSD matrices between symmetry -related dimers at pH 8.4 (c) and pH 7.3 (d) reveal minimal structural deviations across the hexamer, consistent with the focused Ramachandran distributions and a uniform *R^f^_6_* state. (e, f) At pH 6.4 (e) and more prominently at pH 5.1 (f), the plots reveal a marked redistribution of backbone sampling, characterized by the coexistence of high-frequency bars across multiple (ϕ,ψ) regions rather than confinement to a single dominant basin. Although locally high-frequency (favored) peaks persist, they are fragmented and interspersed with widespread low-to-moderate-frequency sampling (purple–red) that extends into allowed and generously allowed regions. This multi-modal landscape reflects the increased conformational heterogeneity and enhanced backbone plasticity associated with the *T_3_R^f^_3_* state. The effect is most pronounced at pH 5.1, where the broadened torsional distribution, together with elevated global RMSD values (∼0.83 Å), supports cooperative and dynamically reconfigurable rearrangements within the hexamer. The 2mFₒ–DFc electron-density maps are contoured at 1.0σ and carved within 1.6 Å of the displayed atomic models.

**Figure S7.**
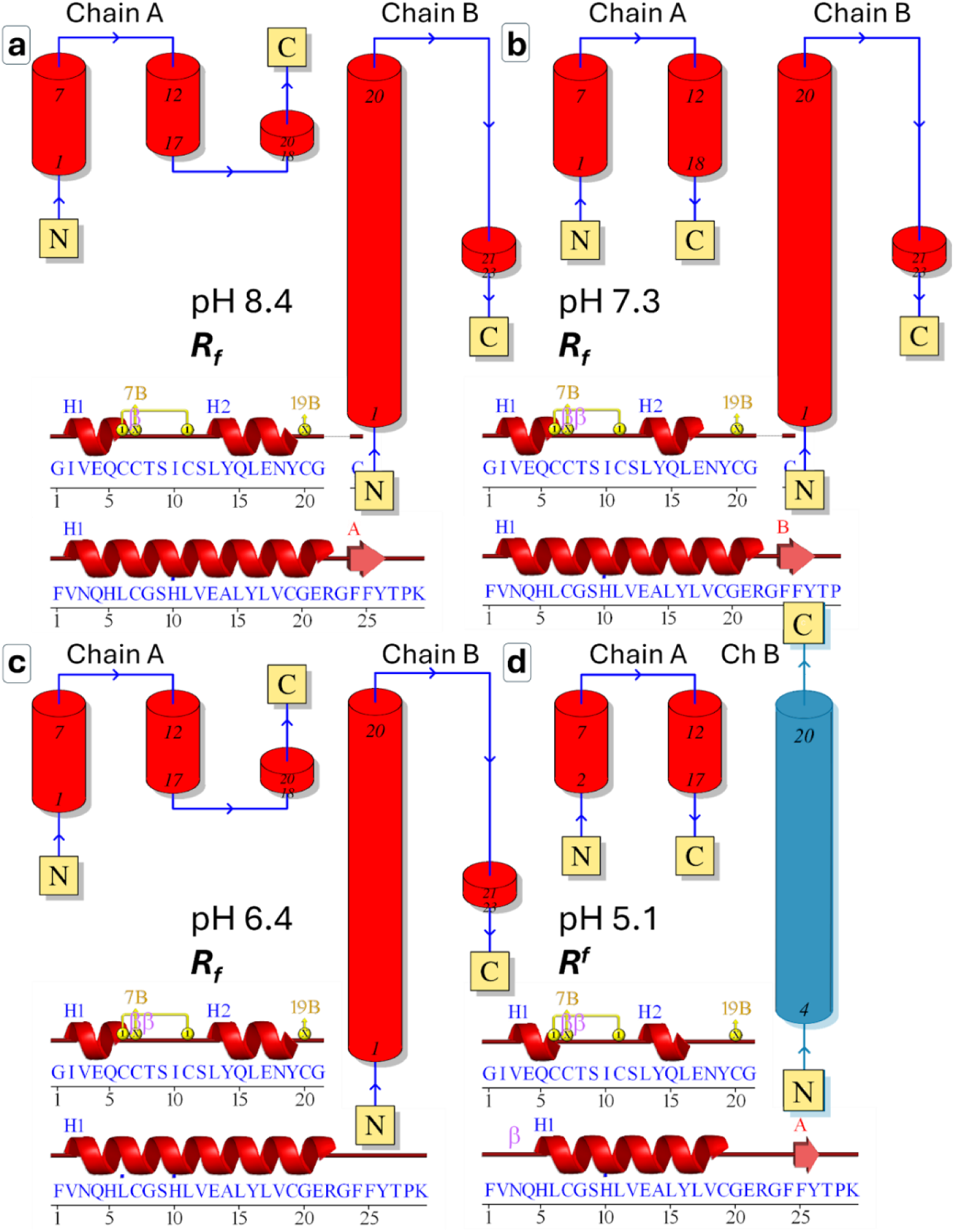
pH-dependent reorganization of secondary-structure topology in insulin glargine monomers. Schematic representation of the secondary-structure architecture of insulin glargine at different pH conditions, highlighting chain-specific and pH-dependent changes in helix length, continuity, and topology. a,b, At pH 8.4 (a) and pH 7.3 (b), both chains adopt the frayed-relaxed (*Rᶠ*) conformation, characterized by continuous helical segments along the B chain (B1–B19) and a conserved helical topology in chain A. c, At pH 6.4, the *Rᶠ* topology is retained; however, subtle shortening and reorganization of helices are observed, indicating the onset of pH-dependent structural asymmetry. d, At pH 5.1, pronounced topological remodeling occurs, with chain-specific alterations in helix length and continuity, reflecting a collapse of the higher-pH helical architecture into a reorganized configuration (blue). Although classified as *Rᶠ* based on backbone topology, the pH 5.1 *Rᶠ* subunit occupies a distinct dynamic regime, shaped by protonation-driven electrostatic remodeling and mechanical coupling to neighboring *T* subunits. Cylinders denote α-helices, arrows indicate chain directionality, and residue numbers correspond to helical boundaries derived from the refined crystallographic models

**Figure S8.**
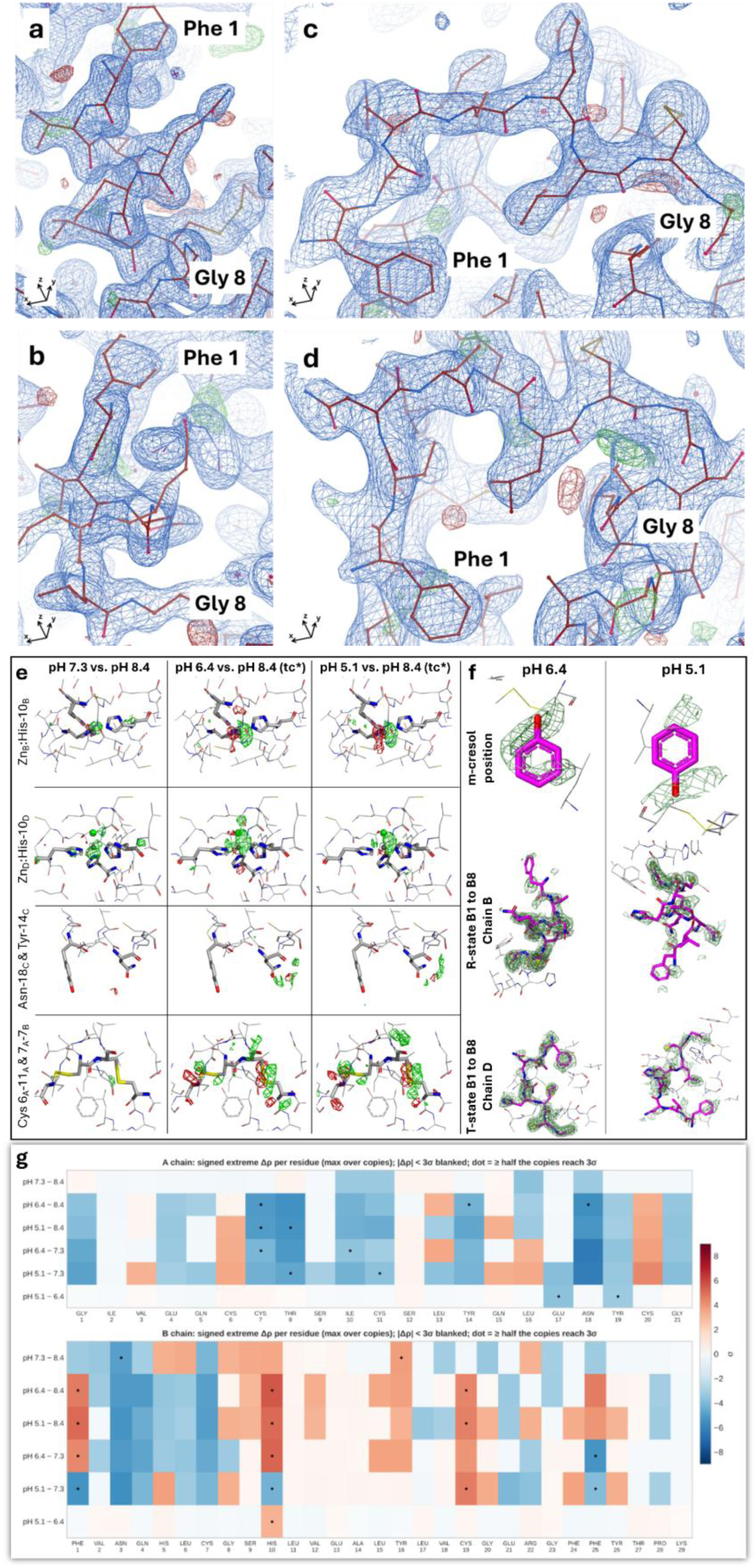
2Fo-Fc and omit maps for pH-dependent *R^f^*–*T* allosteric remodeling at the B-chain N-terminus. **a,** Representative *2Fo-Fc* electron-density maps contoured at 1.0σ (carved within 1.6 Å) illustrate the conformational states of the B-chain N-terminal segment encompassing F1_B_ and G8_B_. Panels **a,b** show the *R^f^* state, characterized by a continuous, well-ordered helical trajectory with tightly confined electron density. Panels **c,d** depict the *T* state, where the density broadens and redistributes, consistent with backbone extension and increased conformational plasticity. Atomic models are shown as sticks, with density maps rendered in blue (2*Fo–Fc*), and difference features in green/red (*Fo–Fc*). **e,** Difference density relative to pH 8.4 dataset, green shows +3.5σ and red indicates −3.5σ; carved 3 Å; Zn in grey color; Cl in green color. **f,** Twin-aware omit Fo-Fc of phenol indicated progressive diminished of the omit density toward lower pHs. Shifting from R to T state in Chain D of lower pHs has been supported by difference density, Plane green color shows +1.5σ; dark green color indicates +2.5σ; omitted atoms in magenta color.

**Figure S9.**
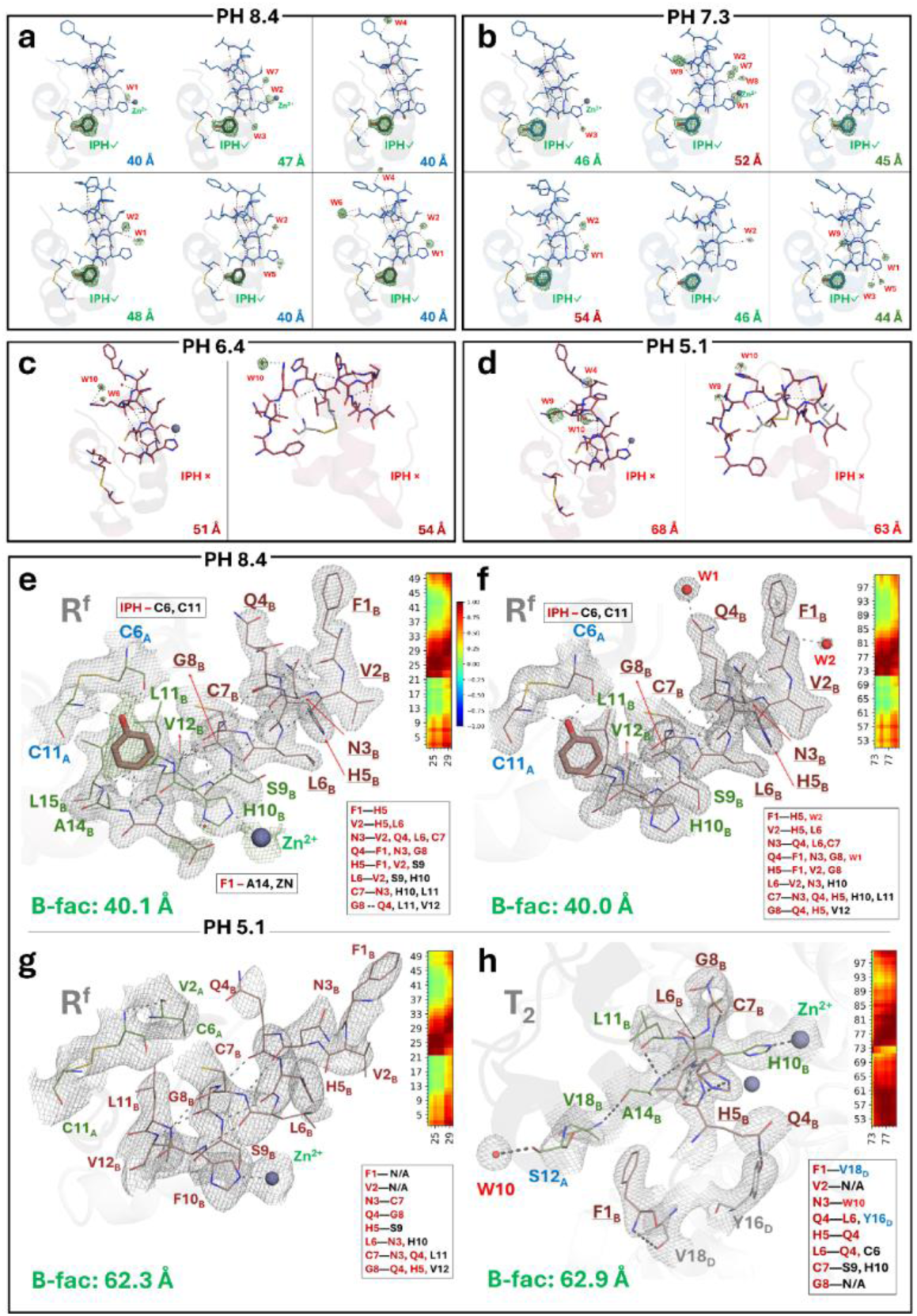
Electron density analysis of our SFX structures reveals pH-dependent reorganization of phenol-binding, solvation, and allosteric networks within the *R^f^_6_* state. a,b, At pH 8.4 and 7.3, phenol molecules and structured water molecules bound to the *R^f^₆* hexamer are shown with their corresponding 2*Fo–Fc* electron-density maps contoured at 1.0σ (carved within 1.6 Å), confirming well-ordered ligand and solvent occupancy within the hydrophobic pockets of *R^f^*-state subunits. c,d, At pH 6.4 and 5.1, phenol binding is absent; however, structured water molecules remain clearly resolved in electron density, occupying positions vacated by phenol and supporting a reorganized solvent network upon acidification. e,f, At pH 8.4, the first two monomers of the hexamer (*R^f^₆*) are shown, highlighting the allosteric B-chain N-terminal segment (B1–B8) and its interacting residues. All side chains involved in intra-monomer contacts are displayed as sticks and are fully supported by electron density, revealing an extensive interaction network within the allosteric region. The allosteric residues harboring this network were subsequently analyzed using GNM cross-correlation heatmaps, which showed moderate collectivity, consistent with moderate *B*-factors (∼40 Å²). g,h, At pH 5.1, the first two monomers of the *T₃R^f^₃* hexamer—corresponding to one *R^f^*-state (*R^f^₃*) and one *T*-state (*T₃*) subunit—are shown. In the *R^f^*-state monomer, the B1–B8 allosteric region engages in fewer interactions, while in the *T*-state monomer, these interactions are further reduced. All observed contacts (or their loss) are explicitly supported by electron density, excluding model bias. GNM cross-correlation analyses of the first monomer (*R^f^*-status) reveal diminished intra-monomer coupling in agreement with the elevated B-factors (∼62Å²), while the second monomer (*T*-status) shows the strongest cooperativity compared to all *R^f^*-state monomers, inverse correlation with the elevated B-factors (∼63 Å²), and indicates enhanced collective plasticity at low pHs. GNM cross-correlation heatmaps use a common color scale from −1 (anti-correlated) to +1 (positively correlated), with 0 indicating no correlation.

**Figure S10.**
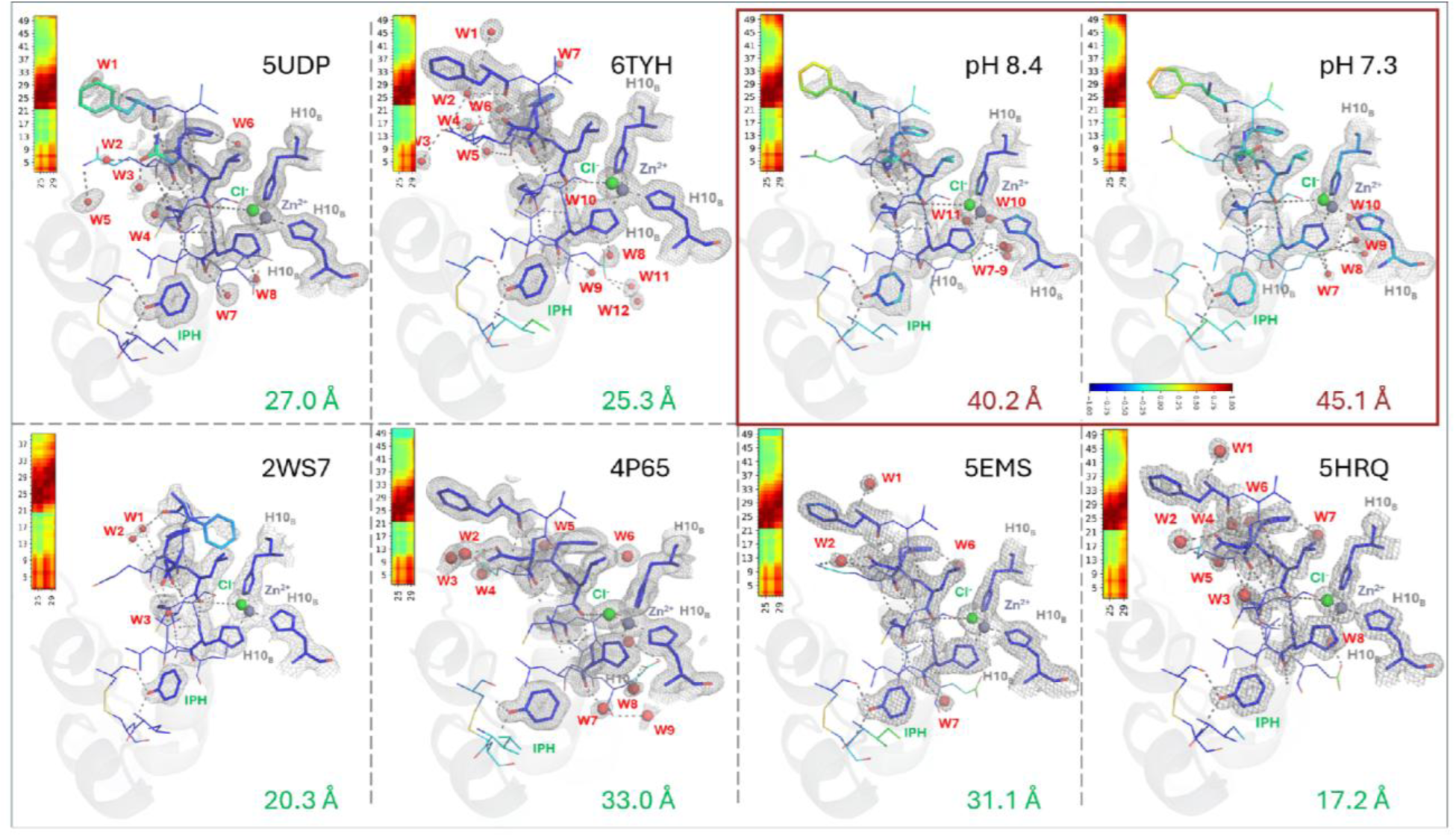
Structural divergence in *R^f^_6_* subunits of monoclinic structures: a comparative analysis of phenol, structured water, and allosteric networks between our ambient SFX structures (red box) versus PDB structures. All residues, phenol molecules (IPH), Zn²⁺ ions, anions, and structured water molecules are shown as sticks and validated by 2*Fo-Fc* electron density maps contoured at 1.0σ (carved within 1.6 Å). Archival PDB structures display compact phenol-binding pockets stabilized by dense structured-water networks and extensive local hydrogen bonding, consistent with lower *B*-factors (≈17–33 Å²) and a rigidified *R^f^₆* conformation. In contrast, the SFX-derived structures exhibit fewer structured water molecules, partially rewired hydrogen-bond networks, and substantially elevated *B*-factors (∼40–45 Å²), indicating a dynamically stabilized *R^f^₆* state under near-physiological conditions (298 K). Despite overall similarity in global fold and Zn²⁺ coordination geometry, cross-correlation analysis by GNM reveals a striking dynamical distinction: while showing a similar cooperative profile across all structures, in the SFX-derived *R^f^₆* subunits, strong long-range dynamical coupling between the allosteric B-chain N-terminus (B1–B8) and the distal B-chain C-terminal region (residues ∼45–50) is uniquely displayed. Overall, this correlated motion is absent in all examined PDB *R^f^₆* structures, in which allosteric dynamics remain locally confined. GNM cross-correlation heatmaps use a common color scale from −1 (anti-correlated) to +1 (positively correlated), with 0 indicating no correlation.

**Figure S11.**
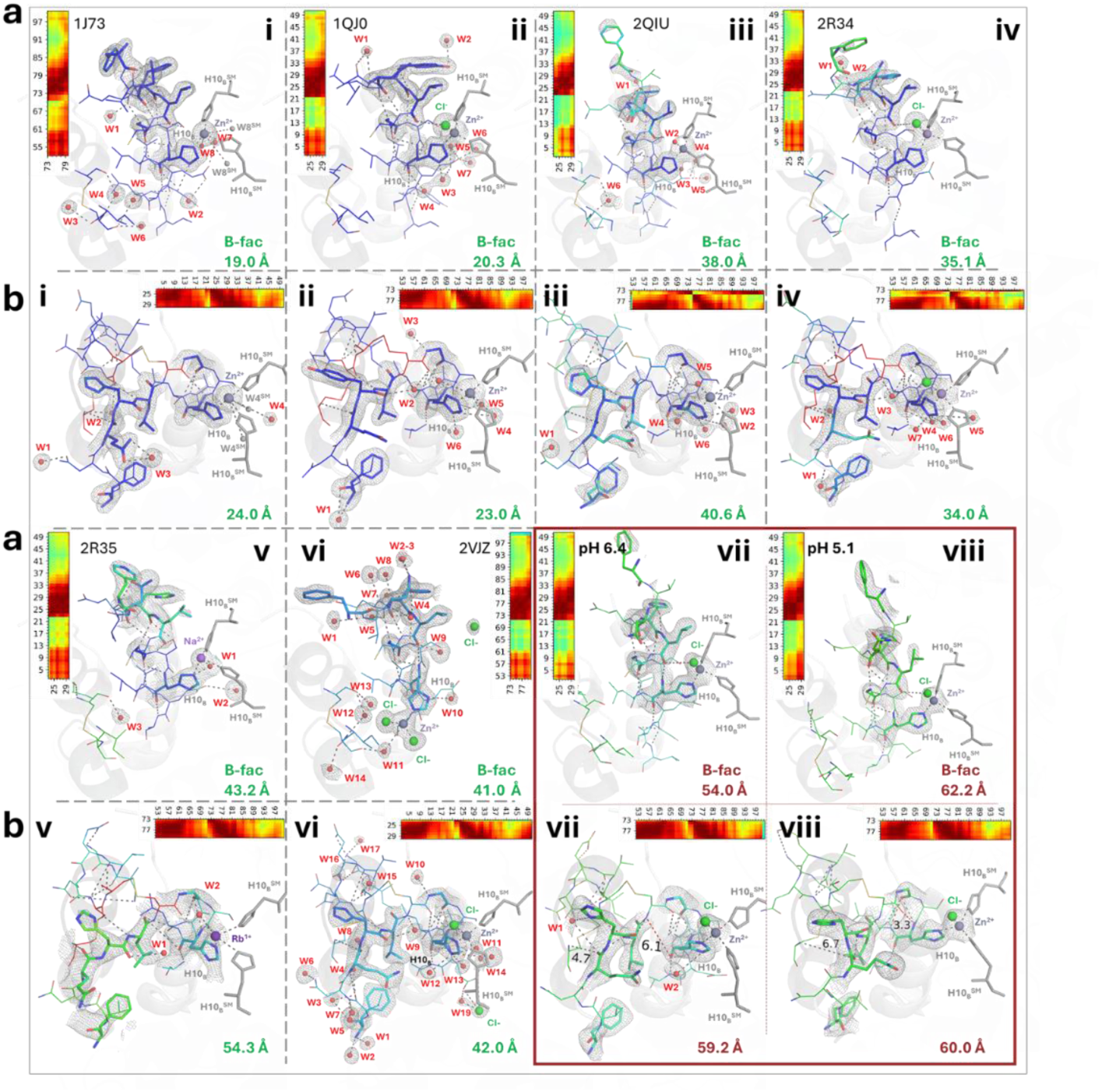
Structural divergence in *T_3_R^f^_3_*subunits of rhombohedral structures: a comparative analysis of structured water and allosteric networks in the absence of the phenol, between our ambient SFX structures (red box) versus PDB structures. Comparative visualization of the allosteric network in phenol-free *T₃Rᶠ₃* glargine hexamers across eight rhombohedral structures (i–viii), including archival PDB models (1J73, 1QJ0, 2QIU, 2R34, 2R35, 2VJZ) and our ambient-temperature SFX structures at pH 6.4 and pH 5.1 (red boxes). a, *R^f^-*subunits (i–viii). For each structure, the *R*-state (*Rᶠ*) monomer is shown with the allosteric B-chain N-terminus and its interaction partners rendered as sticks; structured water molecules (W labels) and relevant ions are included. All displayed features are supported by 2*Fo-Fc* electron density contoured at 1.0σ level (carved within 1.6 Å). b, *T*-subunits (i-viii). Equivalent representation for the *T*-state monomer (sticks + waters/ions + 1.0*σ* density), highlighting pH-dependent rewiring of allosteric contacts in the absence of phenol. For each *R/T* subunit, a GNM cross-correlation heatmap (appended at the panel edge) reports residue–residue dynamical coupling of the allosteric region with adjacent residues within the same monomer. Notably, in the *R* subunits (panels a_i-viii_), the long-range coupling between the allosteric B-chain N-terminus and the distal B-chain C-terminus (residues ∼45–50)—prominent in the *R^f^₆* subunits of our SFX structures (*see, Fig. S10*)—is not observed in any *R^f^*-subunits of *T₃Rᶠ₃* conformers (including our pHs 6.4 and 5.1) with the exception of cooperativ e motion (∼1.00) in 1J73 and 1QJ0. In the *T*-subunits (panels b_i-viii_), archival structures typically exhibit pronounc ed solvent coordination and metal/ion geometry (*eg, octahedral coordination in 1J73_T_ and 1QJ0_T_; tetrahedral coordination in 2QIU_T_ (Zn²⁺ present), and 2R34_T_ & 2R35_T_ (Mn^2+^ and Rb⁺, respectively*). In contrast, our tetrahedrally coordinated *T*-subunits (pH 6.4 and 5.1) show marked depletion of structured waters yet elevated *B*-factors, consistent with increased conformational plasticity under near-physiological conditions. Strikingly, despite reduced solvent stabilization and higher *B*-factors, the GNM cross-correlation profile of the *T*-state remains broadly conserved, as in archival PDB models, indicating that collective allosteric dynamics can be maintained through contact rewiring even when the canonical hydration/coordination spine is absent. GNM cross-correlation heatmaps use a common color scale from −1 (anti-correlated) to +1 (positively correlated), with 0 indicating no correlation

**Figure S12.**
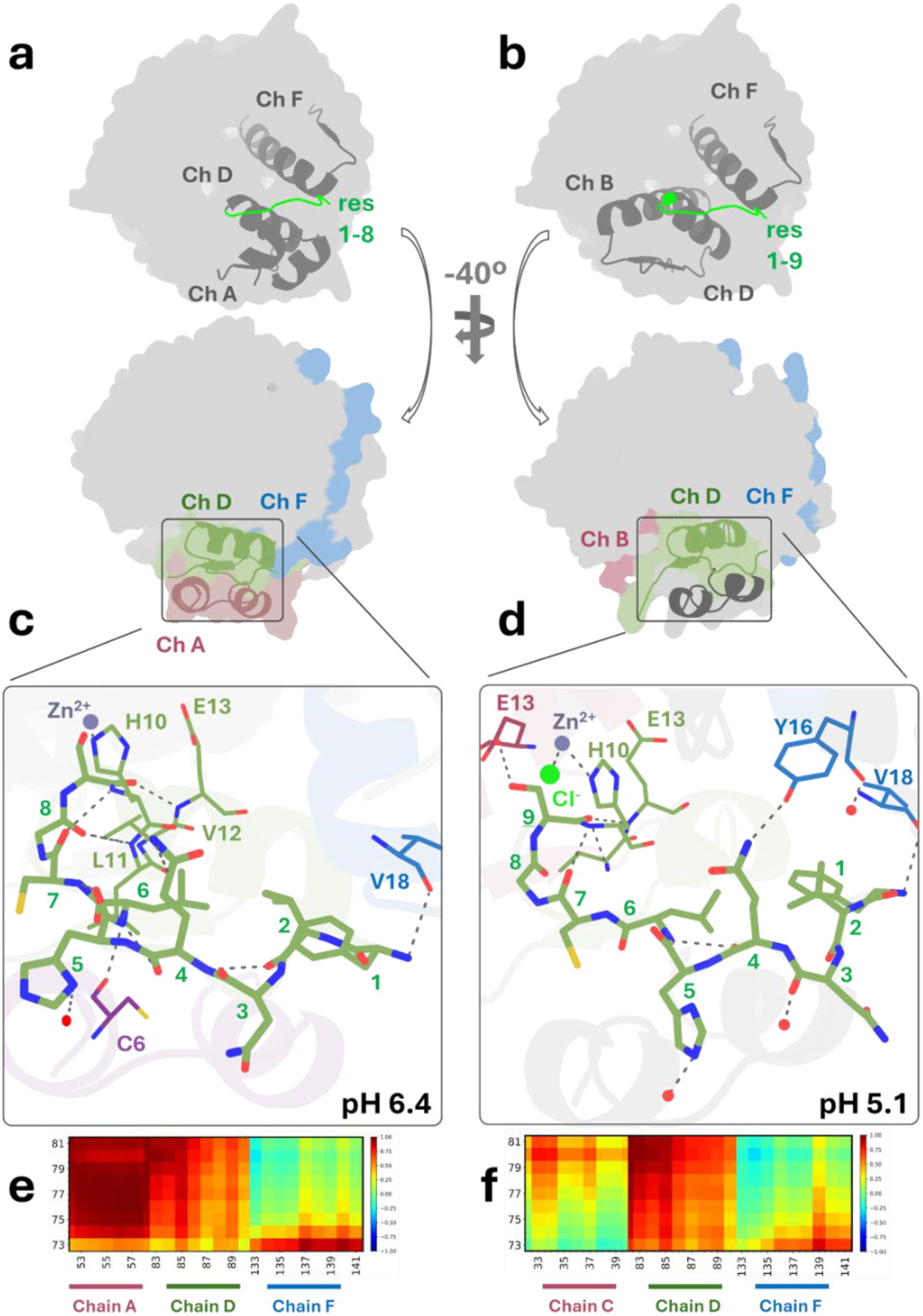
Enhanced cooperativity of the *T*-state despite elevated plasticity and dissociation propensity. The *T*-state monomers at pH 6.4 and 5.1 are characterized by elevated *B*-factors and increased structural plasticity; yet, they exhibit markedly enhanced dynamical cooperativity relative to the *R*-state. a,b, Upon transition to the *T*-conformation, the allosteric B-chain N-terminal segment (residues 1–8 at pH 6.4; residues 1–9 at pH 5.1) undergoes a pronounced reorganization that promotes interactions with neighboring monomers (*eg, chains A and C*), in contrast to the *R*-state. c,d, Close-up views reveal that the allosteric region engages in extensive intra- and inter-monomer contacts at both pH 6.4 and pH 5.1. At pH 6.4, allosteric residues (D1-D8) interact with C6_A_, S9_D_–E13_D_, and V18_F_, whereas at pH 5.1, the allosteric interaction (D1-D9) network is rewired to include E13_C_, H10_D_, E13_D_, Y16_F_, and V18_F_. e,f, GNM cross-correlation heatmaps demonstrate, unlike *R*-subunits, both very strong intra-monomer (*see Fig. S9h and Fig. S11b_i-viii_*) as well as inter-monomer cooperativity originating from the *T*-state allosteric region: (e) coupling is maximal between chains A and D at *T*-status of pH 6.4, while (f) at T-status of pH 5.1 the highest cooperativity is dominated in chain D. Together, these results indicate that, despite increased flexibility, the *T*-state supports an *unpeeling motion* with strong cooperativity driven by intra- and inter-monomer allosteric coupling. GNM cross-correlation heatmaps use a common color scale from −1 (anti-correlated) to +1 (positively correlated), with 0 indicating no correlation

**Figure S13.**
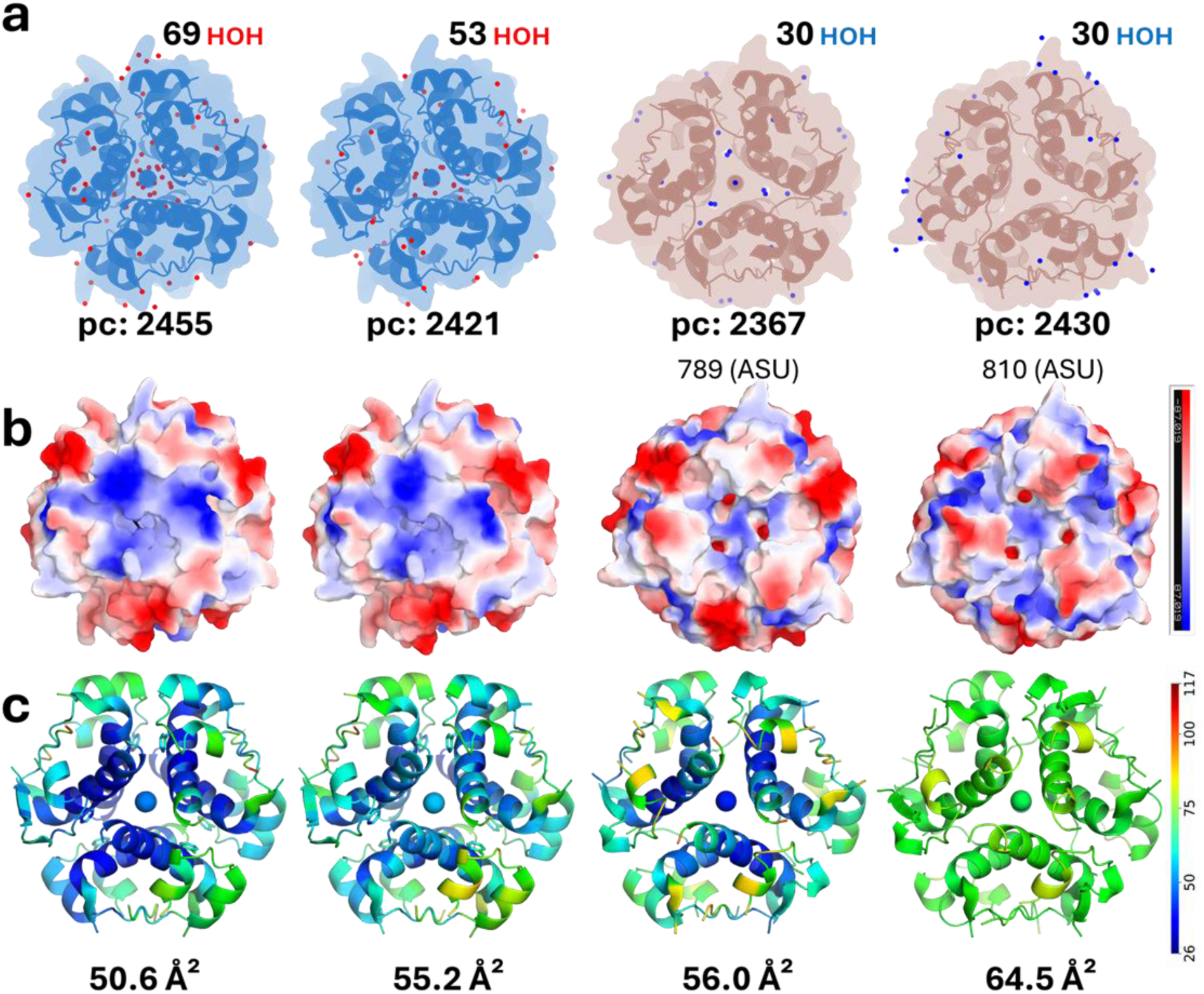
pH-dependent reorganization of hydration, electrostatics, and structural plasticity in the glargine hexamer. a, Comparison of structured water molecules and polar contacts across pH conditions shows a progressiv e reduction in hydration and polar interaction networks with decreasing pH, consistent with a gradual dehydration of the hexamer core. The number and spatial distribution of ordered water molecules (HOH) and polar contact points (pc) are indicated for each condition. b, Electrostatic surface representations calculated by Poisson–Boltzmann analysis reveal a pronounced pH-dependent inversion of electrostatic profile within the hexamer. At high pH (8.4 and 7.3), the hexamer core exhibits a dominant positive electrostatic potential stabilized by Zn²⁺ coordination, whereas acidification (pH 6.4 and 5.1) attenuates this potential, consistent with a protonation-driven electrostatic switch. c, Debye–Waller (*B*-factor) mapping of the hexamer highlights a monotonic increase in structural plasticity toward lower pH, with average *B*-factors rising from ∼50.6 Å² at high pH to ∼64.5 Å² at low pH, indicative of enhanced conformational flexibility accompanying acidification.

**Figure S14.**
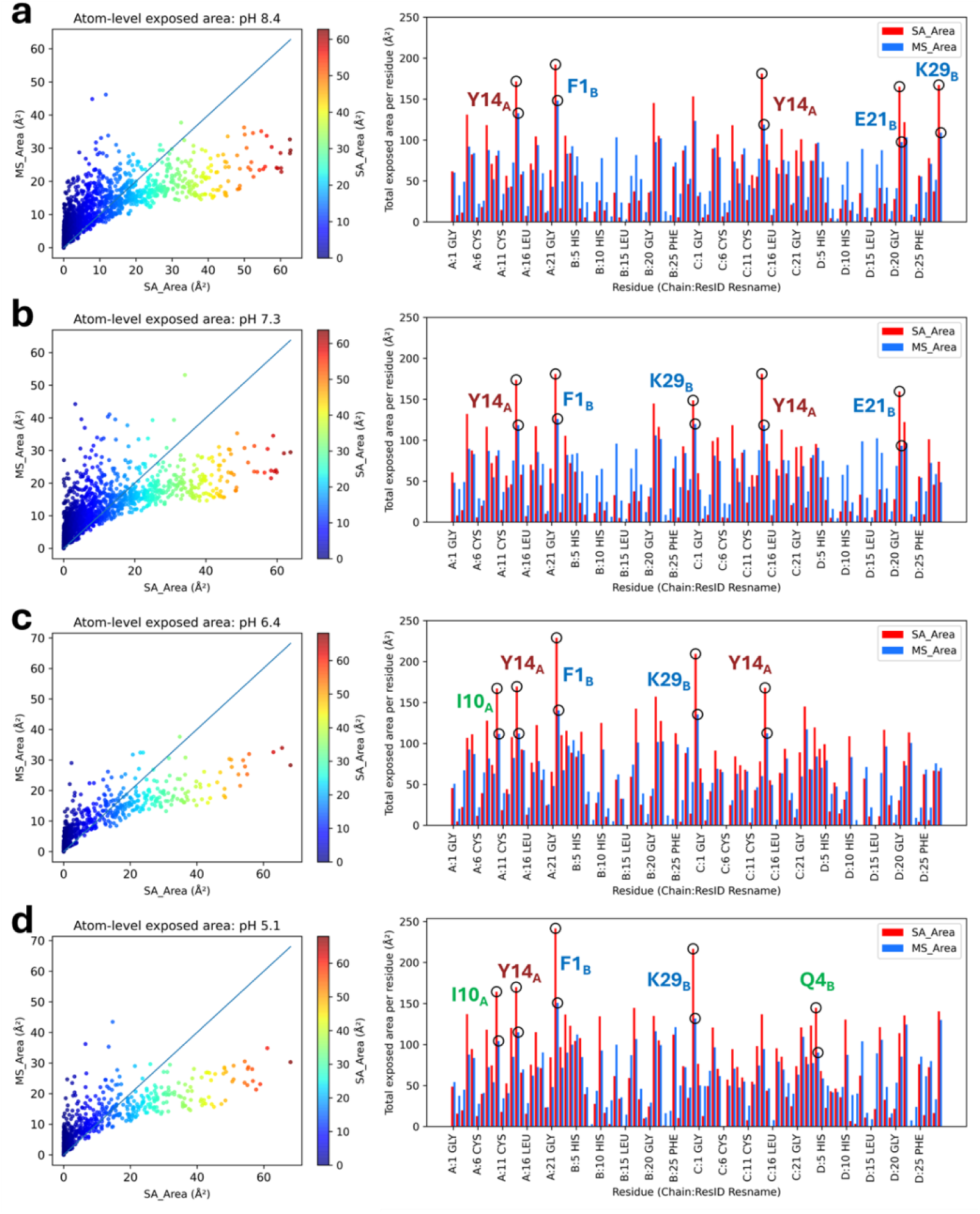
pH-dependent remodeling of surface topology and residue-specific exposure in hexameric insulin glargine. Residue-level analyses of solvent-accessible surface area (SA) and molecular surface area (MS) were computed via *CASTpFold* [61] across the pH gradient (a–d). (Left Panels) Atom-level correlations between SA and MS reveal a largely conserved global topology across all pH values, indicating that the hexameric core retains its overall compactness without widespread unfolding. (Right Panels) However, residue-wise distributions expose a specific reorganization of the surface landscape. At pH 8.4 and 7.3, the surface is dominated by aromatic and charged residues (Y14_A_, F1_B_, E21_B_, K29_B_), consistent with a stable hydrophilic surface. Upon acidification to pH 6.4, the phenol-gatekeeper I10_A_ emerges as a dominant surface residue (c), marking the disruption of the phenol-binding pocket. At pH 5.1, further remodeling is evidenced by the prominent exposure of Q4_B_ (N-terminal anchor) and the concurrent loss of one of the prominent Tyr peaks observed at higher pHs (d). This specific exchange of surface residues—rather than global unfolding—confirms a transition to a ‘molten-like’ topology, with specific hydrophobic patches (Ile, Phe) exposed while others reorient, supporting the reorganization mechanisms detected by fluorescence spectroscopy [72] (*see* Fig. 2).

**Figure S15.**
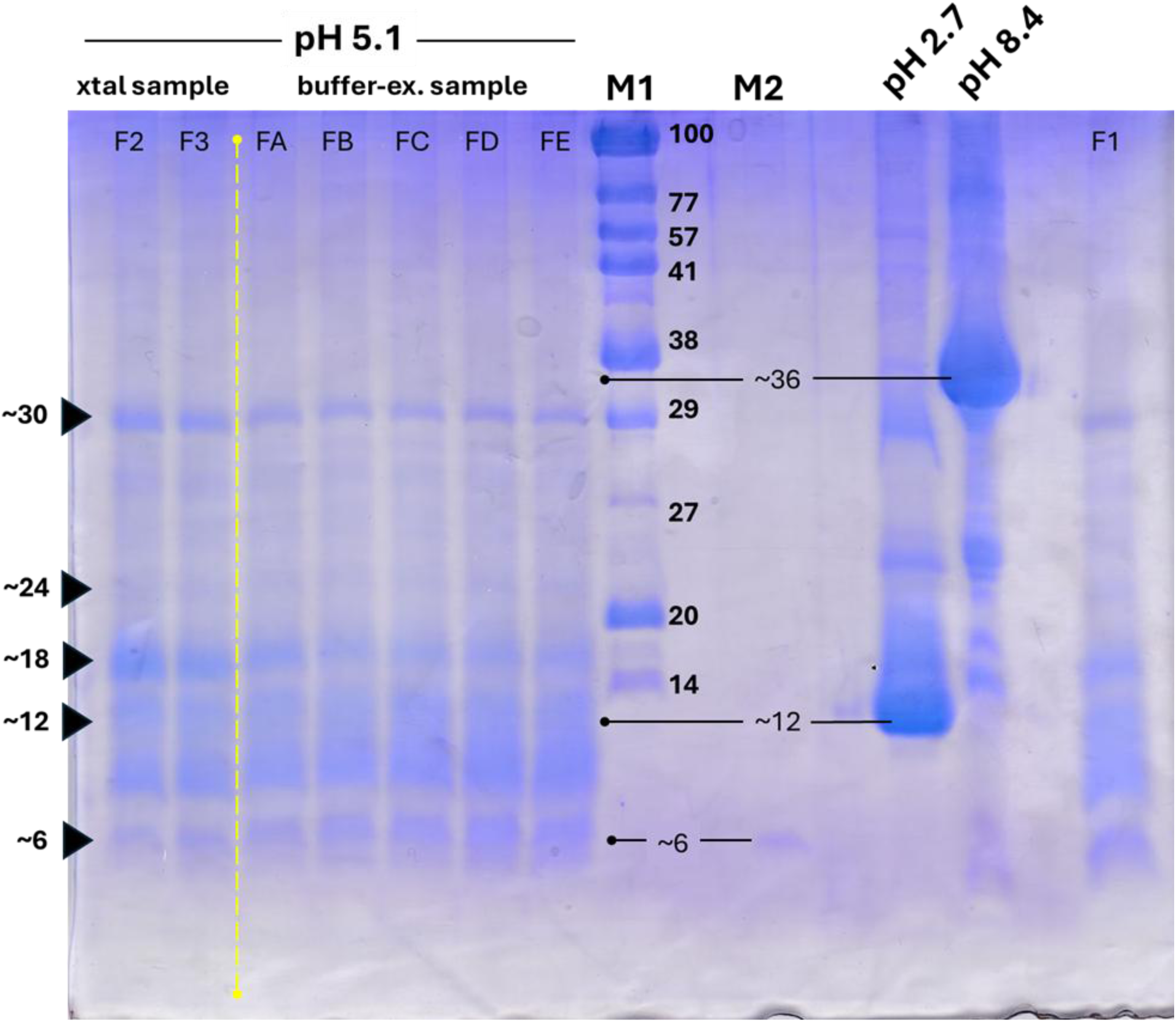
**Uncropped image of the 20% SDS–PAGE analysis**. Lanes F1-F3 correspond to fractions collected from the resolved insulin crystals (referred to as xtal sample), whereas lanes FA-FE correspond to fractions collected from the insulin sample by just exchanging the buffer (referred to as buffer-ex. sample); both methods gave the same peaks, consistent with the broad SEC peak observed at pH 5.1 (*see* Fig. 2a). Prominent bands are observed at approximately 6, 18, and 30 kDa, suggesting the presence of multiple oligomeric states, consistent with a ∼6(x) kDa insulin monomer. Lane M1 shows the molecular weight marker spanning 14–100 kDa (KUTUCARE, Istanbul, Turkiye). Lane M2 corresponds to Lantus (Sanofi) from the pen (pH 4) and was heated at 95 °C for 7 min to induce monomerization, serving as a monomer control. Insulin solutions at pH 2.7 and pH 8.4 display a smeared pattern accompanying the SEC peak.

**Figure S16.**
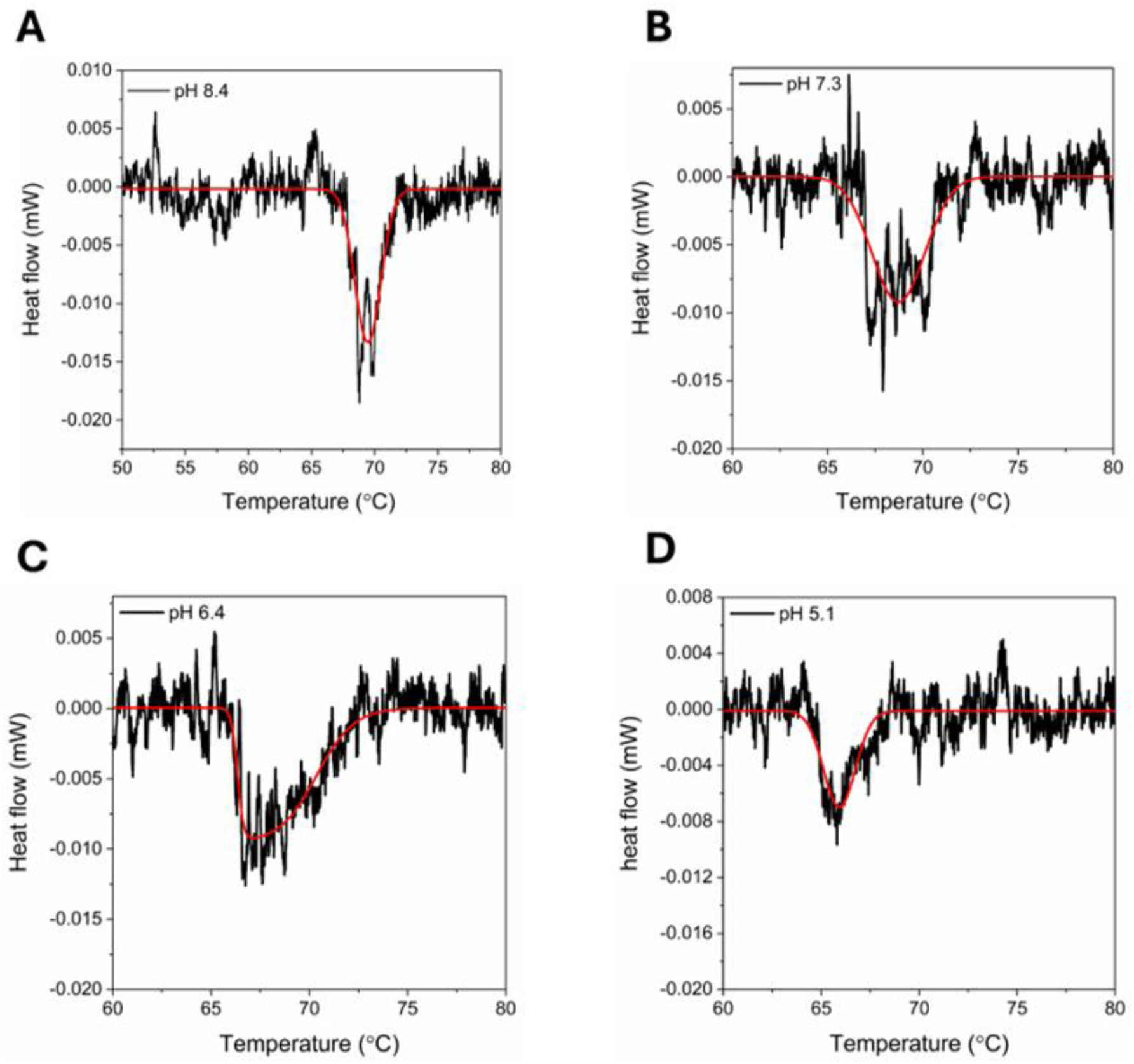
Raw DSC thermograms (a-d) measured under pH 8.4, 7.3, 6.4 and 5.1, respectively. Red line indicates the Gaussian fit to the raw thermal curves.

**Figure S17.**
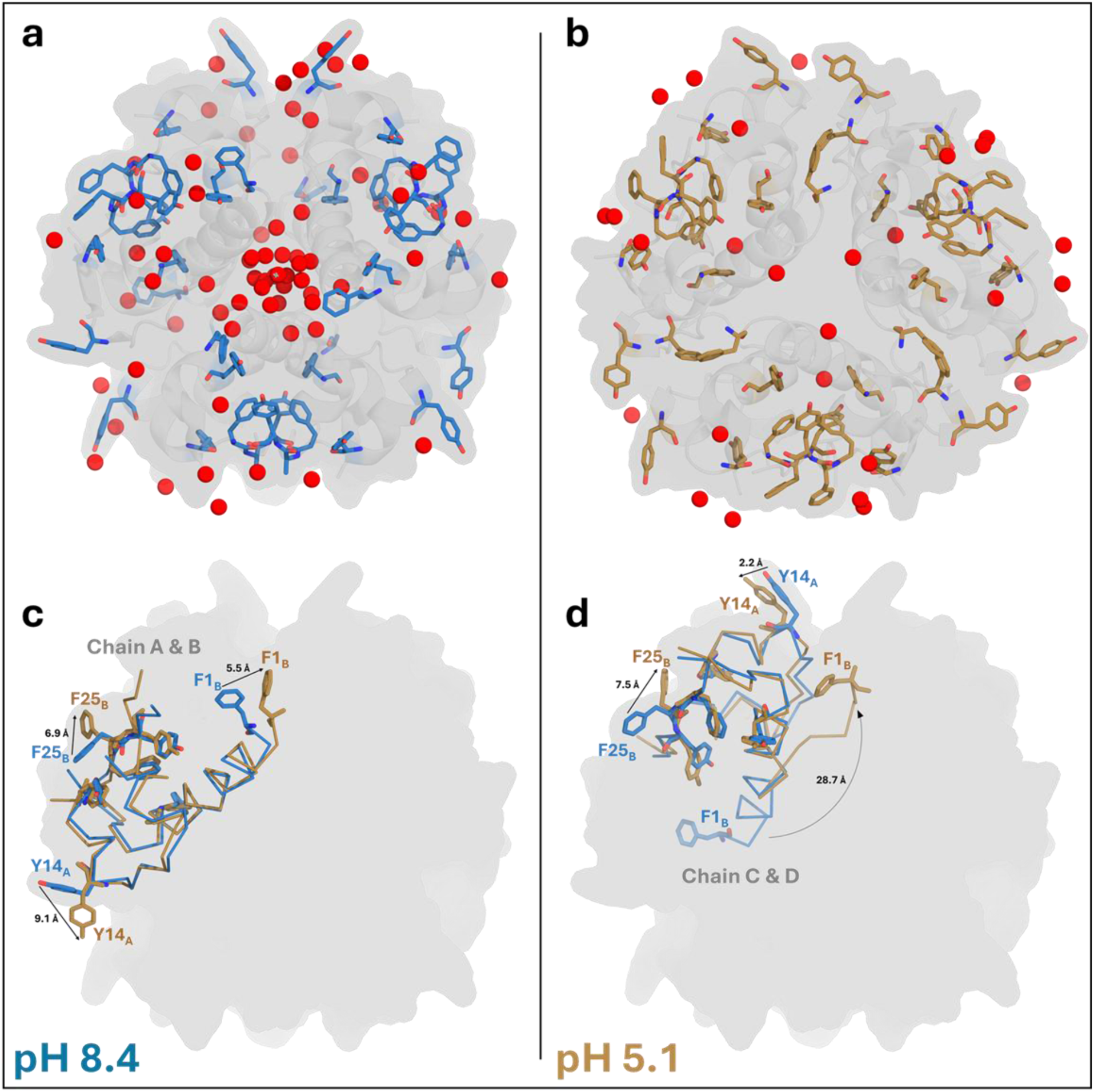
pH-dependent reorganization of aromatic side chains and hydration environment underlying intrinsic fluorescence changes. a,b, Hexamer determined at pH 8.4 (a, blue) and pH 5.1 (b, brown). Red spheres denote structured water molecules. Upon acidification, aromatic side chains undergo pronounced reorientation and redistribution, accompanied by a reduced, reorganized hydration environment. c,d, Structural alignment of the first two monomers from the pH 8.4 (c) and pH 5.1 (d) structures, stressing pH-dependent changes in distances and orientations between selected Tyr and Phe residues (e.g., Y14_A_, F1_B_, F25_B_). At low pHs, altered interaromatic distances and large-scale displacements reflect the repacking of aromatic side chains and the remodeling of the local aromatic microenvironment. This suggests that the hexamer does not simply disintegrate; rather, it undergoes a hydrophobic collapse where aromatic residues are sequestered from solvent (*see* Fig. 2g*; increased fluorescence*), and the loosened subunits move in a highly coordinated, ‘breathing’ motion (*see Fig. S9h; highest cross-correlation*), characteristic of a stable molten-globule intermediate.

**Figure 18.**
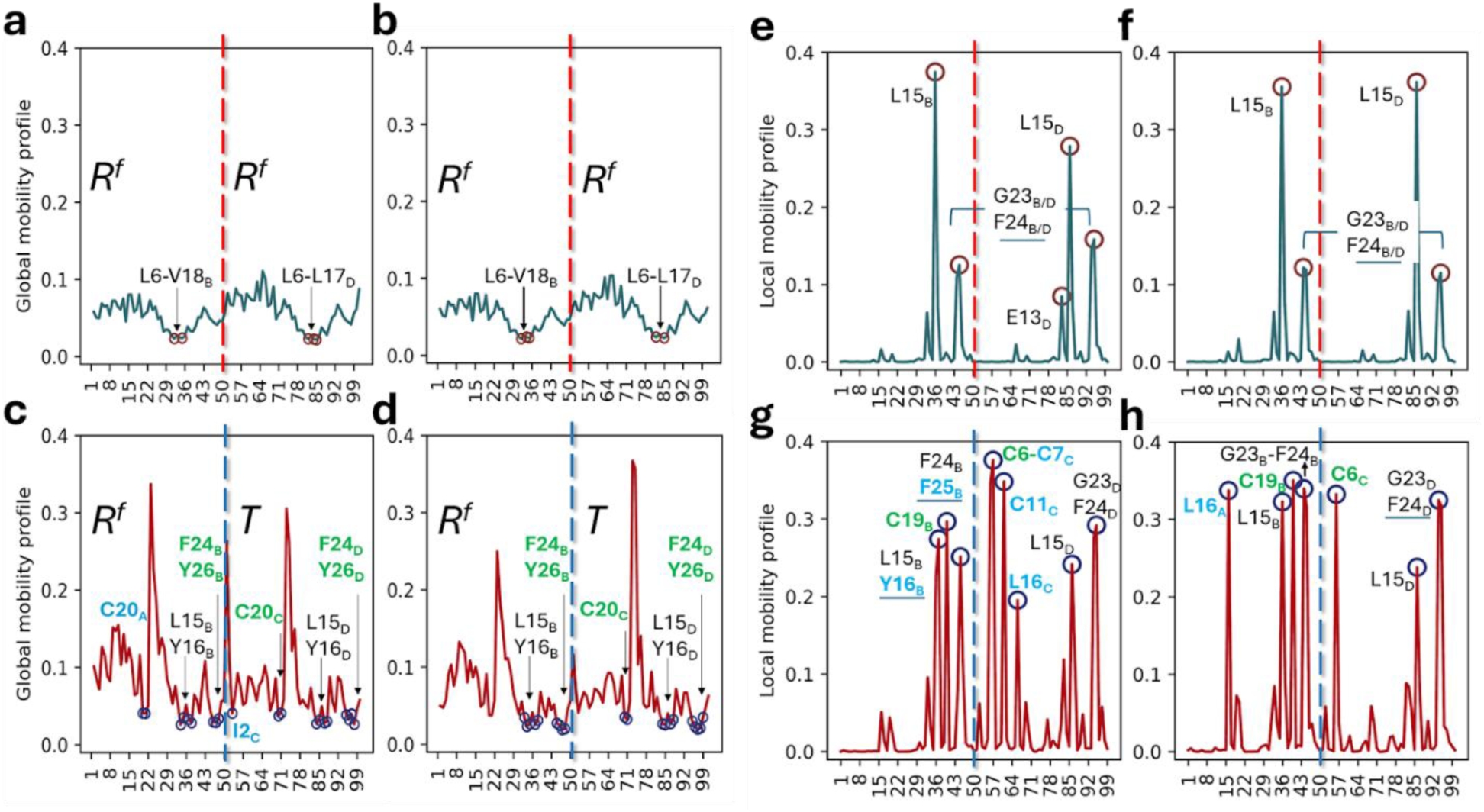
pH-dependent evolution of collective dynamics and local mechanical stiffness in insulin dimers. . Residue mean-square fluctuation (MSF) profiles predicted by the GNM based on (a–d) soft/slow modes with a cumulative covariance contribution > 0.40 and (e–h) high-frequency modes (fastest 10 modes) illustrate how the dynamical control of the first dimers reorganizes along the pH gradient (pH 8.4 to 5.1), transitioning from predominantly rigid-body–like global mechanics at high pH to redistributed, compensatory stabilization at low pH. (a,b) At pH 8.4–7.3, the *R^f^*-state architecture yields a broad, continuous mechanical pivot (dots) spanning the central B/D-chain helix (L6_B/D_–V18_B/D_), consistent with coherent, concerted dimer motion. (c,d) With acidification (pH 6.4–5.1), this continuous hinge is progressively disrupted, consistent with “unpeeling” of the N-terminus; control of global mobility shifts to distinct anchoring regions within the structural core (L15_B/D_–Y16_B/D_) and the C-terminal β-strand (F24_B/D_–Y26_B/D_), together with additional constraints at the A-chain termini (including I2_C_ and C20_A/C_). MSF profiles from the 10 fastest modes highlight kinetically active residues as sharp peaks, indicating local stability centers. (e, f) At high pH, dominant stabilization is localized in the canonical hydrophobic core triad, with prominent peaks at L15_B/D_ and the G23_B/D_–F24_B/D_ segment. (g, h) At lower pH, the loss of helical rigidity recruits new stability centers; peaks broaden and shift to include disulfide - bridge neighbors (C19_B_, C6_C_–C11_C_) and the A-chain core (L16_C_). The rise of the disulfide peaks in the *T*-state (res 50-100 in g, h) at lower pH is not coincidental; it is consistent with the release of phenol, which strongly interacts with CysA6 and CysA11 in the *R*-state (REF). This confirms that upon the loss of ligand-induced helical rigidity, the hexamer recruits these disulfide bridges as essential new stability centers. The progressive diminution of Tyr (Y16_B_) and Phe (F24_B_) from pH 6.4 to pH 5.1 *in-silico* is consistent with *in-solution* fluorescence emission measurements (*see* Fig. 2g) and crystallographic surface topology analyses (*see Fig. S14c-d*). Collectively, these changes support a compensatory stiffening of the cystine-knot/disulfide-stabilized framework, helping maintain dimer integrity as the B-chain termini become more molten-like and dynamically labile.

**Figure S19.**
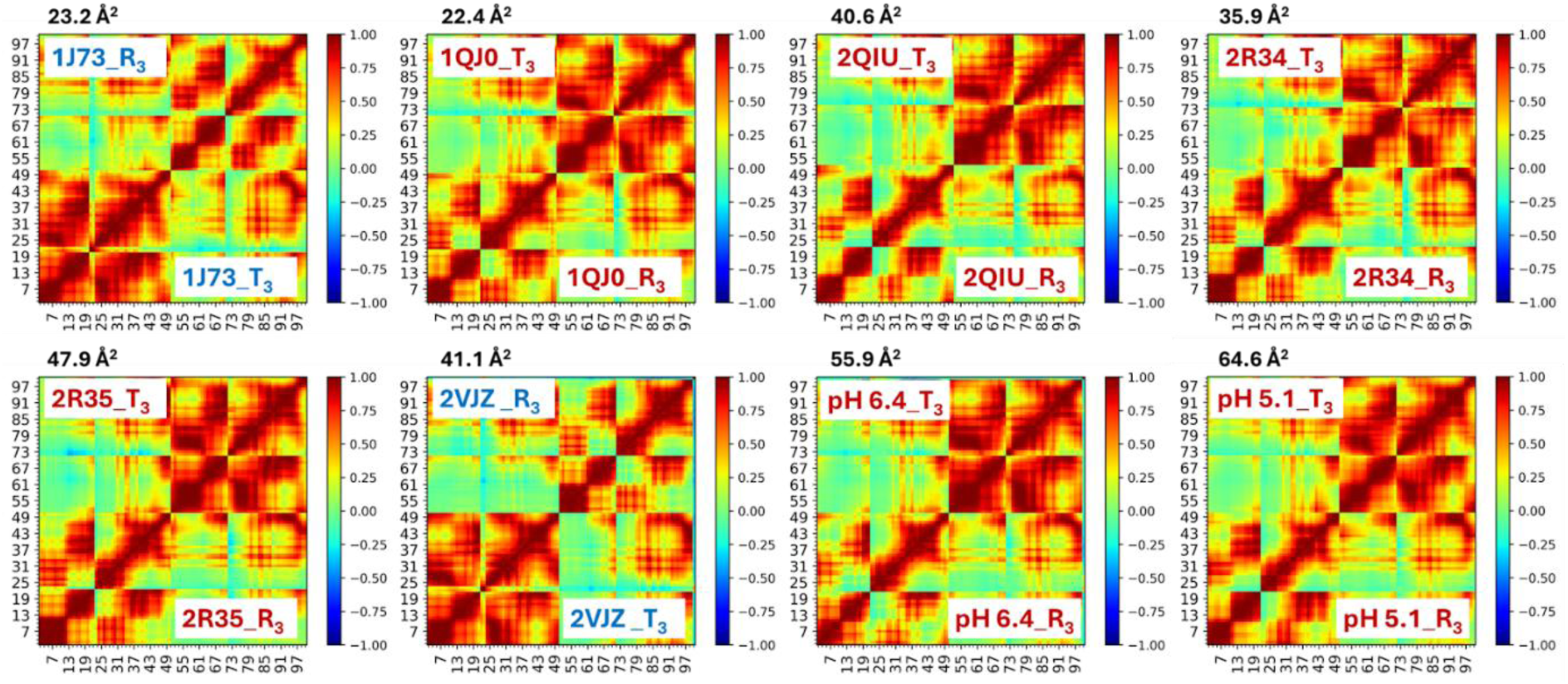
Cross-correlation heatmaps (GNM) comparing the *T₃R^f^₃* subunits of rhombohedral insulin hexamers across ambient SFX and PDB structures. Shown are pairwise cross-correlation maps derived from GNM analysis (cumulative variance of ≥0.40), comparing SFX *T₃R^f^₃* structures (pH 6.4 and pH 5.1) to previously available PDB entries (1J73, 1QJ0, 2QIU, 2R34/35, 2VJZ). Across all datasets, *R₃* subunits (referred to as XXXX_R_3_) consistently exhibit moderate cooperativity with distinct local correlation patterns, whereas *T₃* subunits (referred to as XXXX_T_3_) display stronger, more collective dynamics. This reinforces the view that all *T*-subunits—regardless of temperature or structural origin—exhibit greater global flexibility and collective dynamics than all *R^f^*-subunits, thereby further supporting our concept of a molten-like insulin globule for the pH 6.4 and pH 5.1 SFX structures.

**Figure S20.**
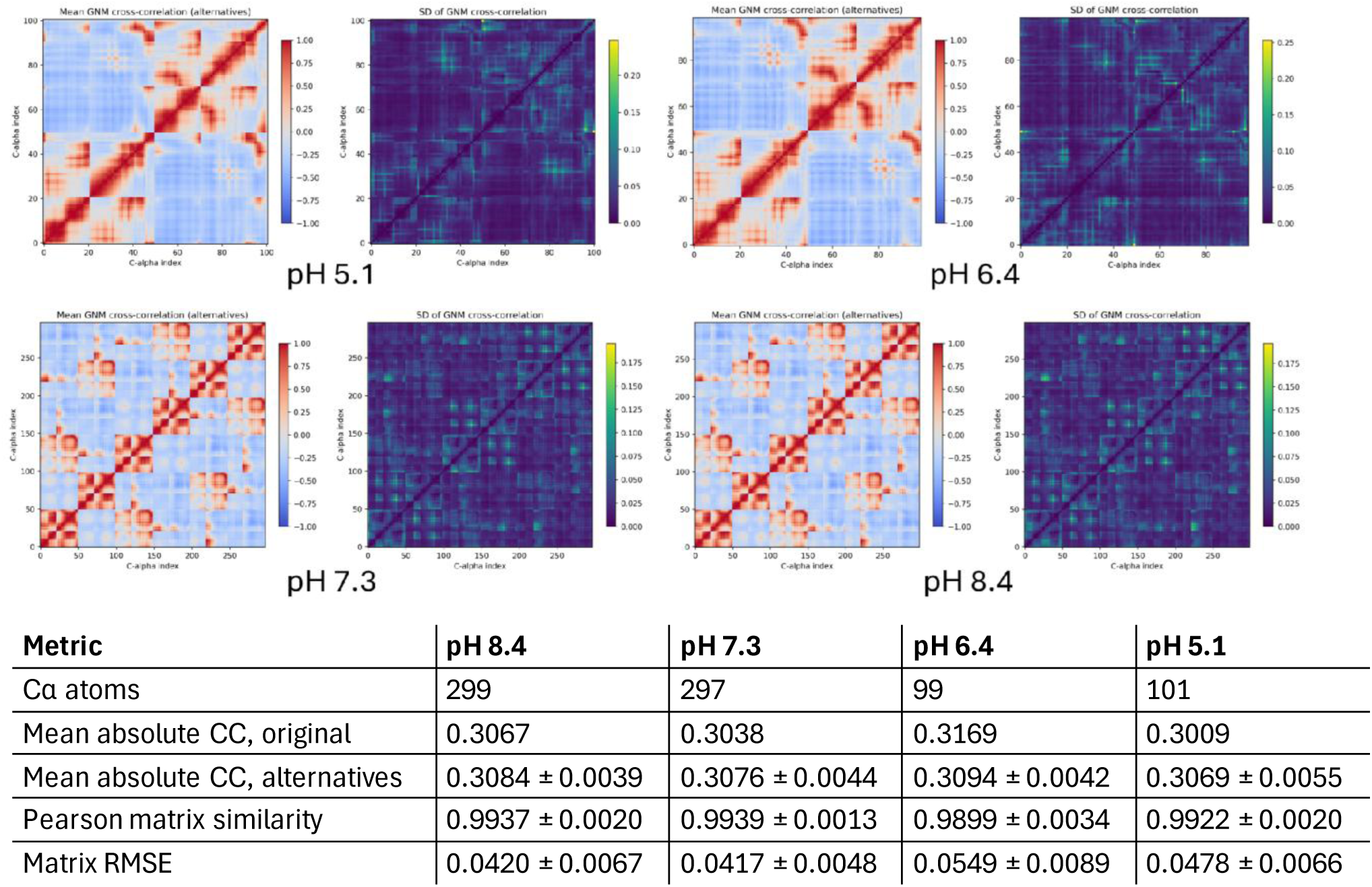
Robustness of GNM-derived cross-correlation patterns to structural perturbations across pH conditions. Mean residue–residue cross-correlation matrices and corresponding standard deviation (SD) maps calculated from 50 alternative structural models at pH 5.1, 6.4, 7.3, and 8.4. Alternative models were generated by applying Gaussian perturbations to Cα coordinates (σ = 0.2 Å). GNM calculations were performed using ProDy with an 8 Å cutoff and the first 11 modes. Mean cross-correlation maps (left panels) illustrate the average residue–residue correlations, whereas SD maps (right panels) indicate their variability across perturbed models. Near-neutral pH structures (8.4 and 7.3) contained 299 and 297 Cα atoms, respectively, whereas acidic structures (6.4 and 5.1) contained 99 and 101 Cα atoms, respectively. The accompanying table summarizes the original and ensemble - averaged absolute cross-correlations, Pearson matrix similarities, and root-mean-square errors (RMSE; mean ± SD). High Pearson similarities (0.9899–0.9939) indicate preservation of the overall correlation patterns despite localized sensitivity to coordinate perturbations.

**Figure S21.**
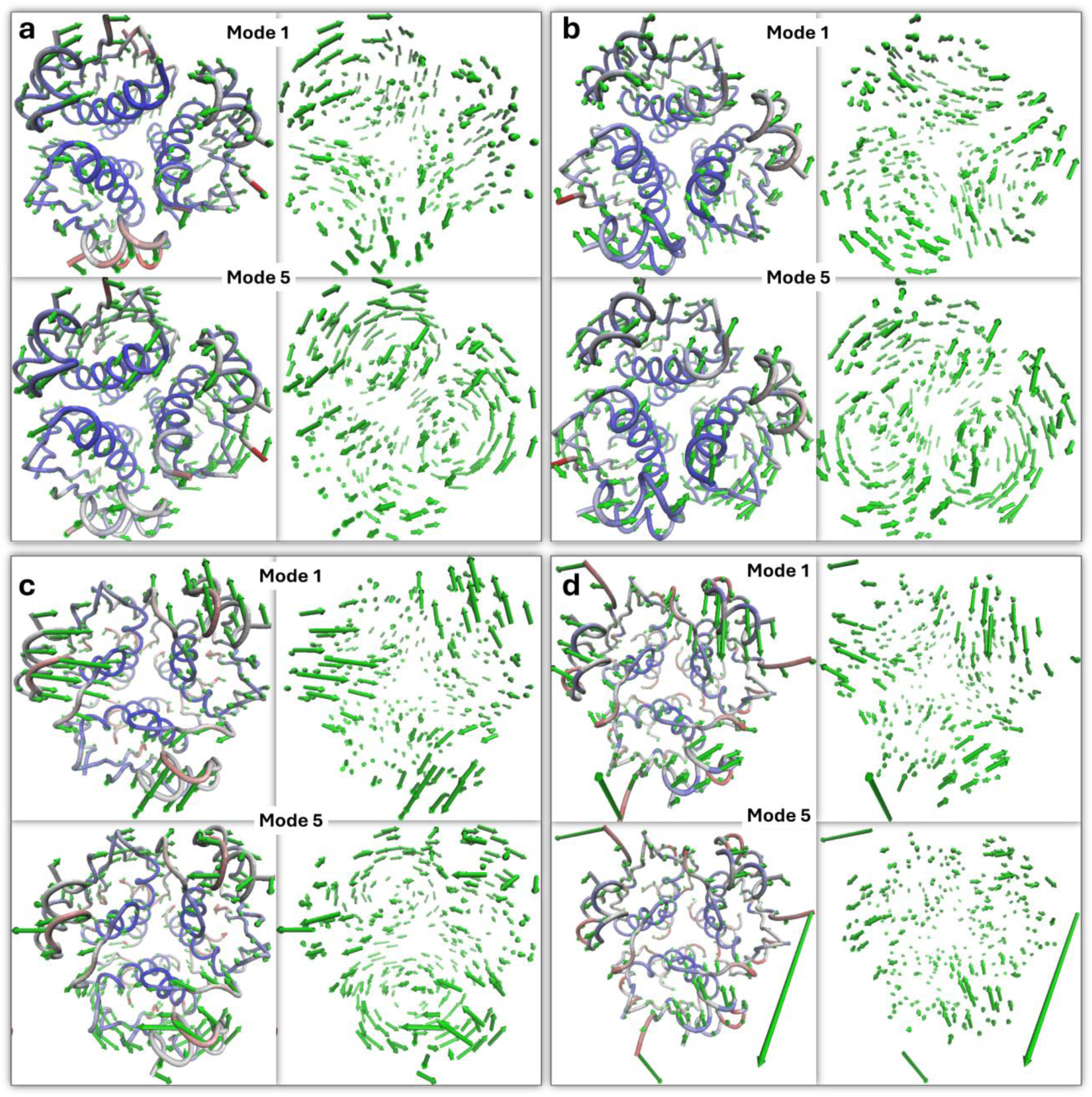
ANM-predicted collective motions of insulin glargine hexamers across different pH conditions. a–d, Representative low-frequency anisotropic network model (ANM) modes calculated from the SFX-derived hexamer structures at pH 8.4, 7.3, 6.4, and 5.1, respectively. For each pH condition, Mode 1 (upper panels) and Mode 5 (lower panels) are shown as structural representations (left) and corresponding displacement-vector fields (right). Green arrows indicate the predicted direction and relative magnitude of Cα displacements, highlighting differences in collective motion patterns across the pH-dependent conformational states.

**Figure S22.**
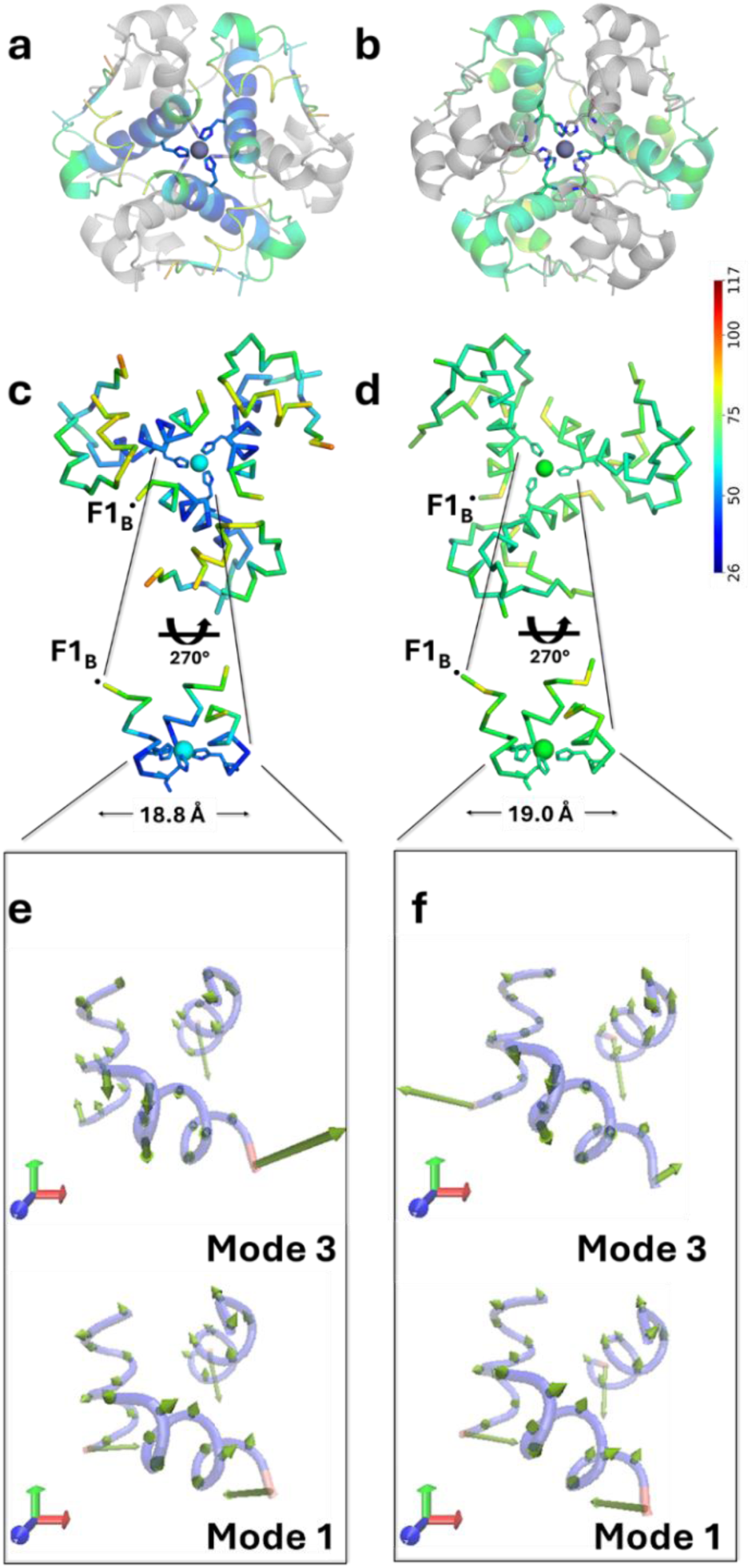
Acid-induced redistribution of allosteric rigidity within the *R_3_* core. (a–b) Trimer cartoons colored by *B*-factor reveal progressively increased structural plasticity upon acidification. (c –d) Ribbon representations of the *R_3_* subunits at pH 6.4 and 5.1. In the *T_3_R_3_* hexamer, the *R_3_* portion is stabilized by Cl⁻ anion and structured water molecules rather than phenol; however, a subtle displacement is observed in the buried allosteric core between pH 6.4 and Ph 5.1 (∼18 Å vs. ∼19 Å), consistent with the increased hexamer plasticity at lower pH. (e–f) ANM projections of the buried allosteric regions (*R* portions) at pH 6.4 and 5.1 along modes 1 and 3, showing a similar motion comparable to that observed under near-neutral pH conditions (pH 8.4 and 7.3).

**Figure S23.**
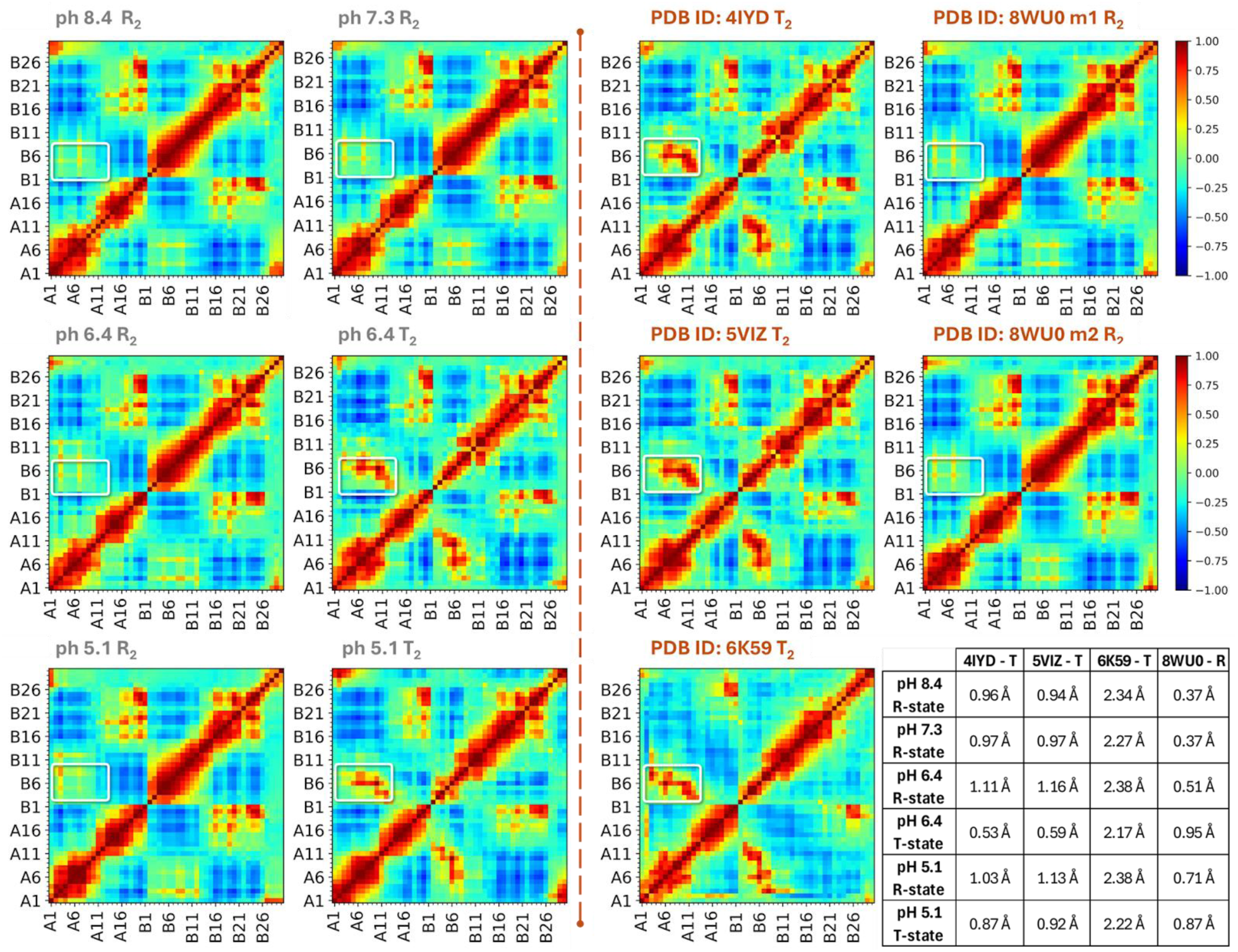
In silico comparison of the SFX-derived glargine dataset with previously reported glargine structures in the PDB. Cross-correlation maps illustrate residue–residue motions driven by GNM soft modes within each subunit of the asymmetric unit (ASU; color scale shown on the right). The axes correspond to residue numbers, primarily residues 1–21 of chain A and 1–29 of chain B. The white square in each map highlights the allosteric region (B1–B8) in *R* and *T* subunits across the analyzed structures. The *R* subunits consistently exhibit weaker collective correlations, whereas the *T* subunits display stronger collective coupling. This pattern supports the pH-dependent reorganization before dissociation observed in our shifted *T_3_R_3_* crystal structures upon acidification and remains broadly consistent even with the dynamicity reported for glargine structures across different lattice types. The lower-right panel illustrates the structural deviation (RMSD) between the R and T subunits across the available glargine models 4IYD, 5VIZ, 6K59, and 8WU0.

**Table S1.** Data collection and indexing statistics for insulin glargine SFX experiments.

| **sample** | **hits** | **hits accepted** | **indexed** | **index hit** |
|---|---|---|---|---|
| **pH 8.4** | **24396** | **24.8%** | **19180** | **78.7%** |
| **pH 7.3** | **17819** | **12.8%** | **11332** | **75.7%** |
| **pH 6.4** | **21980** | **28.0%** | **10880** | **50.6%** |
| **pH 5.1** | **3170** | **2.80%** | **1801** | **53.0%** |

